# Propionic acid-related inhibition during anaerobic digestion: insights into methane production and microbial community adaptation

**DOI:** 10.1101/2025.05.26.656080

**Authors:** Xiaojun Liu, Chloé Soulard, Véronique Jamilloux, André Pauss, Laura André, Thierry Ribeiro, Sabrina Guérin-Rechdaoui, Vincent Rocher, Carlyne Lacroix, Chrystelle Bureau, Cédric Midoux, Olivier Chapleur, Ariane Bize, Céline Roose-Amsaleg

## Abstract

Propionic acid (HPr) accumulation is a major indicator of anaerobic digestion (AD) dysfunction, yet the relative contributions of acidity, undissociated HPr, and propionate ions (Pr⁻) to process inhibition remain poorly understood. We investigated these effects in mesophilic batch AD microcosms fed with municipal sewage sludge, using a comparative design involving HPr, sodium propionate (NaPr), NaCl, and HCl treatments across two series of experiments. While 20 mM HPr caused a 22% reduction in the maximal methane production rate, 81 mM HPr led to complete inhibition, with the initial pH dropping to 5.1. By contrast, 81 mM NaPr reduced methane production rate by only 40%, and 81 mM NaCl caused no inhibition, demonstrating that acidity is the dominant inhibitory factor, with Pr⁻ exerting a secondary concentration-dependent effect. *16S rRNA* gene amplicon sequencing revealed strong, compound-specific shifts in microbial community composition, affecting key functional groups including syntrophs and methanogenic archaea. The proportion of methanogens dropped from 2-3% in control reactors to less than 0.2% under 81 mM HPr, consistent with the observed methane production inhibition. Under HPr81, over 100 ASVs were differentially abundant compared to controls, a pattern largely shared with HCl-treated reactors, further confirming the predominant role of acidity. The number of differentially abundant ASVs was negatively correlated with methane production rates (R² = 0.97), underscoring the link between community reshaping and process impairment. These results provide a unifying framework for propionate inhibition in AD and suggest that microbial community profiling could serve as an early warning tool for process imbalance detection.

## 1. Introduction

Anaerobic digestion (AD) is an environmental biotechnology process for organic waste and effluent treatment, that produces renewable energy (biogas), whose large-scale deployment is a key target of European energy policy, highlighting the need to strongly understand and optimize the AD process.

AD involves four microbial-mediated successive steps: hydrolysis, acidogenesis, acetogenesis and methanogenesis. Acidogenesis converts long-chain fatty acids, amino acids, and carbohydrates into volatile fatty acids (VFAs) like acetic, propionic (HPr), and butyric acids. These VFAs may subsequently be consumed by acetogenic microorganisms producing acetic acid, H_2_ and CO_2_, which are the main precursors for methane synthesis. However, VFAs are also major inhibitors of AD, potentially leading to AD failure, as they can accumulate due to an imbalance between acid production and consumption mediated by acidogenic and acetogenic bacteria (reviewed in (Alavi-Borazjani et al., 2020)).

HPr (pK_a_ = 4.88) content is considered as an indicator predicting the dysfunction of AD since it can accumulate in case of process dysfunction and is also toxic to the microbial activities. During AD, HPr is rapidly dissociated into propionate (Pr^-^) ions upon exposure to AD medium, usually basic. When accumulating in important amounts, HPr, along with other VFAs, can be responsible for the acidification of digesters by decreasing the pH (Sawyer et al., 1954). Undissociated HPr, a form favored by lower extracellular pH (acidified condition), penetrates the intracellular medium and dissociates there. The dissociation causes a decrease in intracellular pH. To remediate this and maintain the proton gradient necessary for ATP formation, the cell actively transports protons to the extracellular medium, thereby reducing the ATP available for its growth and metabolism (Jiang et al., 2018). The intracellular accumulation of dissociated HPr is also toxic to microorganisms, as it increases intercellular ionic strength (Roe Andrew et al., 1998).

The inhibition of AD related to HPr has been studied for a variety of organic waste. Nevertheless, the conclusions are very variable according to the study. Several works observed no inhibition with HPr concentration up to 50 mM (equivalent to 3700 mg L^-1^) during AD of diverse waste under both thermophilic and mesophilic conditions (Qiao et al., 2013; Yang et al., 2015). In contrast, other studies (Demirel & Yenigün, 2002; Wang et al., 2009) reported a decrease in biogas yield at HPr concentrations below 12.2 mM (903 mg L^-1^). Han and colleagues (Han et al., 2020) concluded that reversible inhibition with an increased lag phase could occur at HPr concentrations above 22.7 mM (1,680 mg L^-1^). This variability is likely due to differences in the nature of the substrate and inoculum, as well as other operating conditions. Moreover, HPr (Demirel & Yenigün, 2002; Wang et al., 2009) or sodium propionate (NaPr, a weak base) (Yang et al., 2015) were used, with obvious different effects. This highlights the need for further research to better disentangle the effects of these different parameters through comparative analysis.

Understanding the microbial community response to increased HPr concentrations during AD is moreover of interest to determine the extent, reproducibility and limits of microbial community adaptation, but investigations have been limited to date. During AD, propionate is converted by propionate oxidizers, particularly syntrophic propionate oxidizing bacteria (SPOB) interacting with acetogenic and hydrogenotrophic methanogens (reviewed in (Westerholm et al., 2022)). These two functional groups are likely to be affected by the presence of HPr and are also known to be more sensitive to perturbations compared to hydrolytic and acidogenic bacteria (Sikora et al., 2017). According to previous studies, a shift from acetoclastic to hydrogenotrophic methanogenesis enabled the recovery of a digester inhibited by HPr (Han et al., 2020). By contrast, Li and colleagues suggested the positive role of *Methanotrichaceae*, which comprises exclusively acetoclastic methanogens, during propionate degradation in the presence of high ammonia concentrations (Li et al., 2017). This emphasizes the specificity of the microbial response to given operating parameters. A recent study demonstrated that long-term exposure to extreme propionate concentrations (up to 250 mM, i.e. 24 g L^-1^) in mesophilic anaerobic co-digestion of chicken litter and sodium propionate (NaPr) led to the development of a functionally redundant microbiome capable of sustaining methane production; a shift in archaeal populations at around 62.5 mM (i.e. 6 g L^-1^ of NaPr), in particular from *Methanolinea* to *Methanoculleus*, both hydrogenotrophs, was observed (Ochoa-Bernal et al., 2025).

The present study is specifically focused on municipal sewage sludge (MSS) digestion, which is one of the major waste resources feeding anaerobic digesters (Scarlat et al., 2018). In 2023, 1.1 million tons (Mt) of dry matter (DM) MSS were produced in France and 1.6 in Germany, requiring effective valorization methods (EUROSTAT). To reach a unifying view on inhibition by HPr, the following questions are tackled. How does the presence of HPr affect the physico-chemical dynamics of AD as a function of its concentration? Is the observed inhibition primarily due to the presence of dissociated propionate or to the drop in pH caused by HPr dissociation? Finally, to what extent do microbial communities adapt to the presence of propionate and is this adaptation specific to the form in which propionate is added?

To address these issues, two series of tests in model systems, namely in mesophilic AD batch microcosms, were conducted according to an original experimental design. This study demonstrates that propionate itself exerts an inhibitory effect, albeit less pronounced than that of acidity, and highlights a gradual and substantial adaptation of microbial communities, which is nevertheless insufficient to completely offset the inhibition, and which is highly specific to the added compound.

## 2. Material and methods

### 2.1 Experimental study on propionic acid inhibition

#### 2.1.1 Origin of inocula and substrates

All incubation experiments were conducted in AD microcosms. The substrate was a mixture of primary, biological, and tertiary MSS collected from the Seine-Aval WWTP, which processes 1 500 000 m^3^ per day and serves the Paris conurbation (France). Digested sludge used as inoculum was obtained from one of the mesophilic digesters at the same WWTP. Different batches of both types of sludge were utilized for each test series, consistently sourced from the same sampling points. The total solid content (TS), volatile solid content (VS), and pH of the substrate ranged from 3.3% to 4.7% of fresh mass, 71% to 77% of TS and 6.0 to 6.3, respectively; for the inoculum, these values were 2.3% to 2.6%, 61% to 63% of TS, and 7.4 to 7.6, respectively (Supplementary file 1).

#### 2.1.2 Anaerobic digestion tests and reactor sampling

Two series of Biochemical Methane Potential (BMP) tests were conducted and involved different treatments (Table 1). The incubations were conducted using the AMPTS II (Automatic Methane Potential Test System II, Bioprocess, Lund, Sweden), under batch and mesophilic (37°C) conditions, following the guidelines of (Holliger et al., 2016).

**Table 1.**
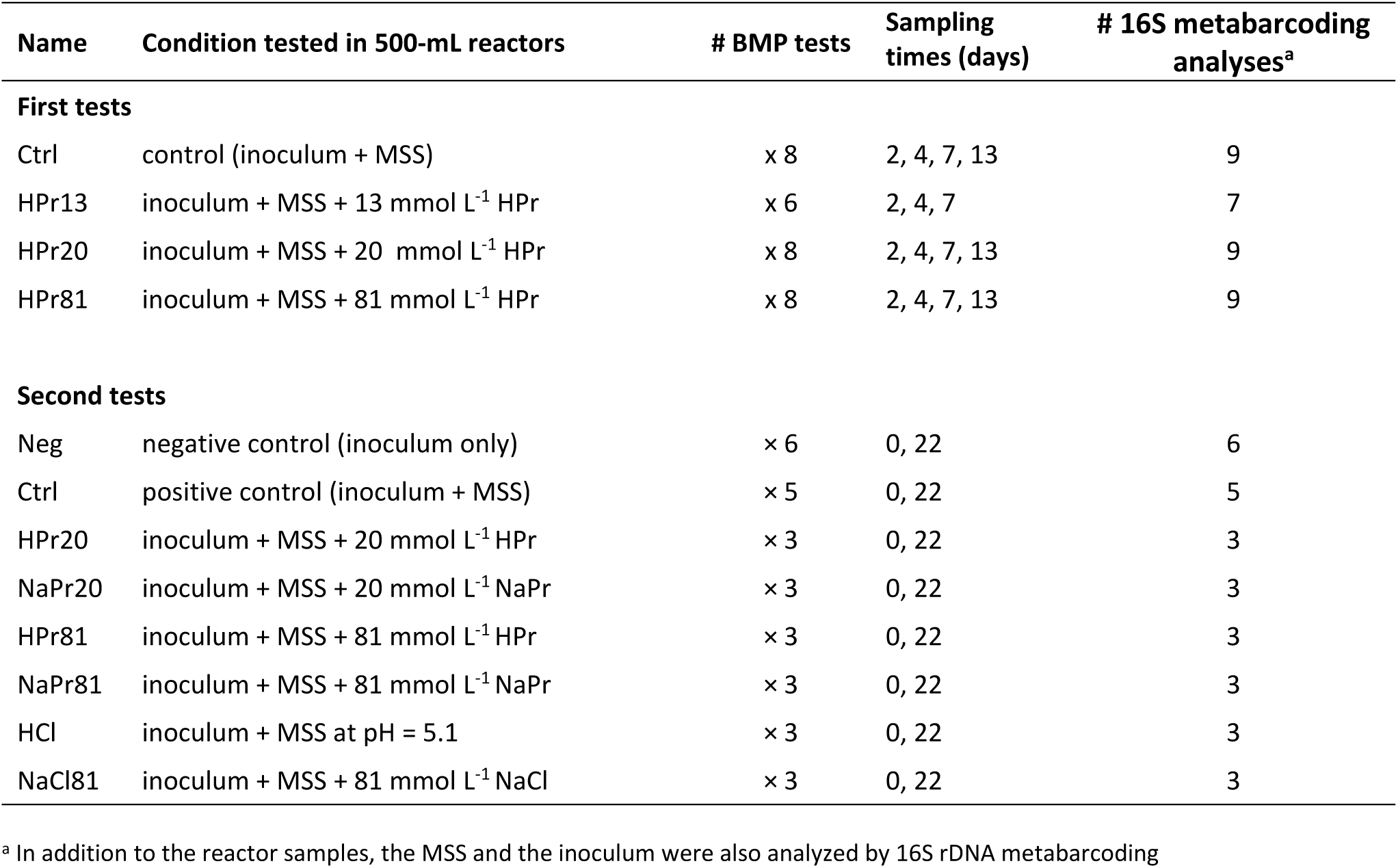
Experimental design for the first and second tests.

| Name | Condition tested in 500-mL reactors | # BMP tests | Sampling times (days) | # 16S metabarcoding analyses<sup>a</sup> |
| --- | --- | --- | --- | --- |
| **First tests** |  |  |  |  |
| Ctrl | control (inoculum + MSS) | x 8 | 2, 4, 7, 13 | 9 |
| HPr13 | inoculum + MSS + 13 mmol L<sup>-1</sup> HPr | x 6 | 2, 4, 7 | 7 |
| HPr20 | inoculum + MSS + 20 mmol L<sup>-1</sup> HPr | x 8 | 2, 4, 7, 13 | 9 |
| HPr81 | inoculum + MSS + 81 mmol L<sup>-1</sup> HPr | x 8 | 2, 4, 7, 13 | 9 |
| **Second tests** |  |  |  |  |
| Neg | negative control (inoculum only) | x 6 | 0, 22 | 6 |
| Ctrl | positive control (inoculum + MSS) | x 5 | 0, 22 | 5 |
| HPr20 | inoculum + MSS + 20 mmol L<sup>-1</sup> HPr | x 3 | 0, 22 | 3 |
| NaPr20 | inoculum + MSS + 20 mmol L<sup>-1</sup> NaPr | x 3 | 0, 22 | 3 |
| HPr81 | inoculum + MSS + 81 mmol L<sup>-1</sup> HPr | x 3 | 0, 22 | 3 |
| NaPr81 | inoculum + MSS + 81 mmol L<sup>-1</sup> NaPr | x 3 | 0, 22 | 3 |
| HCl | inoculum + MSS at pH = 5.1 | x 3 | 0, 22 | 3 |
| NaCl81 | inoculum + MSS + 81 mmol L<sup>-1</sup> NaCl | x 3 | 0, 22 | 3 |

<sup>a</sup> In addition to the reactor samples, the MSS and the inoculum were also analyzed by 16S rDNA metabarcoding

In the first tests, the reactors contained 100 mL substrate (fresh MSS), 100 mL inoculum (digested MSS), and 200 mL deionized water. HPr was added at 0, 13, 20, or 81 mM. To analyze microbial communities and chemical parameters during the course of AD, two reactors from each treatment were sacrificed on days 2, 4, 7, and 13, to collect their content, except for treatment HPr13 that was not analyzed on day 13.

For the second tests, the operation lasted 22 days. Except for the controls without substrate (Neg, 6 replicates) that contained the inoculum (200 mL) and deionized water (200 mL), all other reactors were filled with both inoculum (200 mL) and substrate (200 mL). Either no additional compound was added (Ctrl), or one among HPr, NaPr, NaCl and HCl (Table 1). Regarding HCl, a 10% HCl solution was added dropwise to adjust the initial pH to 5.1, which was the lowest initial pH observed experimentally, under HPr81 (Supplementary File 2, Figure S1). The liquid phase was sampled for microbial community and chemical analysis exclusively before incubation and at the final time point.

For both tests and at each sampling, 1.7 mL aliquots of the liquid phase were centrifuged at 10,000 g for 5 minutes. The pellets and supernatants were stored separately at -20°C until subsequent DNA extraction and total VFA analysis (Supplementary File 3), respectively. The pH values were determined on the fresh liquid phase samples, immediately after collection, using a pH meter (Mettler Toledo, Switzerland).

#### 2.1.3 Curve modeling by modified Gompertz equation

AMPTS recorded daily methane production for each reactor. The data were processed using Office Excel 2019 (Microsoft^®^, Redmond, US). The modified Gompertz equation (**Equation 1**) was used to estimate the kinetic parameters related to methane production for each treatment, namely the biomethane potential and maximum production rate, as well as the lag time. This thereby enabled the quantification and comparison of the possible inhibition among different conditions. Non-linear regression for kinetic modeling was performed using Scilab software (Dassault Systèmes, Vélizy-Villacoublay, France). The model’s goodness-of-fit was evaluated based on adjusted R^2^, sum of squared errors, root mean squared errors and mean absolute percentage error.

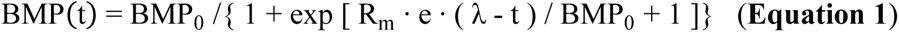

where BMP(t) is the cumulative methane production at time t (NL_CH4·_kg_VS_^−1^ at 0°C, 1 atm), BMP_0_ is the maximum methane yield of the substrate tested (NL_CH4·_kg_VS_^−1^), R_m_ is the maximum methane production rate (NL_CH4·_kg_VS_^−1^.d^-1^), λ is the lag time (d), t is the incubation duration (d) and e ≈ 2.7182.

ANOVA followed by a post hoc Tukey test were performed using R Studio (R Studio Inc., Massachusetts, USA) to assess significant differences between experimental results.

### 2.2 Microbial community analysis

#### 2.2.1 DNA extraction

DNA extraction was conducted from the frozen pellets using the kit Nucleobond^®^ RNA Soil with the complementary DNA Set, following the manufacturer’s instructions (Macherey-Nagel, Düren, Germany). Mechanical cell lysis was performed with TissueLyser II (Qiagen, Hilden, Germany). DNA concentrations were measured using Quantifluo with a SAFAS spectrophotometer, employing the fluorescent dye Picogreen® (ThermoFisher, Waltham, US).

#### 2.2.2 *16S rRNA* gene amplicon sequencing

Sequencing of *16S rDNA* amplicons was performed using an Ion torrent PGM instrument (Life Technologies, Carlsbad, US) after PCR amplification of the V4-V5 hypervariable regions with the fusion primers 515F (5’-Ion A adapter - Barcode – GTGYCAGCMGCCGCGGTA-3’) and 928R (5’- Ion trP1 adapter – ACTYAAAKGAATTGRCGGGG-3’), as previously described (Poirier et al., 2016). A total of 36 samples were processed for the first tests, and 29 samples for the second tests. Raw reads were submitted to the ENA under project accession PRJEB87704.

#### 2.2.3 Bioinformatic processing

Raw amplicon sequencing data were pre-processed with deepomics16S pipeline (https://forge.inrae.fr/deepomics/deepomics16S v14.10.2022) on the servers of the INRAE MIGALE bioinformatics platform. This pipeline employs DADA2 (Callahan et al., 2016), for the generation of Amplicon Sequencing Variants (ASVs) and FROGS (Bernard et al., 2021) for the taxonomic affiliation of ASVs, relying on Silva database (silva_138.1_16S_pintail100). Only the V4 region was considered for samples of the second tests due to a lower sequence quality, using primer sequences 5’-GTGYCAGCMGCCGCGGTA and 5’-GGACTACHVGGGTWTCTAAT for trimming.

Each sample retained more than 15,000 sequences after bioinformatics processing (Supplementary File 2, Figure S2). The raw sequences, biom files and physicochemical data were deposited in a FAIR mode in DeepOmics information system (https://deepomics-info.hub.inrae.fr/).

#### 2.2.4 Biostatistics analysis

The pre-processed sequencing data were analyzed using R version 4.3.2 (Team, 2021) in RStudio (2024.04.2+ 764) environment. Several specific packages were utilized, such as ComDim 0.9.51 (https://github.com/f-puig/R.ComDim) for Common Component Analysis (CCA), DESeq2 1.42.0 (Love et al., 2014) for differential abundance analysis and phyloseq 1.46.0 (McMurdie & Holmes, 2013) for barplot representation. The taxonomic tree was built with GraPhlAn (https://github.com/biobakery/graphlan).

For the first tests, 157 Amplicon Sequence Variants (ASVs) were selected, by removing the ASVs with less than 400 reads in the dataset. For the second tests, ASVs representing less than 0.5% of the reads in the dataset were removed, resulting in 169 ASVs for analysis. However, for the differential analysis, data from the second tests were filtered less strictly, removing ASVs representing less than 400 reads, resulting in a total of 276 ASVs.

## 3. Results

### 3.1 Inhibition of methane production increases with propionic acid (HPr) concentration

To decipher AD inhibition by HPr, four distinct concentrations of HPr: 0, 13, 20 and 81 mM, were added to batch AD microcosms (first tests of this study, Table 1), aiming to obtain contrasted inhibition levels. Based on cumulative methane production, inhibition increased gradually with HPr concentration, reaching complete inhibition under HPr81 (Figure 1A). Additionally, the pH values overall decreased with increasing HPr concentration, which can be explained by both the dissociation of the added HPr and the accumulation of various volatile fatty acids due to acidic inhibition of the process (Supplementary File 2, Figure S3). Indeed, under HPr81, the pH values remained at 5 or 5.1 throughout the entire incubation period (Supplementary File 2, Figure S1). Such pH values are known to inhibit the AD process, especially methanogenesis (Clark & Speece, 1971).

**Figure 1.**
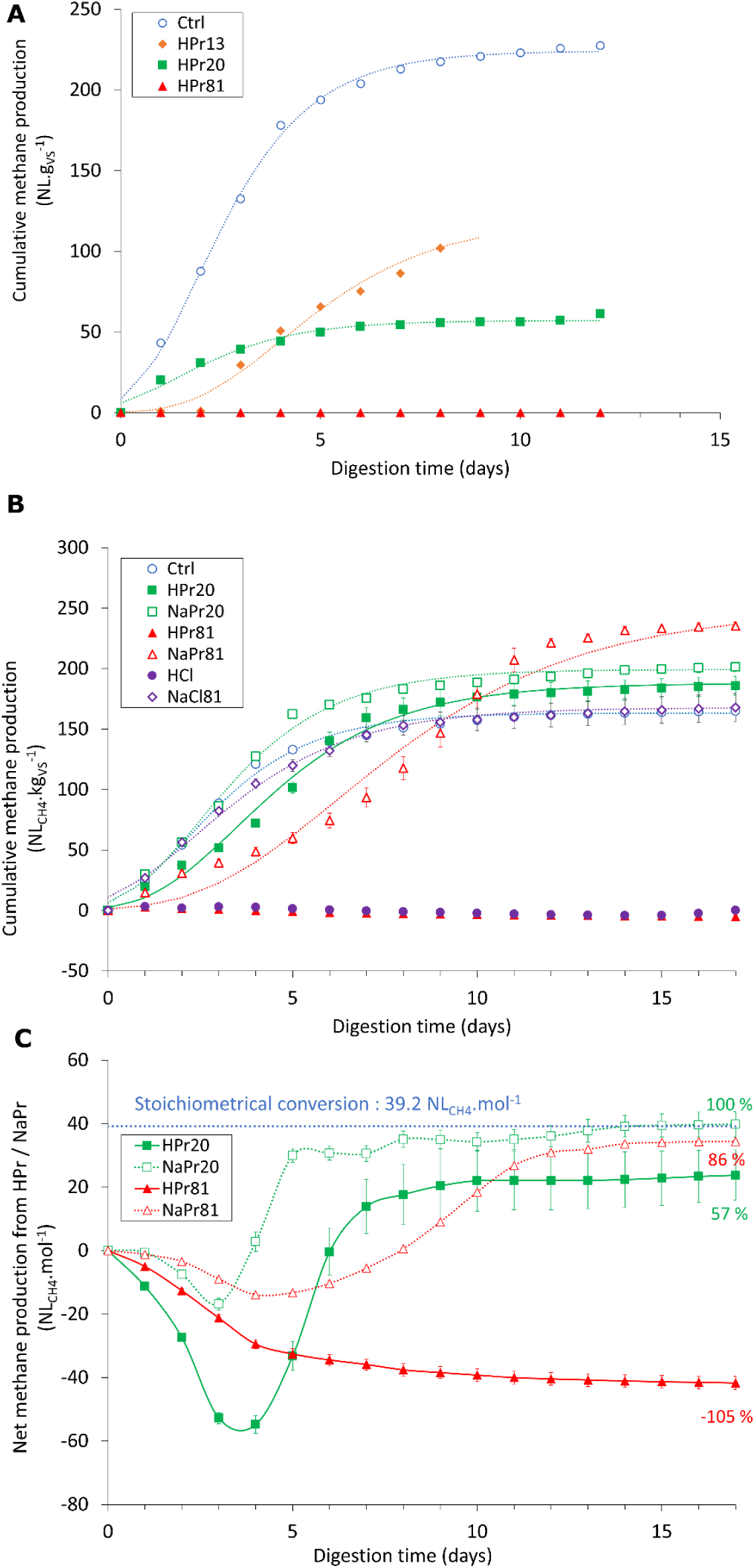
Methane production dynamics during the AD incubations. A. During the first tests. B. During the second tests. C. Net methane production over time during the second tests, obtained by subtracting methane production under Ctrl to the methane production under the considered conditions The curves are the averages of the replicates, the dotted lines are the results of Gompertz identifications (Equation 1).

### 3.2 Methane production is more sensitive to acidic pH and propionic acid (HPr) than to propionate ion (Pr^-^)

The negative effect on methane production following HPr addition may be attributed to the propionic acid itself, its ionized form (propionate, Pr^−^) and/or the low pH condition caused by HPr dissociation, generating protons (H^+^). To investigate different possible effects, distinct incubation conditions were established in second AD tests, by varying the nature of the counterions or targeting specific pH values (Materials and methods, Table 1). Pairwise comparisons between the treatments enabled investigation of the specific effects of certain ions. Comparison of HPr20 and NaPr20 on the one hand, HPr81 and NaPr81 on the other, enabled to assess the contribution of acidity to the observed effects. In addition, NaCl81 treatment aimed to verify the absence of inhibition by sodium ions. Finally, HCl was added to obtain an initial pH value of 5.1, the same as that measured under HPr81, to assess the effect of a pH decrease that was not caused by the addition of HPr. Besides, treatments HPr20 and HPr81 also enabled comparison with the first tests.

The cumulative methane production of MSS under the different conditions was calculated by subtracting the mean gas production obtained under Neg from the raw methane productions (Figure 1B). The negative controls yielded 56 ± 2 NL_CH4_ kg_VS_^-1^, which meets the acceptance criteria for BMP validity (Holliger et al., 2016). Ctrl yielded 250 NL_CH4_.kg_VS_^-1^, with no significant difference compared to production under NaCl81 (253 NL_CH4_.kg_VS_^-1^). This indicates that the concentrations of Na^+^ and Cl^−^ ions tested ([Na^+^] = [Cl^−^] = 81 mM) did not inhibit the AD process. By contrast, under HPr81 or HCl, both leading to an acidic pH of 5.1, no methane was produced.

The addition of propionate at lower levels, either in its HPr or NaPr form, contributed to a delay in methane production, as indicated by the longer lag phases (λ) compared to Ctrl (Table 2). However, it ultimately improved the methane yield (Table 2), which increased up to 348 ± 5 NL_CH4_.kg_VS_^-1^ for NaPr81. With the addition of HPr or NaPr, methane was produced not only through MSS degradation, but also by the consumption of HPr or Pr^−^, which acted as additional substrates for the microorganisms.

**Table 2.** Kinetic parameters of the modified Gompertz equation and fitting goodness for the second tests.

| Treatment assessed | BMP<sub>0</sub><br>(NL CH<sub>4</sub>·kg VS<sup>-1</sup>) | λ<br>(day) | R<sub>m</sub><br>(NL CH<sub>4</sub>·kg VS<sup>-1</sup>·d<sup>-1</sup>) | MAPE*<br>(%) | R<sup>2</sup><br>(-) |
| --- | --- | --- | --- | --- | --- |
| Ctrl | 250 ± 2 | 0.35 ± 0.08 | 56.1 ± 3.9 | 2.16 ± 0.23 | 0.995 ± 0.002 |
| HPr20 | 280 ± 14 | 1.13 ± 0.07 | 43.8 ± 2.9 | 8.70 ± 0.66 | 0.991 ± 0.001 |
| NaPr20 | 291 ± 5 | 0.55 ± 0.01 | 57.8 ± 0.8 | 3.59 ± 0.14 | 0.995 ± 0.000 |
| HPr81 | NA** | NA | NA | NA | NA |
| NaPr81 | 348 ± 5 | 2.45 ± 0.22 | 33.7 ± 1.2 | 15.2 ± 1.0 | 0.981 ± 0.001 |
| HCl | NA | NA | NA | NA | NA |
| NaCl81 | 253 ± 16 | 0.13 ± 0.03 | 43.4 ± 1.0 | 1.69 ± 0.14 | 0.996 ± 0.001 |

\* Mean absolute percentage error of modeling
\*\* Results are not available (NA) because the Gompertz equation was not suitable for the shape of curve
BMP<sub>0</sub> is the maximum methane yield of the substrate tested, λ is the lag time and R<sub>m</sub> is the maximum methane production rate

The methane production under Ctrl was subtracted from that of the other treatments and compared to the stoichiometric production of methane from propionate and propionic acid, which is 39.2 NL_CH4_ mol^-1^ according to **Equation 2**:

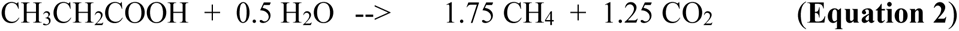

In Figure 1C, the comparison of NaPr and HPr treatments at the same molar concentrations highlighted the significant effect of pH and/or the undissociated form of HPr, compared to that of Pr^-^ ion. Methane was produced even for high Pr- concentration, despite a transient inhibition during the initial hours of the tests. The production reached almost 86% of the maximal methane production under NaPr81.

Overall, although the effect of HPr or the counterion H+ appears to be largely dominant in the inhibition phenomenon, a decrease in methane production rate was observed in the presence of high NaPr concentrations, highlighting the inhibitory effect of Pr- itself.

### 3.3 Significant adaptation of microbial communities to propionic acid (HPr) occurs during ecological successions, even with the smallest addition of HPr

Since the effects on methane production are generally mediated by the response of the catalytic microbial communities to the operating conditions, the adaptive dynamics of microbial community composition was subsequently analyzed, through *16S rDNA* metabarcoding.

Among the 16 bacterial phyla represented in the first tests, the six most abundant at day 13 were Bacteroidota, Spirochaetota Pseudomonadata, Bacillota, Chloroflexota, and Candidatus Cloacimonadota (Supplementary File 2, Figure S4), all commonly identified in anaerobic digesters (Calusinska et al., 2018; Jiang et al., 2021).

Archaea consistently represented less than 4% of the reads in the samples and predominantly belonged to *Methanosarcina* and *Methanolinea* (Figure 2A). The first genus includes metabolically-versatile methanogens encoding three methanogenesis pathways (hydrogenotrophic, acetoclastic and methylotrophic) (Liu & Whitman, 2008), while the second corresponds to strictly hydrogenotrophic methanogens (Sakai et al., 2012).

**Figure 2:**
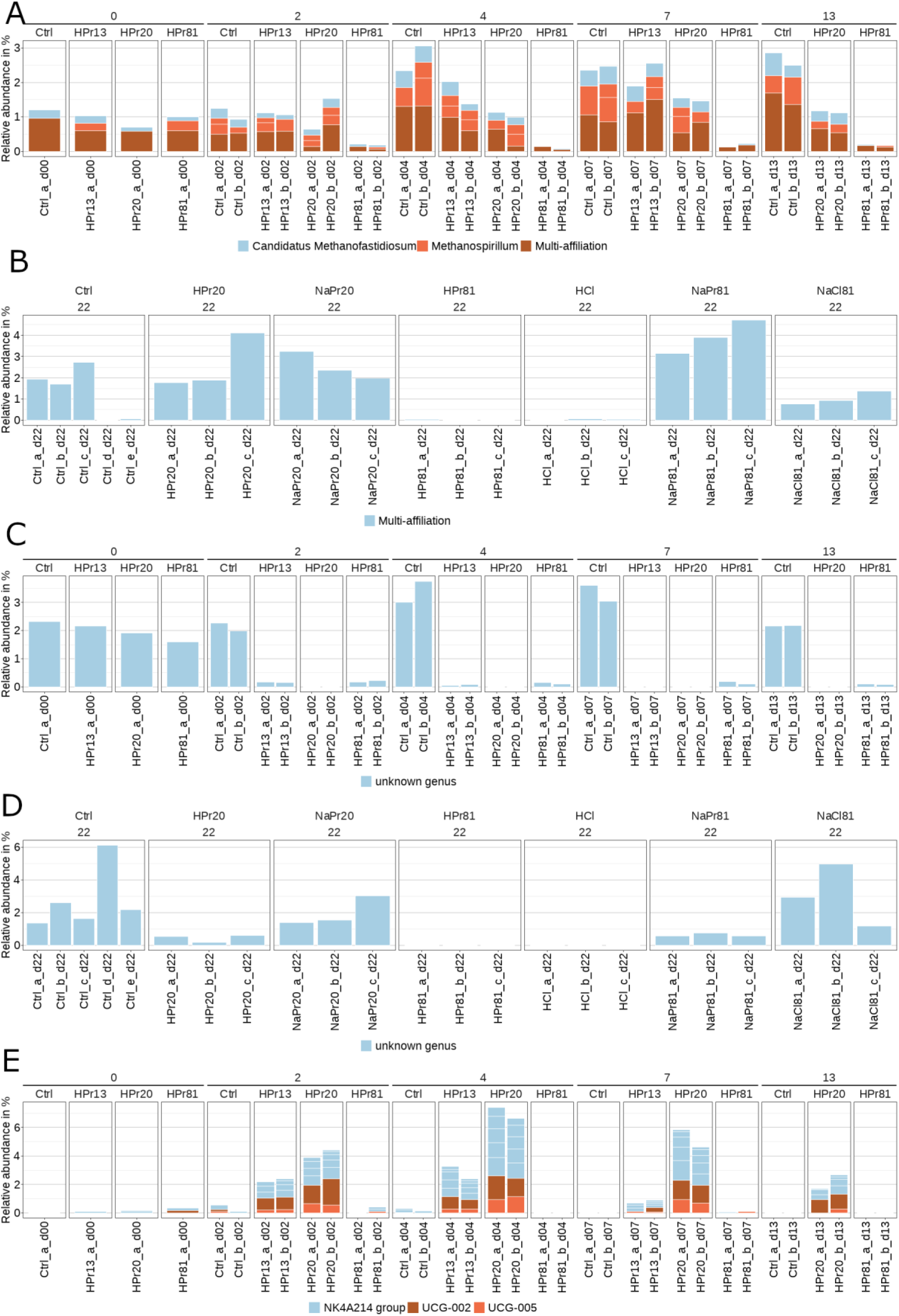
Barplots of selected microbial groups of interest. A. Relative abundance of archaea in the first tests, at the genus level. B. Relative abundance of the archaea in the second tests, at the genus level. C. Relative abundance of the family ST-12K33 (Sphingobacteriales order) in the first tests. D. Relative abundance of the family ST-12K33 (Sphingobacteriales order) in the second tests. E. Relative abundance of the family Oscillospiraceae in the first tests. For the first and second tests, multi-affiliation corresponds to *Methanothrix* and/or *Methanomicrobium*, most likely *Methanothrix*.

Based on PERMANOVA applied to the Bray-Curtis distance matrix of compositional data, 38% of the variance was explained by the addition of HPr and 24% by the incubation time (‘dist.bc ∼ treatment * day of incubation’, 9999 permutations, p-value = 10^-4^ for both factors). The dynamic adaptation of microbial communities to the presence of HPr was thus clearly demonstrated.

A pronounced decrease in the proportion of reads attributed to archaea as a function of the applied HPr concentration was observed (Figure 2A). At the end of the experiment (day 13), this proportion reached nearly 3% in control reactors, between 1% and 2% in reactors with intermediate HPr concentrations, and less than 0.2% in reactors exposed to the highest HPr concentrations.

### 3.4 HPr differentially affects over time a high number of microbial groups with diverse functions

To highlight microbial community patterns more thoroughly, a Common Component Analysis (CCA) was conducted. This methods groups variables by their shared effects on data dispersion, optimally weighting each variable’s contribution (Jouan-Rimbaud Bouveresse & Rutledge, 2024; Puig-Castellví et al., 2020). To extract relevant information from CCA, different visualizations were cross-referenced. For each common component (CC), Figure 3A illustrates the average scores of each treatment over time, revealing distinct longitudinal profiles occurring in the various treatments. Figure 3B presents a taxonomic tree of 75 distinct ASVs underlying these patterns (see also Supplementary File 2, Figure S5 and Supplementary File 4. Finally, the temporal evolution of these 75 ASVs was examined individually through relative abundance barplots (Supplementary File 5, Figures S6-S91), with Figure 3C illustrating an abundant ASV associated with CC1, assigned to the family Anaerolineaceae.

**Figure 3.**
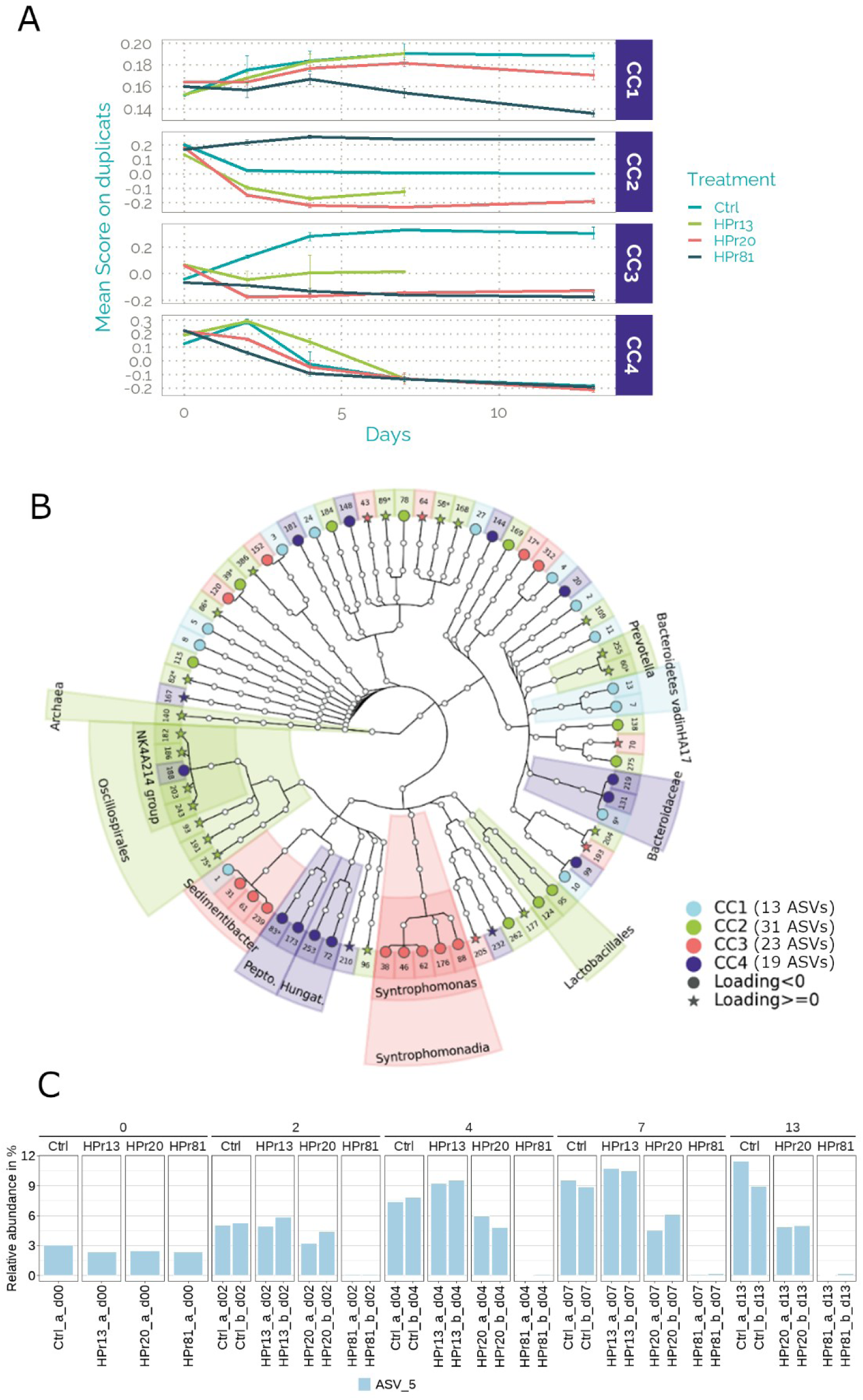
Illustration of Common Component Analysis (CCA) results for the first tests. A. Profiles over time of the AD reactors for the 4 first Common Components (CC); the average and standard deviation are shown, 2 replicates. B. Taxonomic tree of the ASVs contributing the most to the four first CCs. These ASVs were selected based on their loadings, exceeding 1.5 times the standard deviation of all ASVs (Supplementary File 2, Figure S5). C. Example of an ASV associated to CC1 (ASV_5: Bacteria, Chloroflexi, Anaerolineae, Anaerolineales, Anaerolineaceae, unknown genus).

Clear and distinct trends of microbial community dynamics were observed across the four CCs (Figure 3A). CC1 exhibited a particular trend for the highest HPr concentration (HPr81), noticeable from day 4 and intensifying progressively over time. CC2 discriminated high (HPr81), intermediate (HPr20), and low (HPr13 and Ctrl) HPr concentrations. CC3 revealed distinct evolutions between Ctrl and other treatment groups. Finally, CC4 exhibited consistent trends among the different treatments, primarily reflecting temporal evolutions: such ecological successions are anticipated due to progressive substrate depletion and bioconversion under batch operating conditions.

In more detail, the 13 underlying ASVs identified for CC1, representing 12 families and 6 phyla, were among the most abundant, with most exceeding 5% relative abundance in at least one sample. Four ASVs—assigned to Anaerolineaceae, Sedimentibacteraceae, Dysgonomonadaceae, and Bacteroidetes vadinHA17—were strongly impaired under high HPr exposure (HPr81), with ASV_1 (*Sedimentibacter*) being particularly sensitive even to minimal HPr additions. In contrast, three ASVs—from Candidatus Cloacimonadaceae, Tannerellaceae, and Lysobacteraceae—were selected under HPr81. Based on MIDAS database (Dueholm et al., 2024), these 13 ASVs were predominantly acidogens, consistent with the abundance of fermentable molecules in wastewater sludge.

The 31 ASVs associated to CC2 were distributed over 22 families, mostly from the phyla Firmicutes, Bacteroidota and Proteobacteria. Corresponding to subdominant taxa, they encompassed a wide range of functional groups. Notably, important shifts in fermenter populations were observed with increasing HPr concentrations: six ASVs assigned to Oscillospiraceae (Gophna et al., 2017) were selected under low to intermediate HPr concentrations, while ASV_60 (*Prevotella*) and ASV_95 (*Lactobacillus*) were favored under HPr81, reaching more than 6% and 3% of the reads, respectively. Besides, three ASVs revealed highly sensitive to HPr, namely, ASV_140 (*Methanospirillum*), ASV_39 (*Smithella*) and ASV_17 (Sphingobacteriales, ST-12K33).

CC3 corresponded to subdominant taxa, with 13 out of 23 ASVs more abundant under Ctrl. Eleven of these, including *Sedimentibacter* and *Syntrophomonas* members, were highly sensitive to HPr. *Sedimentibacter* presumably corresponded to protein fermenters (Imachi et al., 2016). *Syntrophomonas* bacteria oxidize butyrate or saturated fatty acids, producing acetate and H_2,_ and, in some cases, propionate, depending on the substrate (Sekiguchi, 2015). Consequently, they can develop syntrophic interactions with hydrogenotrophic methanogens (Sousa et al., 2007). The remaining 10 underlying ASVs displayed various patterns depending on HPr concentration: some reached peak abundance at intermediate HPr concentrations (from families Tannerellaceae, Butyricicoccaceae) while others were more abundant under the highest HPr concentration (from families Veillonellaceae and Paludibacteraceae).

Overall, CCA confirmed that exposure to HPr affected the abundance of numerous ASVs across diverse functional and phylogenetic groups, leading to several major shifts in microbial composition as the initial HPr concentration increased.

### 3.5 Each added compound affects specifically the microbial community composition

To determine the extent to which the adaptation of the microbial community was driven by the specific compound added, and in particular by the nature of the counterion (e.g. H^+^ or Na^+^), *16S rDNA* metabarcoding was applied to samples from the second tests, collected in each reactor at the end of the experiment, after 22 days of incubation. Out of 18 different bacterial phyla, the six dominant ones were Bacteroidota, Bacillota, Candidatus Cloacimonadota, Pseudomonadata, Chloroflexota, and Synergistota (Supplementary File 6, Figure S93). This composition was similar to the first test except for Synergistota instead of Spirochaetota; Spirochaetota was still highly represented in certain samples of the second tests. Methanogenic archaea belonged to *Methanothrix* and/or *Methanomicrobium*. Their proportion varied significantly across treatments (Figure 2B), reaching approximately 2-3% in control reactors, slightly higher proportions under NaPr conditions, and slightly lower ones under NaCl conditions, compared to Ctrl. Archaea were detected in extremely low proportions in the most inhibited reactors (HPr81 and HCl).

To identify the microbial groups specifically influenced by the treatments at a finer taxonomic level, differential analysis was conducted, targeting the ASVs of archaea and bacteria (Figures 4-5). Of the 276 ASVs, 145 were differentially abundant compared to Ctrl, with 122 less abundant and only 26 more abundant in at least one treatment, likely highlighting the limits of microbial community adaptation over the short term. Adaptation was condition-specific: all treatments except NaPr20 had at least one uniquely differentially abundant ASV (Supplementary File 6, Figure S94). Besides, increasing the concentration of the compound of interest led to the detection of additional differentially abundant ASVs. This probably reflected variable toxicity thresholds among microorganisms, with a greater number of strains being affected as the concentration increased. Acidity had a stronger impact than non-acidic compounds, with >100 differentially abundant ASVs under HPr81 or HCl versus <30 under other conditions.

**Figure 4.**
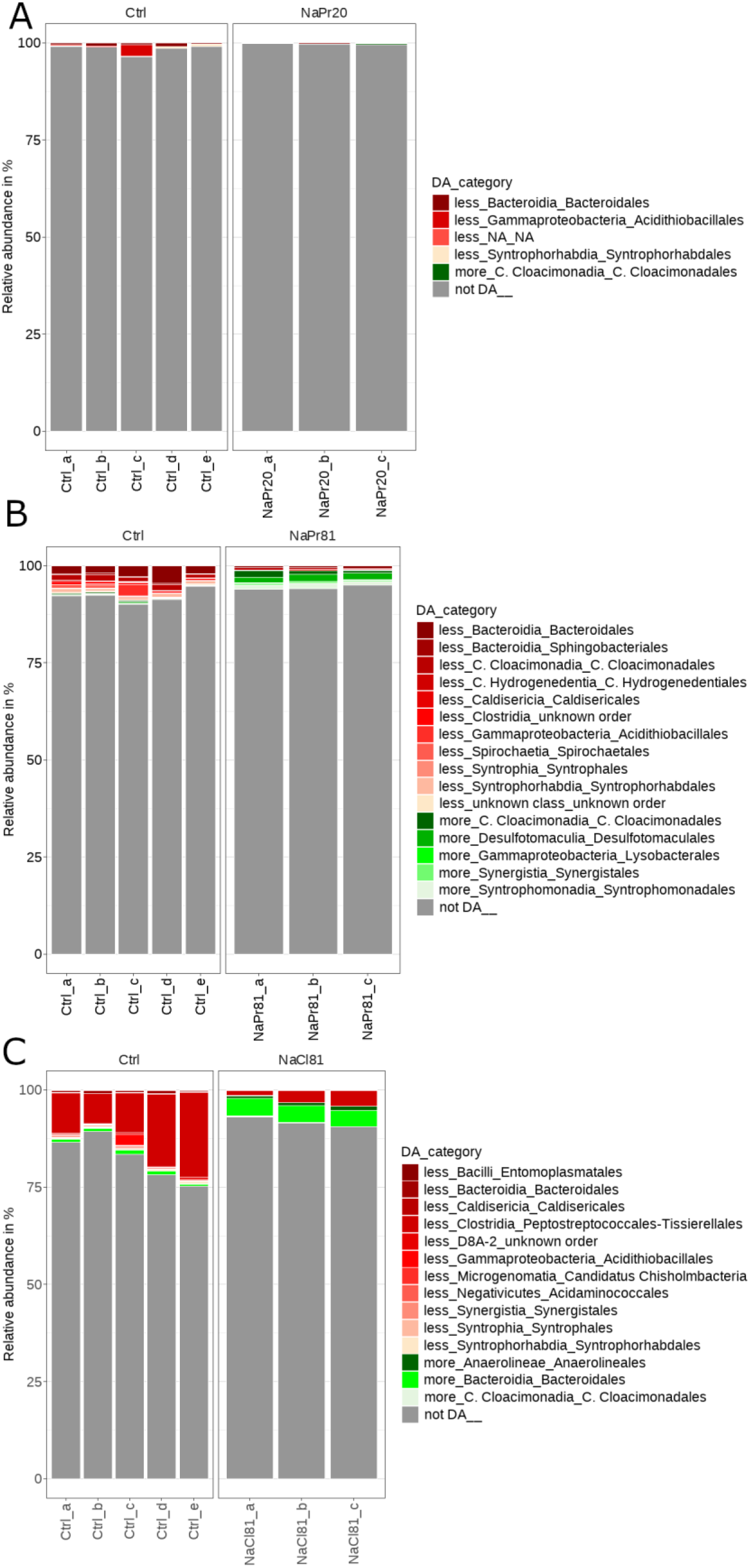
Taxonomy and relative abundance of differentially abundant ASVs identified under various non-acidifying conditions, compared to Ctrl. Red indicates the ASVs that were significantly less abundant in the treatment compared to Ctrl, and green, significantly more abundant in the treatment compared to Ctrl. A. Comparison of NaPr20 to Ctrl. B. Comparison of NaPr81 to Ctrl. C. Comparison of NaCl81 to Ctrl.

**Figure 5.**
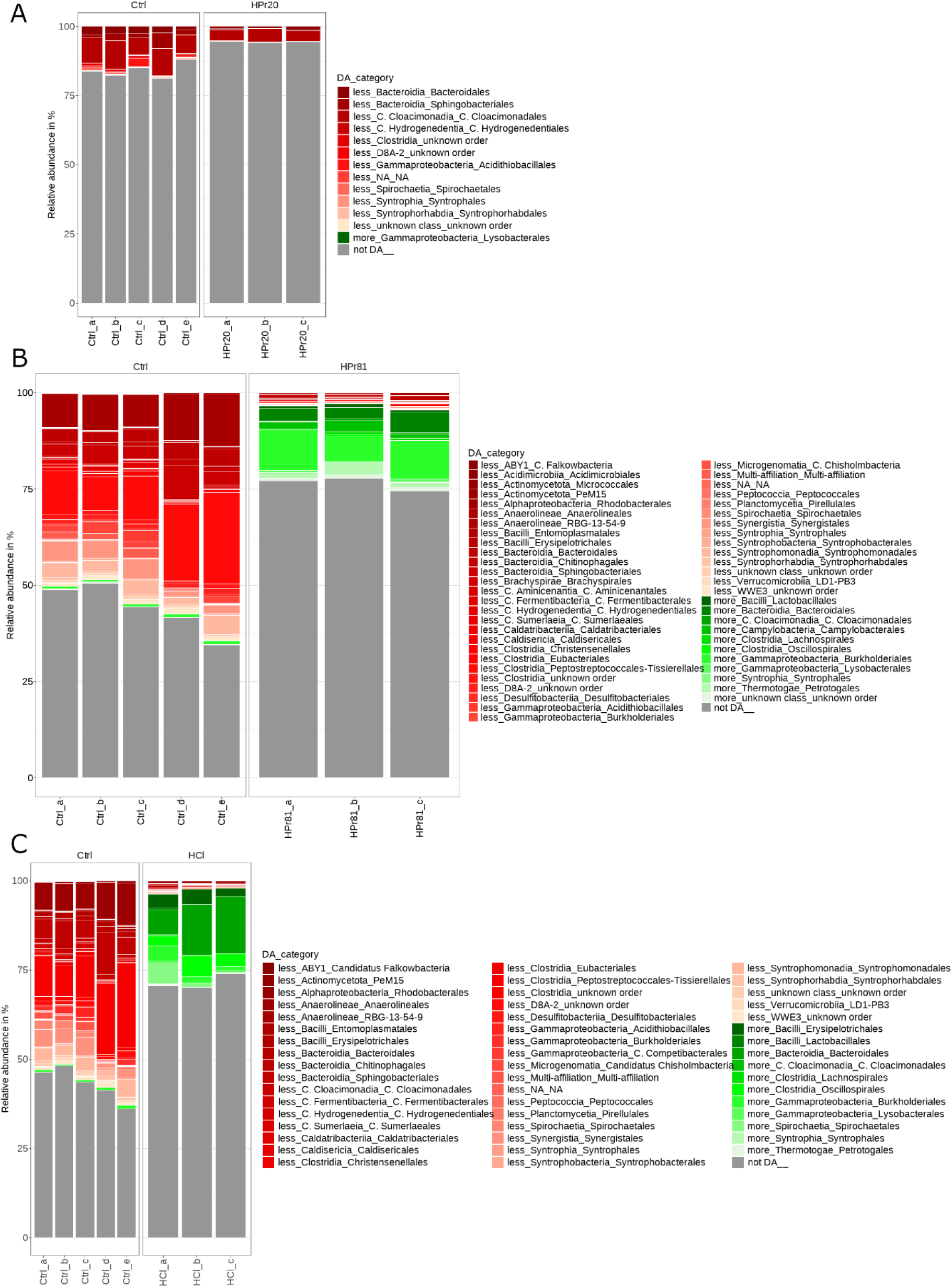
Taxonomy and relative abundance of differentially abundant ASVs identified under various acidifying conditions, compared to Ctrl. Red indicates the ASVs that were significantly less abundant in the treatment compared to Ctrl, and green, significantly more abundant in the treatment compared to Ctrl. A. Comparison of HPr20 to Ctrl. B. Comparison of HPr81 to Ctrl. C. Comparison of HCl to Ctrl.

Finally, the number of differentially abundant ASVs was negatively correlated with methane production rates (R² = 0.9698, Supplementary File 6, Figure S95), underscoring the link between AD impairment and microbial community reshaping.

### 3.6 Adaptation of microbial community to propionate (Pr^−^) without acidity shift

Microbial community adaptation to Pr^-^ under non-acidic conditions was assessed by examining conditions NaPr20, NaPr81 and NaCl in the second tests (Figure 4).

Under NaPr81 (Figure 4B),12 ASVs were identified as highly robust regarding adaptation to Pr^-^, as they were differentially abundant when comparing NaPr81 to Ctrl and NaPr81 to NaCl81, but not when comparing NaCl81 to Ctrl (Supplementary File 7). Among them, the most abundant ones (reaching approximately 1% in at least one sample) and those showing strong reproducibility were examined more specifically. In details, the abundances of ASV_2 (order Bacteroidales, family Prolixibacteraceae) and ASV_208 (order Syntrophorhabdales, genus *Syntrophorhabdus*) were reduced under NaPr81, whereas ASV_68 (order Lysobacterales, genus *Thermomonas*) and ASV_703 (order Synergistales, family Synergistaceae) were on the contrary favored under the same condition. Members of Prolixibacteraceae are facultative anaerobes, capable of fermenting certain carbohydrates (Huang et al., 2014; Zhou et al., 2019). The sole isolated bacterium belonging to *Syntrophorhabdus* can oxidize aromatic compounds in syntrophy with hydrogenotrophic methanogens (Qiu et al., 2008). Some *Thermomonas* strains can assimilate propionate (Denner et al., 2015), which may have explained their higher abundance under the high NaPr concentration. According to the MIDAS database (Dueholm et al., 2024), Synergistaceae bacteria ferment certain acids and amino acids but not systematically carbohydrates, producing H_2_ and other metabolites, sometimes propionate.

NaPr20 had a limited effect on both methane production and microbial community composition, with only ASV_109 (Bacteroidales, Rikenellaceae, S50 wastewater-sludge group) and ASV_453 (Syntrophorhabdales, *Syntrophorhabdus*) showing significant differential abundance. Rikenellaceae includes obligate anaerobic microorganisms, such as fermenters, according to the MIDAS database (Dueholm et al., 2024).

### 3.7 Adaptation of microbial community to propionate (Pr^-^) with acidity shift

To decipher microbial community adaptation to Pr^-^ under acidic conditions, HPr20, HPr81 and HCl were compared to Ctrl in the second tests (Figure 5). Under HPr81, 96 ASVs were significantly impaired, the most abundant being ASV_1 (order Peptostreptococcales-Tissierellales, genus *Sedimentibacter*) and ASV_4 (order Anaerolineales, family Anaerolineaceae, probable fermenters). Only fourteen ASVs were significantly more abundant. Among them, ASV_14 and ASV_35 were both affiliated to Bacteroidales, Tannerellaceae family; they reached more than 1% in average under HPr81, and were not detected under Ctrl. Known Tannerellaceae members are strictly anaerobic carbohydrate fermenters according to MIDAS (Dueholm et al., 2024). ASV_112 was also notable: affiliated to *Lactobacillus* genus (Lactobacillales), it reached 0.74% under HPr81. *Lactobacillus* corresponds to saccharolytic fermenters, facultative anaerobes, producing lactate as major end-product, based on MIDAS (Dueholm et al., 2024). HPr81 and HCl reactors shared specifically 64 negatively-affected ASVs (including ASV_1 and ASV_4) and 10 positively-affected ASVs (including ASV_14, ASV_35 and ASV_112), (Supplementary File 6, Figure S94) suggesting that under HPr81, community adaptation was largely driven by the acidic pH.

Regarding HPr20, three ASVs were notable (Figure 5C) since they both exhibited significant abundances (≥ 1% in at least one treatment) and were less abundant relative to Ctrl: ASV_6 (order Candidatus Cloacimonadales, genus W5), ASV_16 (order Bacteroidales, family Rikenellaceae, genus DMER64), and ASV_21 (order Sphingobacteriales, family ST-12K33). Importantly, genus W5 is believed to correspond to syntrophic propionate oxidizing bacteria (Dyksma & Gallert, 2019). Genus DMER64 is a poorly characterized, potential syntrophic group (Lee et al., 2019). Although less pronounced, it is also worth noting the presence of ASV_68 (order Lysobacterales, genus *Thermomonas*) in two out of three HPr20 reactors, while it was not detected in any of the Ctrl reactors and was also slightly but clearly selected under HPr81 and HCl (Supplementary File 7).

## 4. Discussion

A previous study evidenced that anaerobic digesters can tolerate propionate concentrations as high as 50 mM of NaPr (4,800 mg L^-1^) (Ahring et al., 1995). More recently, it was shown that 41.6 mM of NaPr (4,000 mg L^-1^) can be fully degraded in batch reactors (Yang et al., 2015). The present study is consistent with the moderate resilience of AD reactors to NaPr molar concentrations up to 81 mM (7,750 mg L^-1^), which is far higher than the threshold reported for its acid form (12 mM of HPr, equivalent to 900 mg L^-1^) (Wang et al., 2009; Han et al., 2020). A recent publication (Ochoa-Bernal et al., 2025) confirms the accumulation of acetate and the appearance of a shift in archaeal populations between 62.5 and 125 mM (i.e. 6,000 – 12,000 mg L^-1^) of NaPr, close to the threshold of inhibition that we observed in this study.

The maximum VFA concentration at which AD can operate depends on numerous factors. On one hand, the propionate concentration is always reported relative to the media volume. It does not accurately describe how propionate locally surrounds and interacts with microbial cells. The ratios between inoculum, raw sludge and propionate concentrations in reactors could play a key role and partially explain the differences observed among various studies (e.g. 6.4 g VS L^-1^ / 54 mM in Han et al. (2020) versus 22.4 g VS L^-1^ / 81 mM in the present study). On the other hand, the tolerance of AD reactors to propionic acid and its conjugated base differs greatly. The present study distinguishes between pH drop and propionate effects on methanogenesis inhibition, providing a unifying view.

The first and second tests were conducted under very similar conditions. Consistently, numerous common trends regarding microbial community changes emerged when comparing both tests, as summarized in Figure 6. For both HPr and NaPr, gradual shifts in community composition according to the compound concentration were clearly visible, highlighting distinct ranges of sensitivity and adaptation ability. Beyond this shared general trend, results were different for both compounds. The rearrangements of the microbial community in the presence of high NaPr concentration (NaPr81) were relatively modest in terms of microbial abundance, even though the methane production rate was 40% lower than in the control conditions. By contrast, under HPr81, the highly abundant fermenters *Sedimentibacter* (Sedimentibacteraceae) and Anaerolineaceae were affected, as well subdominant key groups such as methanogens. In addition, several microbial groups specifically affected by the presence of NaPr were identified, whereas in the presence of HPr, the effect appeared to be primarily linked to acidity, with many microbial groups similarly affected in the presence of HCl. In both test series, the proportion of methanogenic archaea decreased markedly with increasing HPr concentration, consistent with the inhibition of methane production. However, the archaeal populations selected were not identical across both test series. This discrepancy may reflect the use of distinct sludge batches for each test series, which, despite originating from the same plant and digester, may have differed slightly in composition. Moreover, since archaea operate at the terminal step of the AD process, they may be particularly sensitive to the cumulative effect of small variations occurring at each upstream stage.

**Figure 6.**
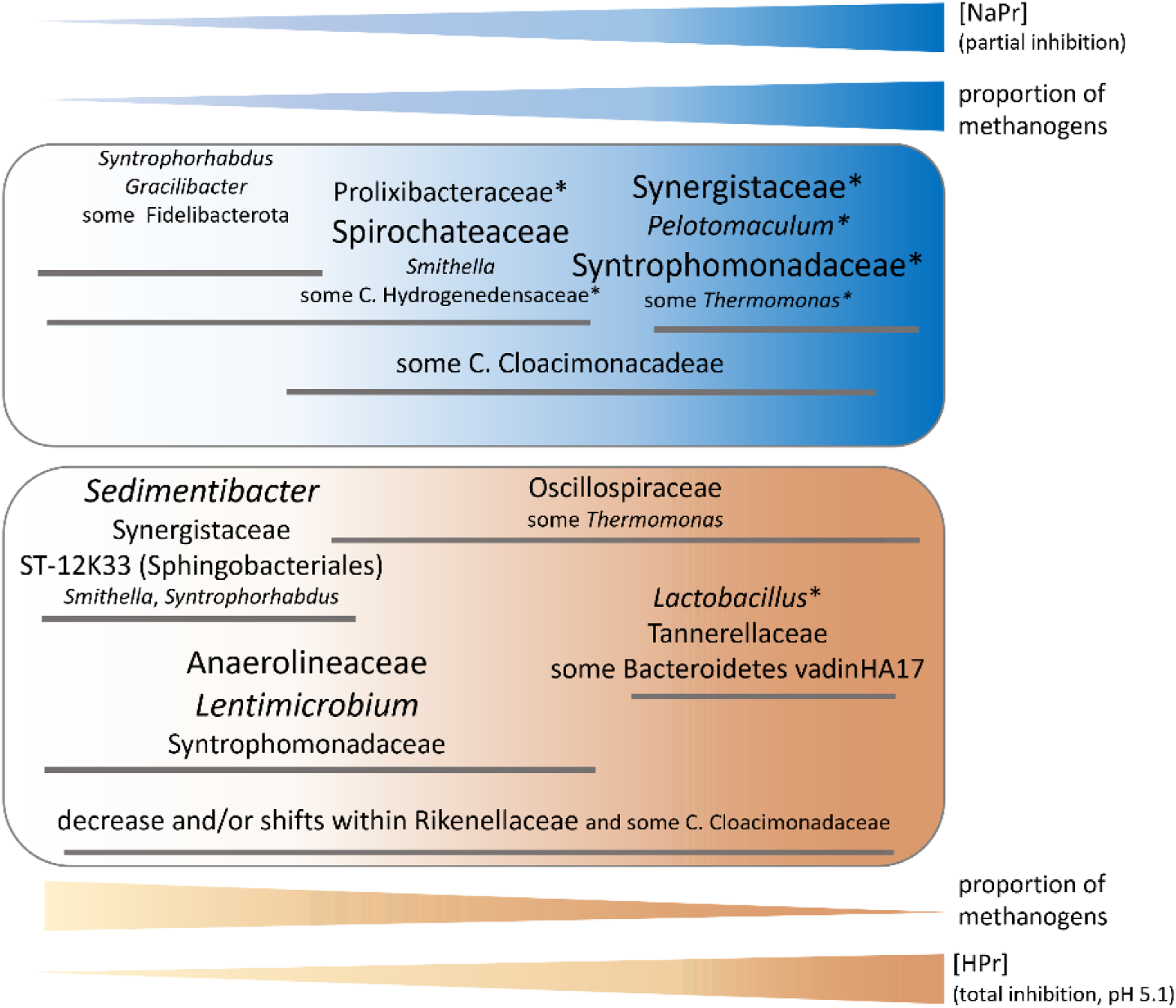
Summary illustration of the adaptation of anaerobic digestion communities exposed to various concentrations of NaPr (top) and HPr (bottom). Asterisks indicate that the effect observed was specific for the tested conditions, i.e. not observed under equivalent NaCl (top) or HCl (bottom) treatment. The font size reflects the abundance of the microbial groups involved.

Worth highlighting, in a previous study available as preprint (Camargo et al, 2023, bioRxiv), acid inhibition with HPr accumulation naturally occurred during the dry co-digestion of wastewater treatment sludge and horse manure, due to high organic loading; some of the microorganisms negatively affected under these conditions were similar to those identified in the present study. These included members of the families Sedimentibacteraceae, Smithellaceae, and ST-12K33 (Sphingobacteriales), which were also inhibited by HPr in the present study. *Sedimentibacter* was also negatively affected during the semi-continuous mesophilic digestion of dairy manure disrupted through HPr spikes (Khafipour et al., 2020). The same study documented a decrease in the abundance of *Syntrophomonas* in response to HPr, which is also consistent with observations in the present study.

When propionic acid was added to the reactors, one might have expected antagonistic effects on propionate-consuming bacteria: on the one hand, selection of these bacteria due to an increase in available substrate; on the other hand, a negative effect due to acidic inhibition. In the present dataset, a significant number of families possibly contained SPOB (Westerholm et al., 2022). However, this activity was difficult to confirm on the basis of *16S rDNA* sequencing alone. Indeed, the selected method yielded taxonomic assignation at the genus level, and it is known that not all species within a given genus are SPOB. Based on previous knowledge and on the abundance profiles, SPOB in the present study could presumably belong to *Smithella* (detected under non-inhibiting conditions), *Pelotomaculum* (order Desulfotomaculales, selected under high NaPr concentration), *Syntrophomonadaceae* (inhibited by HPr but enriched under high NaPr concentrations), and to members of Candidatus Cloacimonadaceae (showing variable responses depending on conditions) (Westerholm et al., 2022).

## 5. Conclusions

This study provides new insights into propionic acid inhibition during anaerobic digestion of municipal sewage sludge. While 20 mM HPr caused a partial inhibition of methane production (22% reduction in the maximal production rate), 81 mM HPr led to complete inhibition, primarily driven by the associated pH drop to 5.1 rather than by propionate ions themselves — as evidenced by the fact that 81 mM NaPr only reduced the methane production rate by 40% while leaving pH unaffected. Propionate ions (Pr⁻) nonetheless exert a secondary, concentration-dependent inhibitory effect even under neutral pH conditions. Microbial community analysis revealed a substantial and gradual adaptive response, insufficient to fully offset the inhibition of methane production. Community restructuring was compound-specific, with HPr-treated reactors sharing numerous features with HCl controls, underscoring the dominant role of acidity, while NaPr induced distinct and more limited microbial shifts. Diverse functional groups were affected, including syntrophs and methanogenic archaea. These findings reconcile conflicting reports in the literature by distinguishing the roles of pH, undissociated HPr, and Pr⁻ ions, and carry direct practical implications: microbial community profiling could serve as an early warning tool for process imbalance detection. Further research under continuous feeding conditions could help identify operational biomarkers.

## Acknowledgements

This study was funded by the French research program MOCOPEE.

## Author Contributions

**XL**: Conceptualization, Formal analysis, Writing – original draft, **CS**: Investigation, Formal analysis, Writing – review and editing **VJ**: Data curation, Formal analysis, **AP**: Conceptualization, Supervision, Writing – review and editing, **LA**: Investigation, Conceptualization, Supervision, Writing – review and editing, **TR**: Conceptualization, Supervision, Writing – review and editing, **S G-R**: Conceptualization, Supervision, Writing – review and editing **VR**: Conceptualization, **CL**: Investigation, Conceptualization, **CB**: Investigation, Writing – review and editing, **CM**: Software, Writing – review and editing, **OC**: Conceptualization, Writing – review and editing, **AB**: Conceptualization, Formal analysis, Writing – original draft, **C R-A**: Conceptualization, Supervision, Writing – review and editing.

