## Supplementary file 5 for "Propionic acid-related inhibition during anaerobic digestion: insights into methane production and microbial community adaptation"

### Supplementary Figures S6 to S91

Barplots of the relative abundance of 75 distinct ASVs contributing the most to the patterns obtained with the Common Component Analysis applied to the 16S rDNA metabarcoding data of the first tests.

The ASVs are grouped according to the CC. A same ASV can be identified for several CCs, so that the total number of barplots is 86.

## CC1

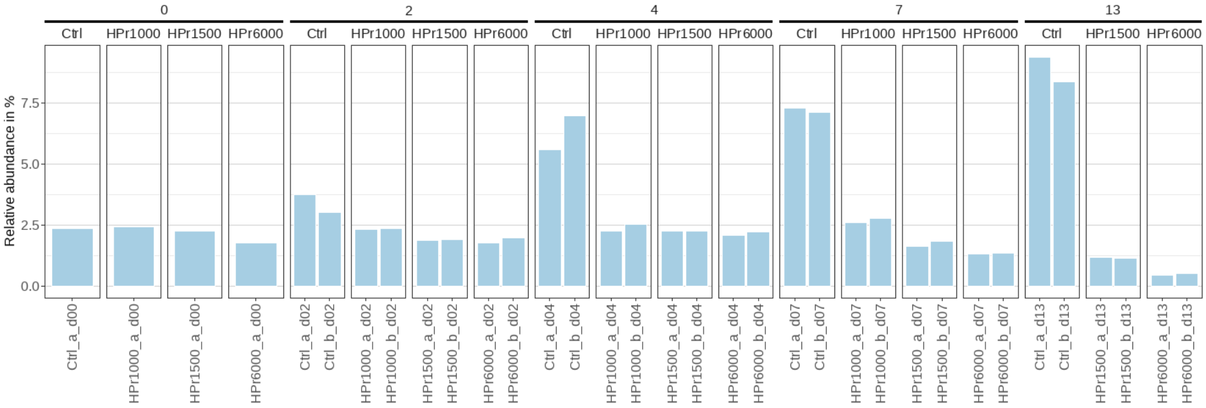


Figure S6. ASV 1. Bacteria, Firmicutes, Clostridia, Peptostreptococcales-Tissierellales, Sedimentibacteraceae, Sedimentibacter, unknown species

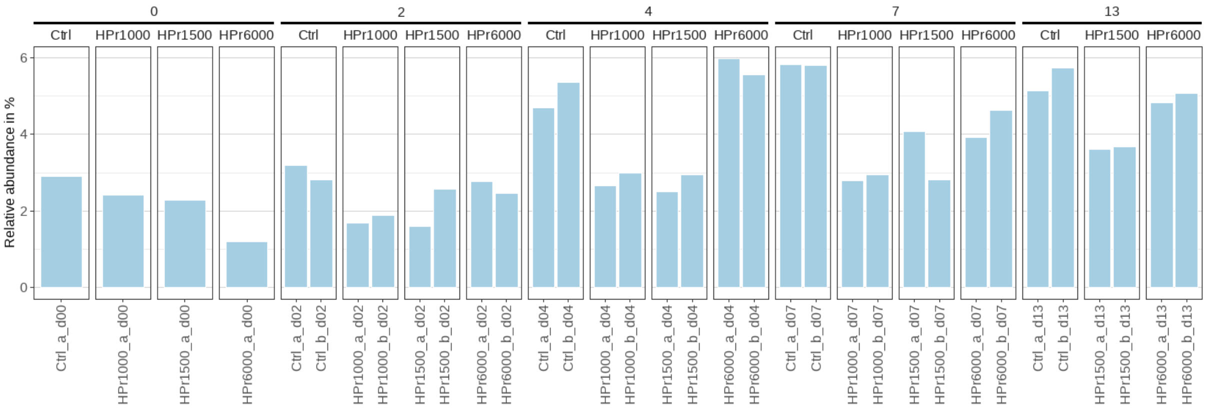


Figure S7. ASV 2. Bacteria, Bacteroidota, Bacteroidia, Bacteroidales, Prolixibacteraceae, unknown genus, unknown species

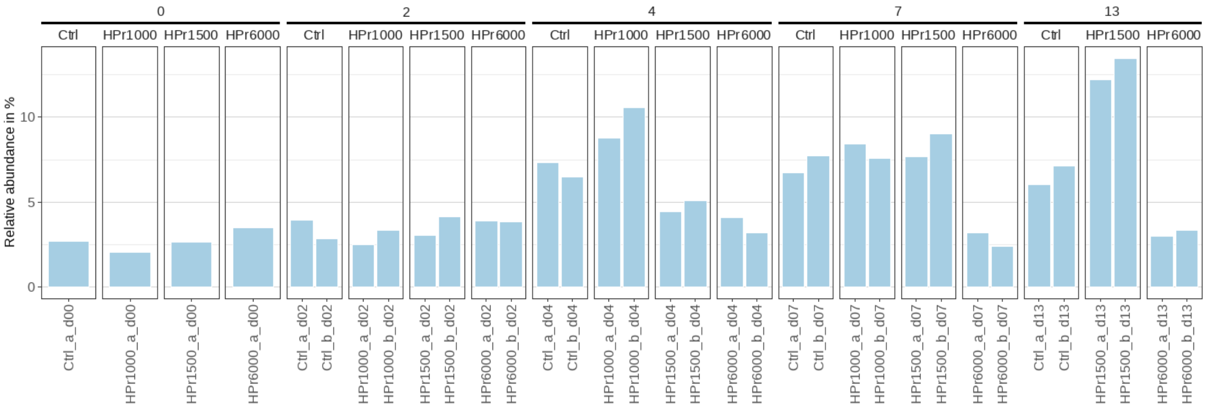


Figure S8. ASV 3. Bacteria, Spirochaetota, Spirochaetia, Spirochaetales, Spirochaetaceae, unknown genus, unknown species

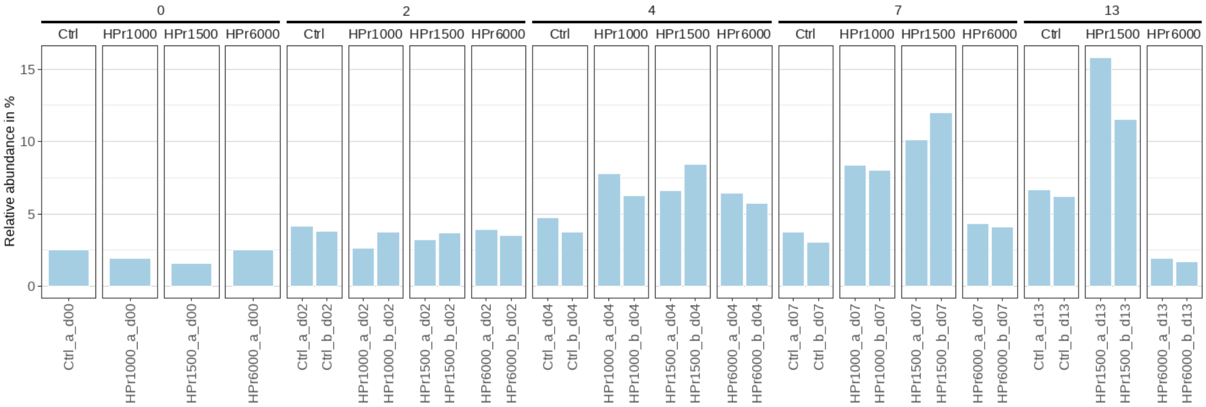


Figure S9. ASV 4. Bacteria, Bacteroidota, Bacteroidia, Sphingobacteriales, Lentimicrobiaceae, Lentimicrobium, Bacteroidetes bacterium ADurb.BinA012

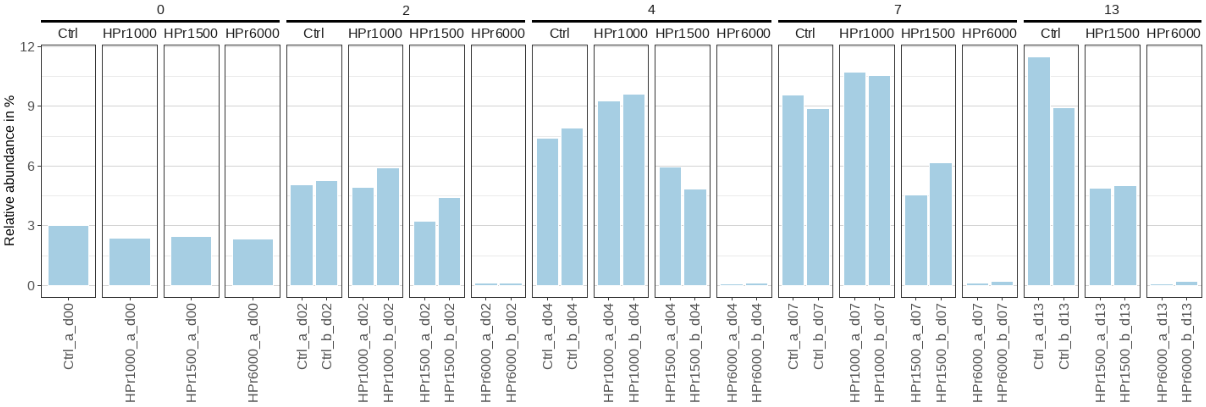


Figure S10. ASV 5. Bacteria, Chloroflexi, Anaerolineae, Anaerolineales, Anaerolineaceae, unknown genus, Multi-affiliation

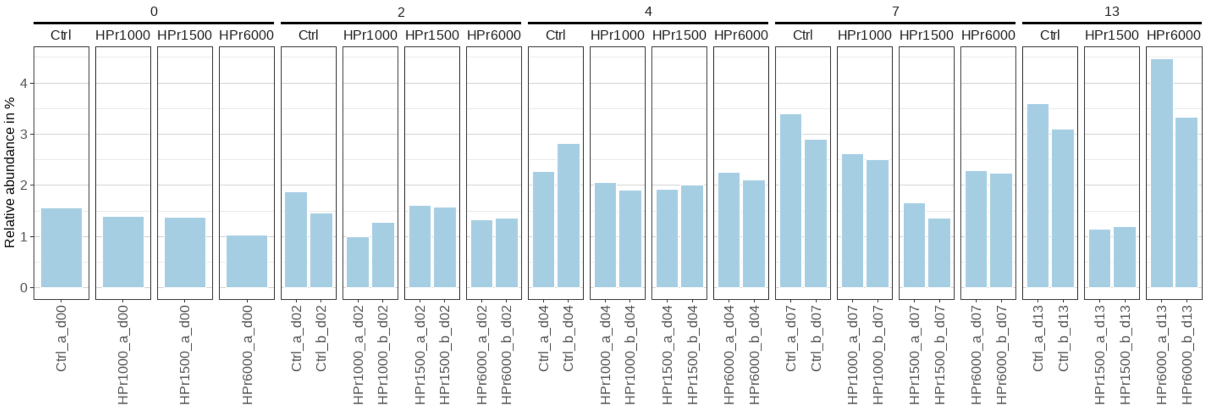


Figure S11. ASV 7. Bacteria, Bacteroidota, Bacteroidia, Bacteroidales, Bacteroidetes vadinHA17, unknown genus, Multi-affiliation

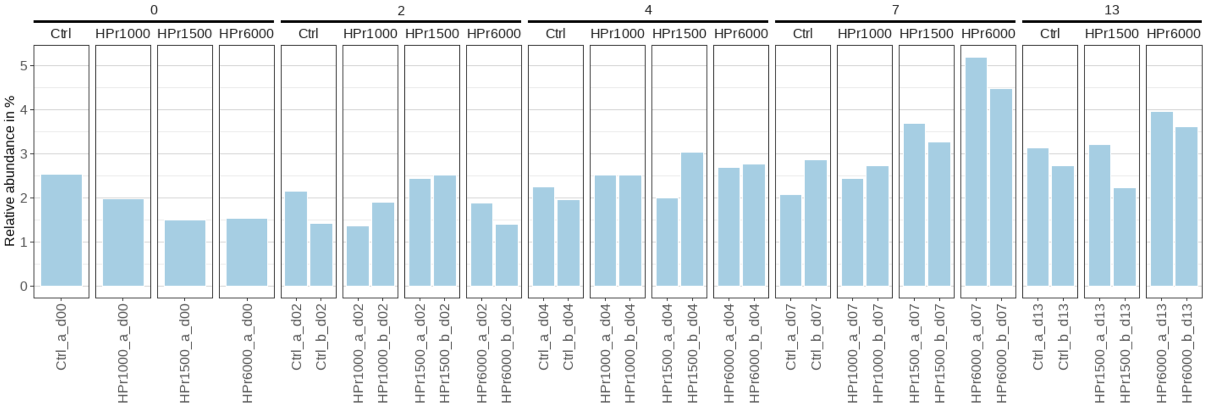


Figure S12. ASV 8. Bacteria, Cloacimonadota, Cloacimonadia, Cloacimonadales, Cloacimonadaceae, Candidatus Cloacimonas, metagenome

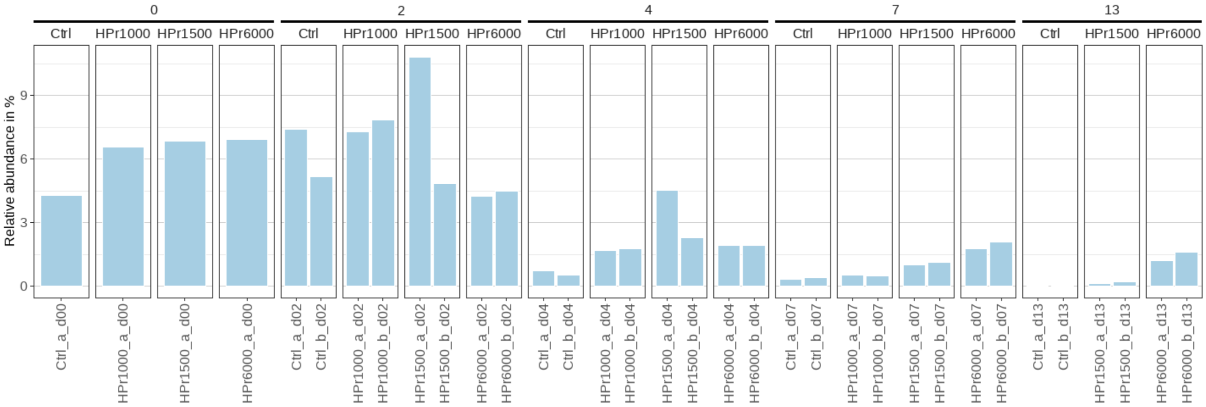


Figure S13. ASV 9. Bacteria, Bacteroidota, Bacteroidia, Bacteroidales, Bacteroidaceae, Bacteroides, Multi-affiliation

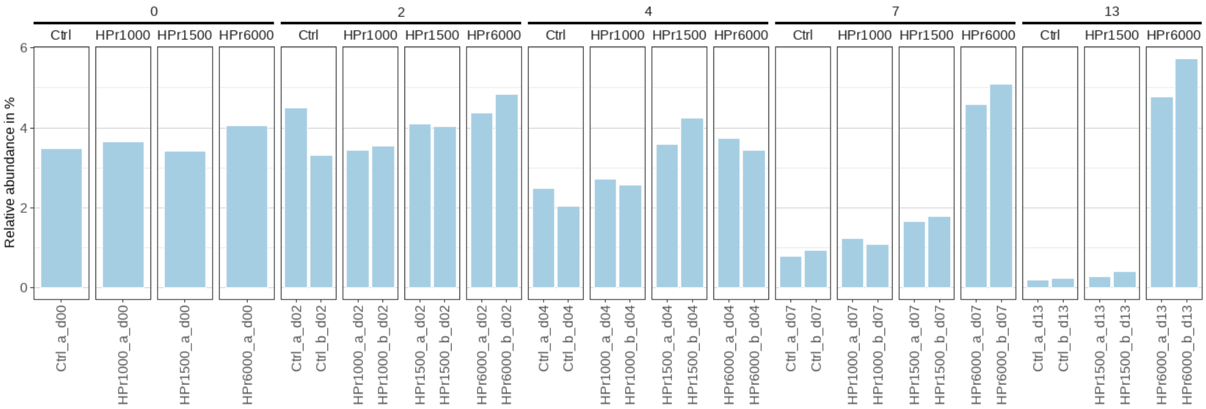


Figure S14. ASV 10. Bacteria, Bacteroidota, Bacteroidia, Bacteroidales, Tannerellaceae, Macellibacteroides, Multi-affiliation

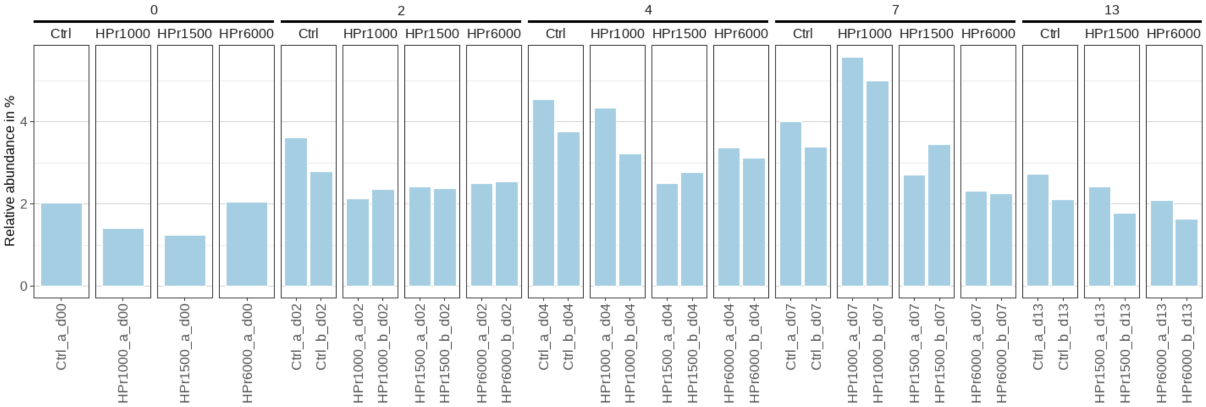


Figure S15. ASV 11. Bacteria, Bacteroidota, Bacteroidia, Bacteroidales, Dysgonomonadaceae, Proteiniphilum, Multi-affiliation

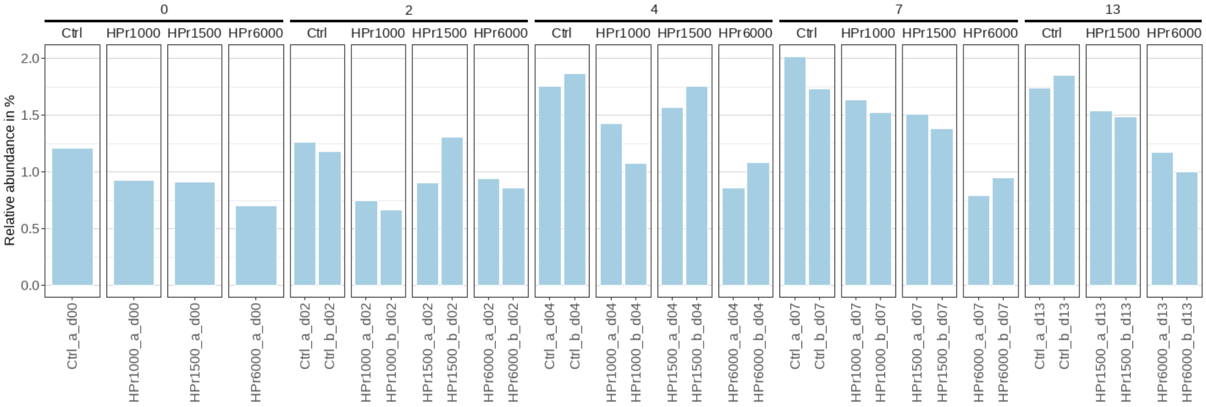


Figure S16. ASV 13. Bacteria, Bacteroidota, Bacteroidia, Bacteroidales, Bacteroidetes vadinHA17, unknown genus, unknown species

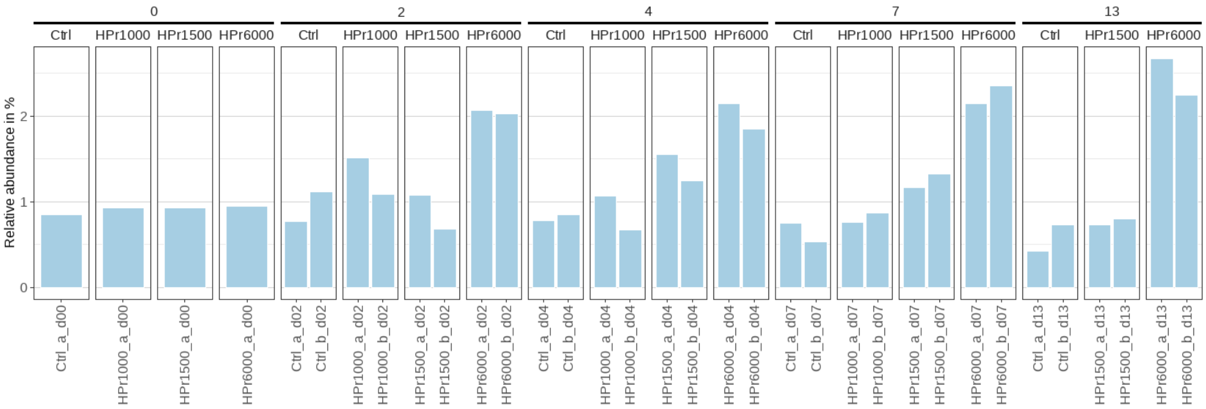


Figure S17. ASV 24. Bacteria, Proteobacteria, Gammaproteobacteria, Xanthomonadales, Xanthomonadaceae, Thermomonas, unknown species

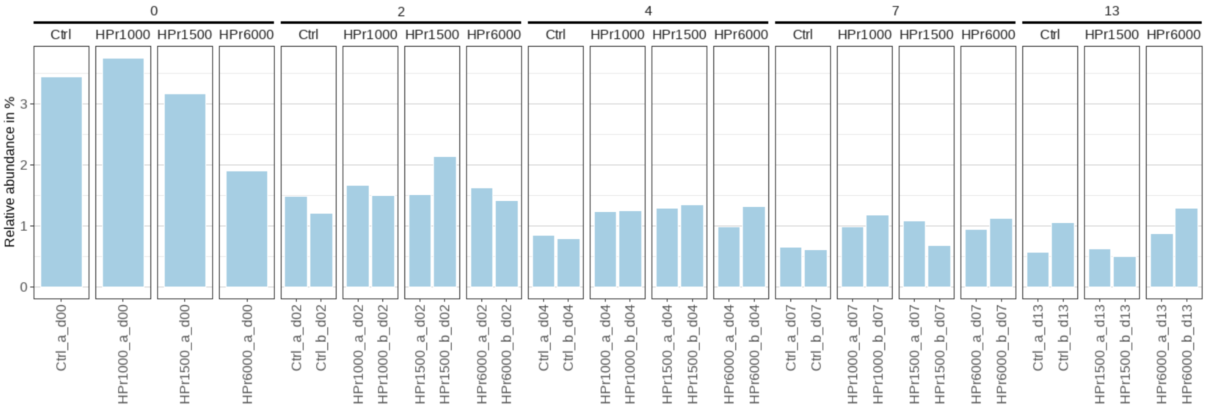


Figure S18. ASV 27. Bacteria, Proteobacteria, Gammaproteobacteria, Burkholderiales, Comamonadaceae, Acidovorax, Multi-affiliation

## CC2

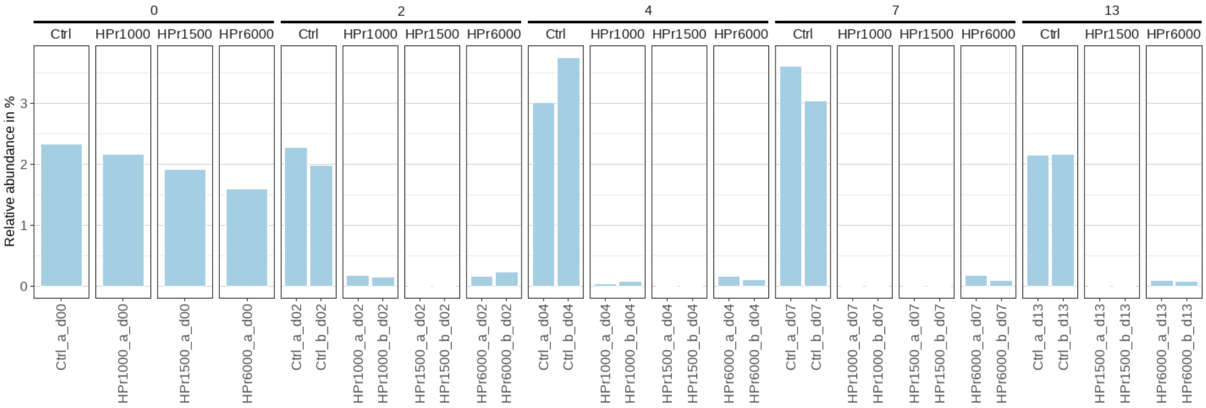


Figure S19. ASV 17. Bacteria, Bacteroidota, Bacteroidia, Sphingobacteriales, ST-12K33, unknown genus, metagenome

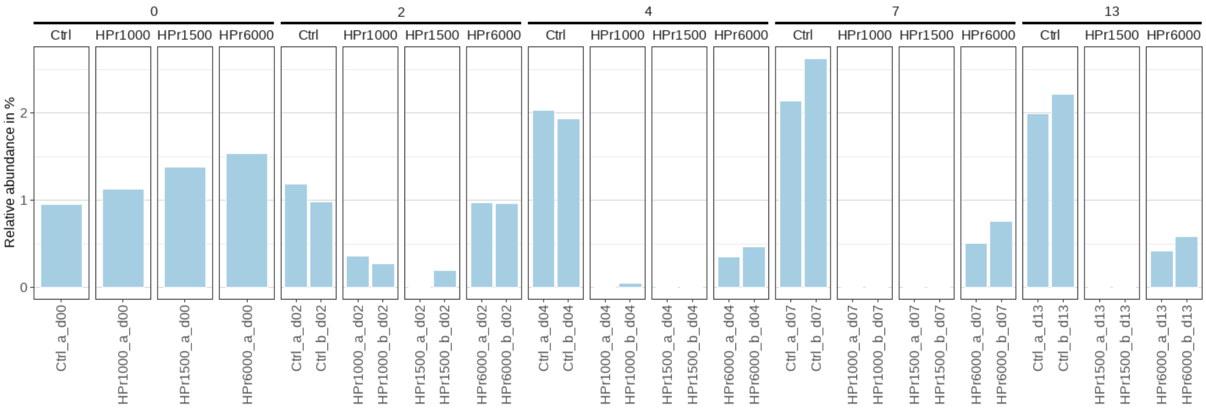


Figure S20: ASV 39. Bacteria, Desulfobacterota, Syntrophia, Syntrophales, Smithellaceae, Smithella, Multi-affiliation

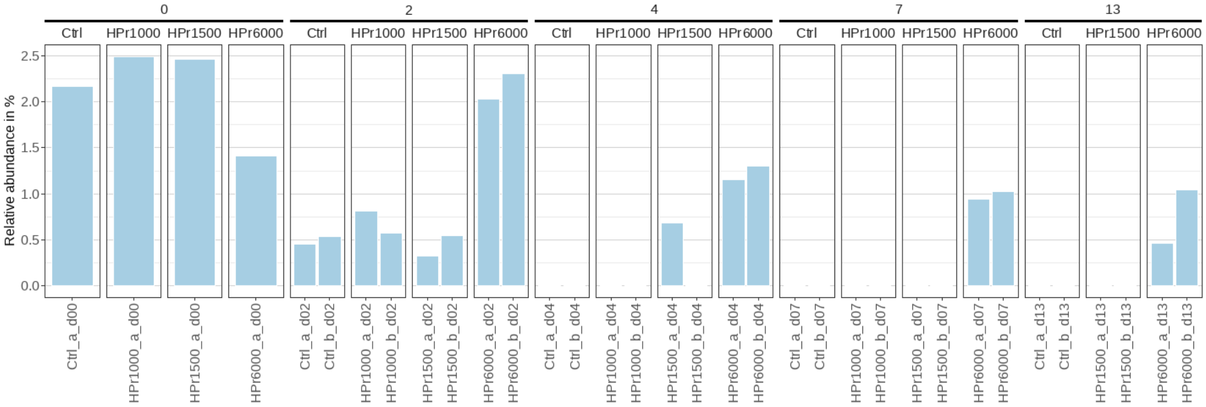


Figure S21: ASV 58. Bacteria, Proteobacteria, Gammaproteobacteria, Burkholderiales, Comamonadaceae, Hydrogenophaga, unknown species

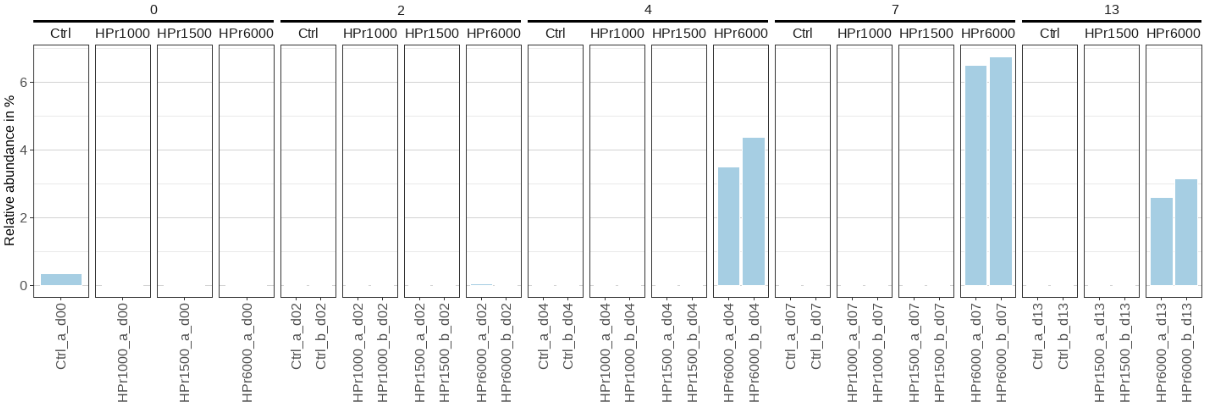


Figure S22: ASV 60. Bacteria, Bacteroidota, Bacteroidia, Bacteroidales, Prevotellaceae, Prevotella, unknown species

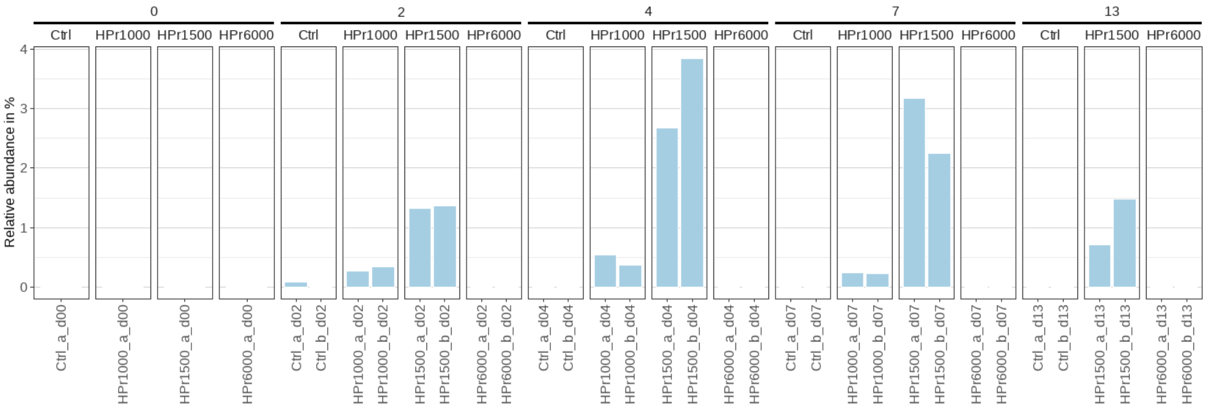


Figure S23. ASV 75. Bacteria, Firmicutes, Clostridia, Oscillospirales, Butyricicoccaceae, UCG-009, unknown species

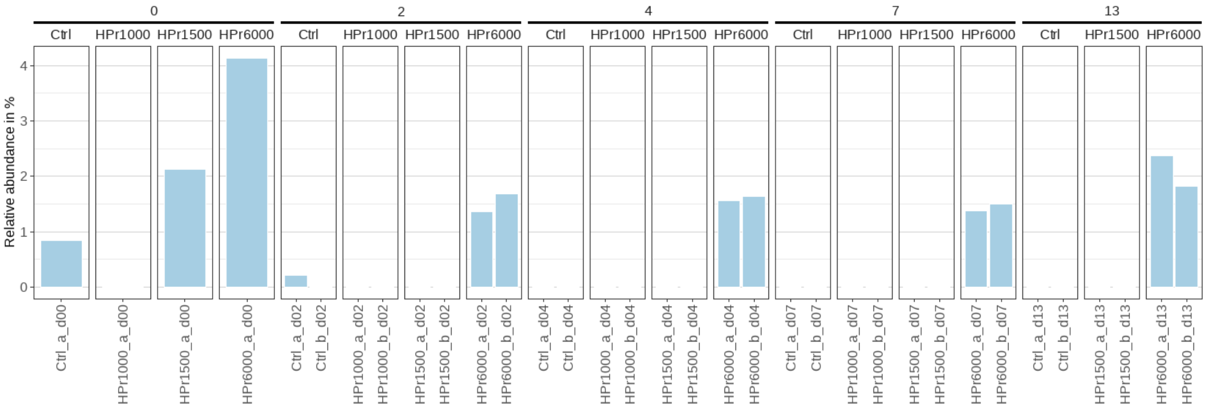


Figure S24. ASV 78. Bacteria, Proteobacteria, Gammaproteobacteria, Burkholderiales, Rhodocyclaceae, Zoogloea, Multi-affiliation

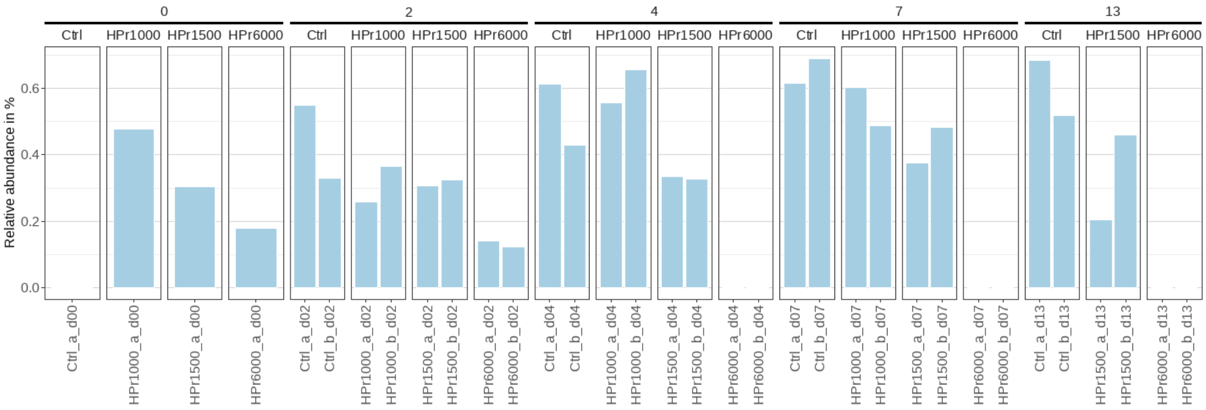


Figure S25. ASV 82. Bacteria, Synergistota, Synergistia, Synergistales, Synergistaceae, unknown genus, unknown species

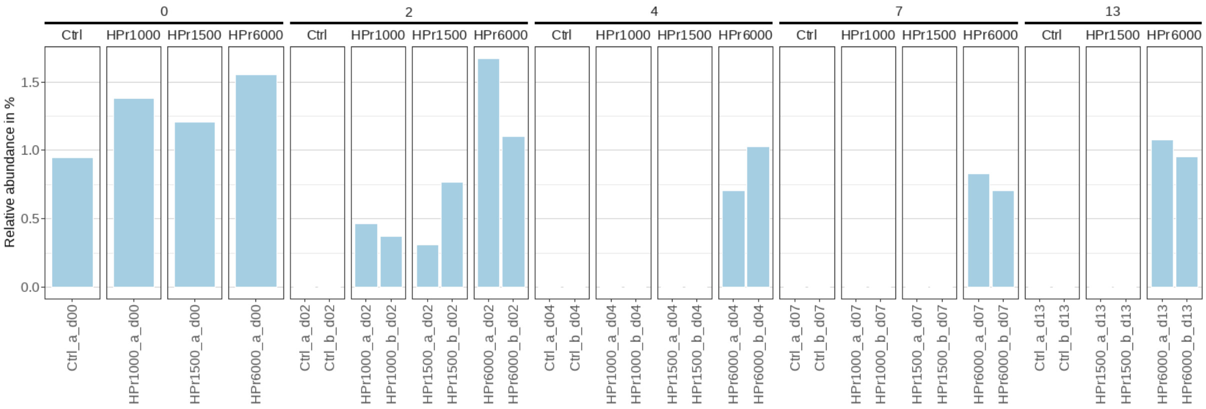


Figure S26. ASV 86. Bacteria, Campilobacterota, Campylobacteria, Campylobacterales, Arcobacteraceae, Arcobacter, Multi-affiliation

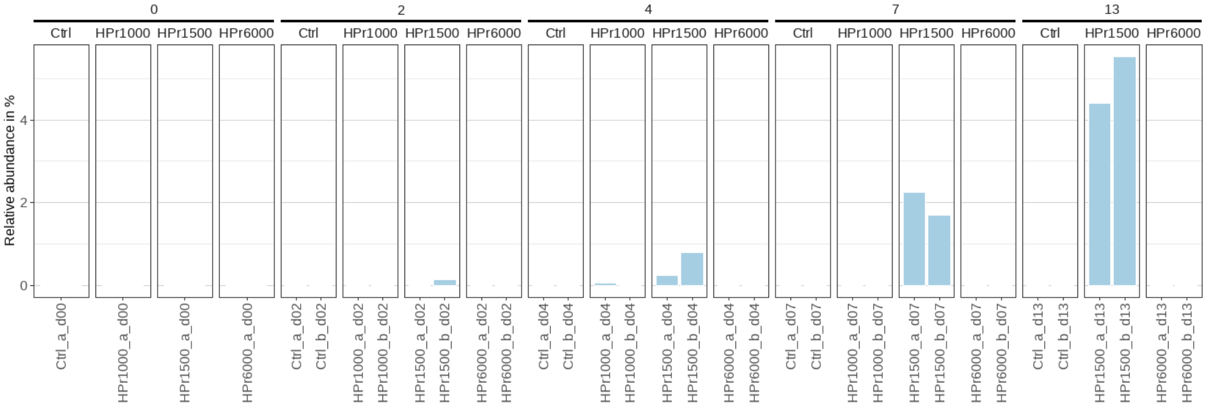


Figure S27. ASV 89. Bacteria, Proteobacteria, Gammaproteobacteria, Burkholderiales, Rhodocyclaceae, unknown genus, unknown species

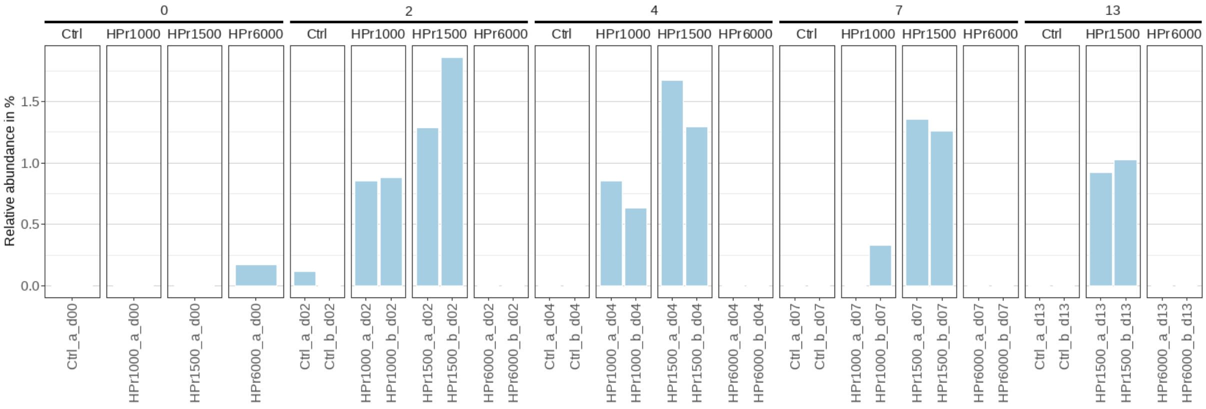


Figure S28. ASV 93. Bacteria, Firmicutes, Clostridia, Oscillospirales, Oscillospiraceae, UCG-002, unknown species

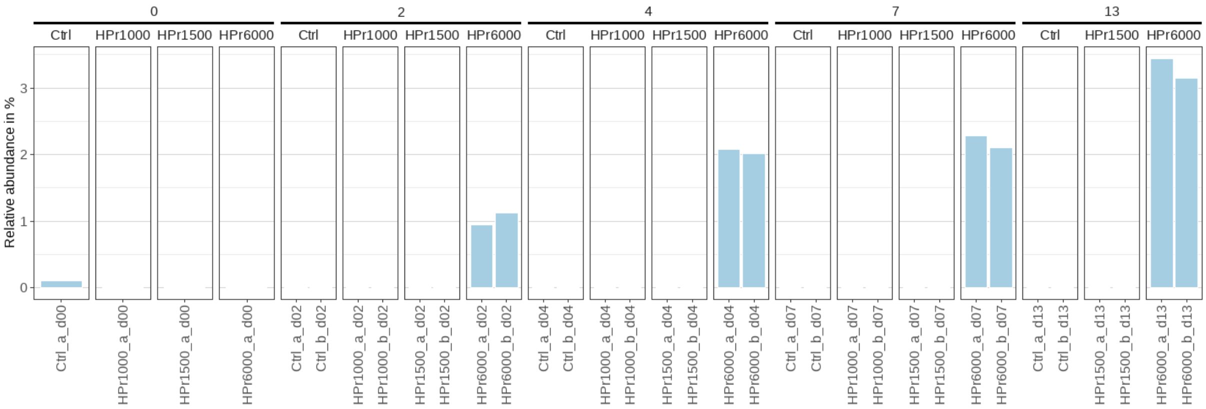


Figure S29. ASV 95. Bacteria, Firmicutes, Bacilli, Lactobacillales, Lactobacillaceae, Lactobacillus, Multi-affiliation

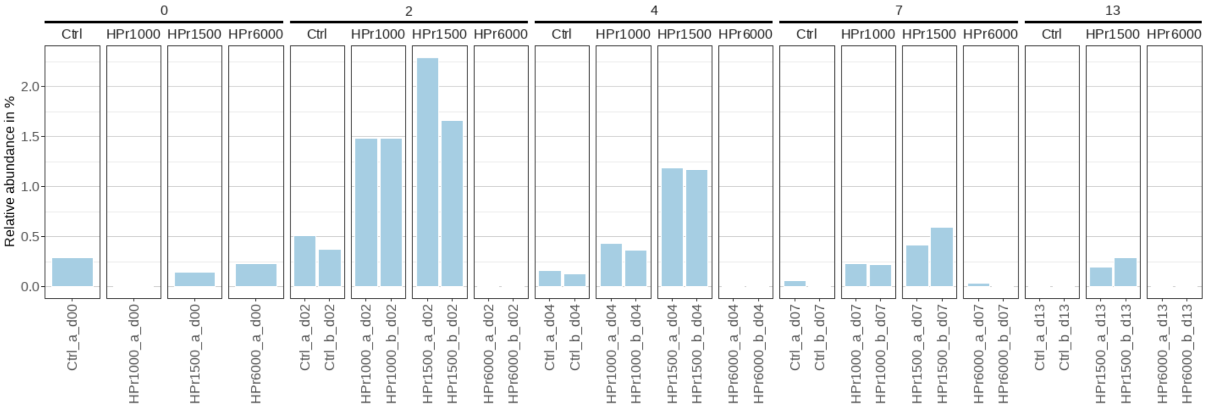


Figure S30. ASV 96. Bacteria, Firmicutes, Clostridia, Lachnospirales, Lachnospiraceae, XBB1006, Multi-affiliation

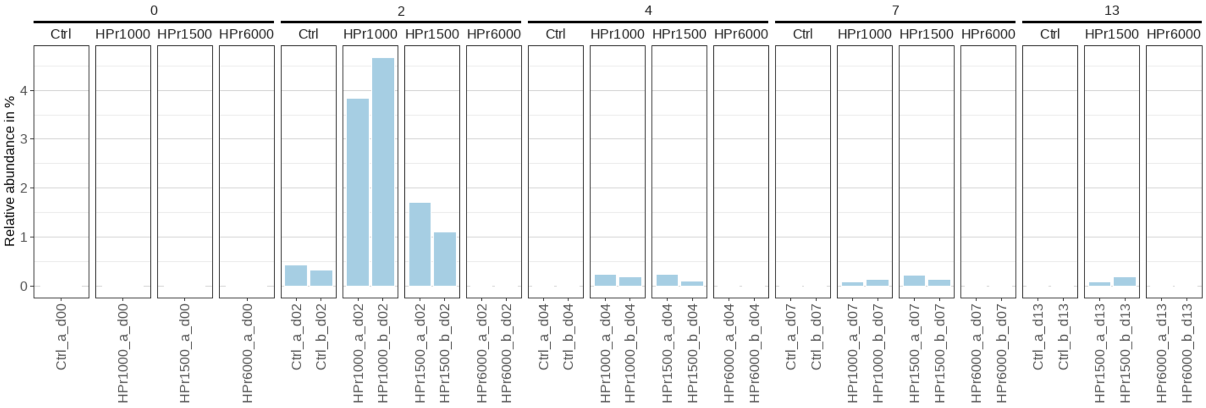


Figure S31. ASV 109. Bacteria, Bacteroidota, Bacteroidia, Bacteroidales, Marinifilaceae, unknown genus, unknown species

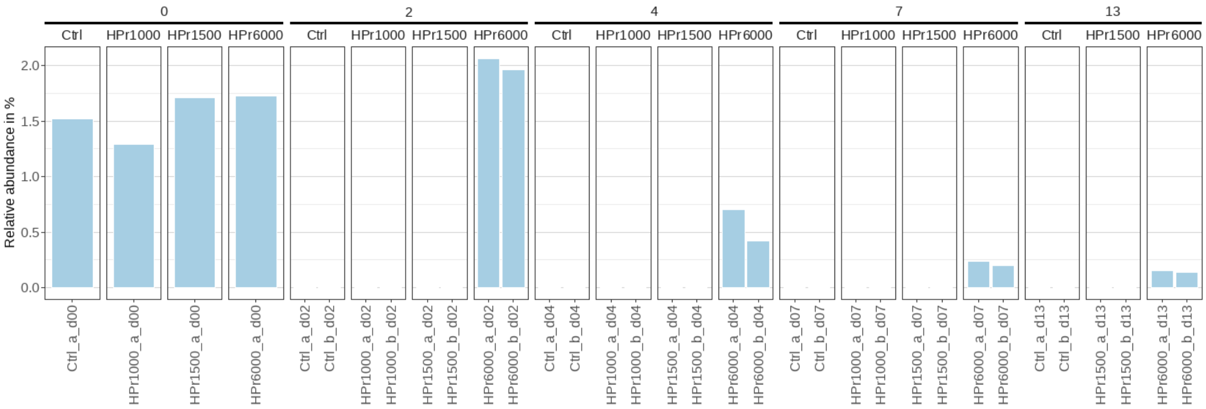


Figure S32. ASV 115. Bacteria, Myxococcota, Polyangia, Nannocystales, Nannocystaceae, Nannocystis, unknown species

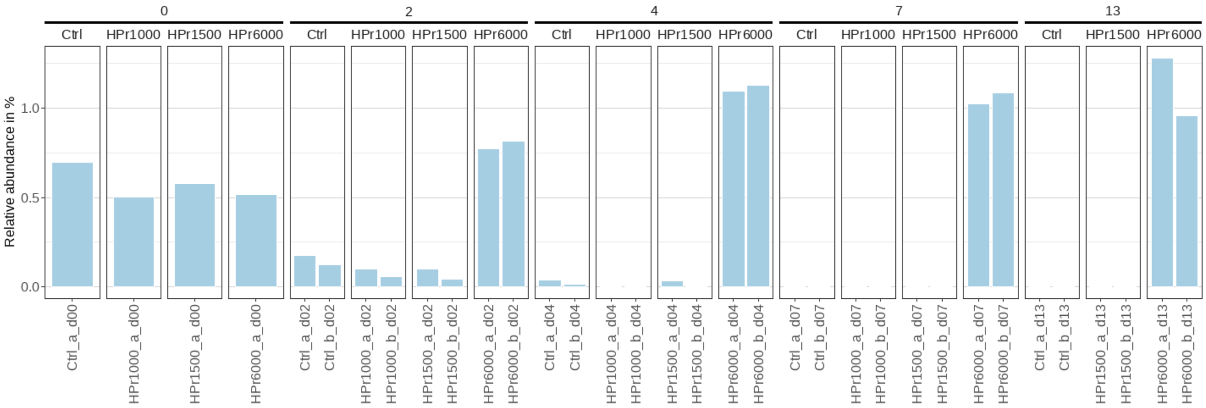


Figure S33. ASV 124. Bacteria, Firmicutes, Bacilli, Lactobacillales, Carnobacteriaceae, Trichococcus, Multi-affiliation

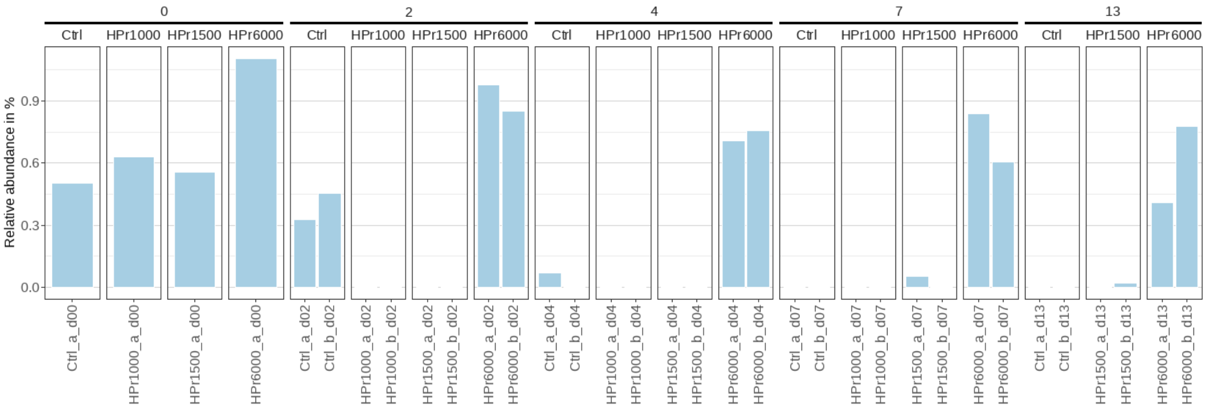


Figure S34. ASV 138. Bacteria, Bacteroidota, Bacteroidia, Bacteroidales, Paludibacteraceae, Paludibacter, unknown species

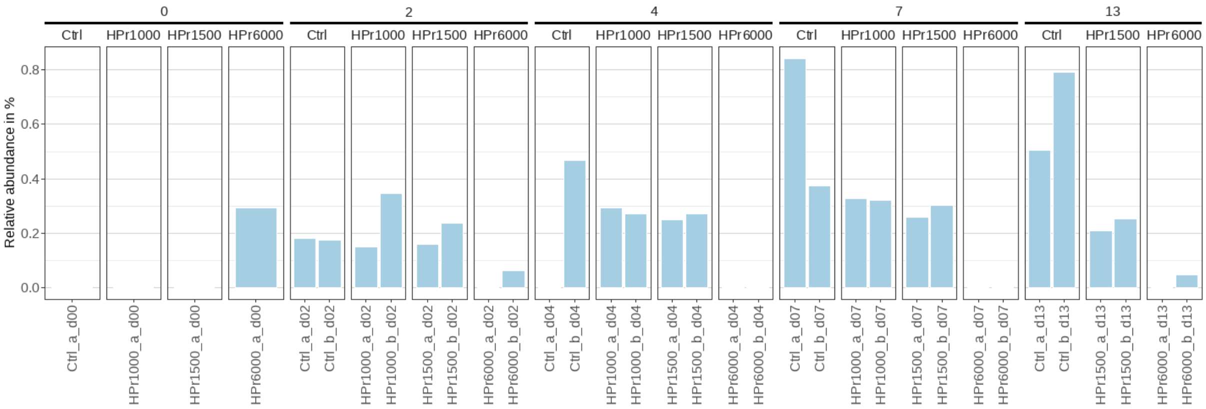


Figure S35. ASV 140. Archaea, Halobacterota, Methanomicrobia, Methanomicrobiales, Methanospirillaceae, Methanospirillum, Multi-affiliation

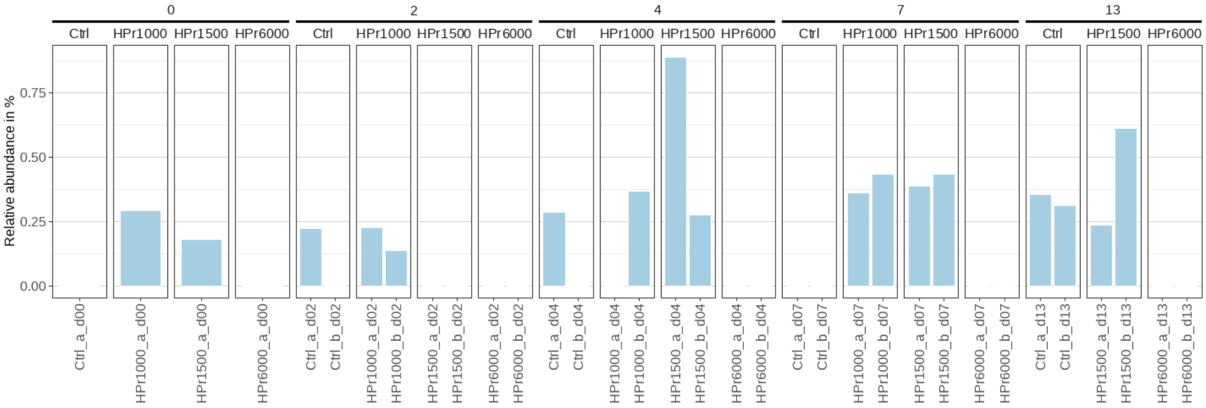


Figure S36. ASV 168. Bacteria, Proteobacteria, Gammaproteobacteria, Burkholderiales, Comamonadaceae, Brachymonas, Multi-affiliation

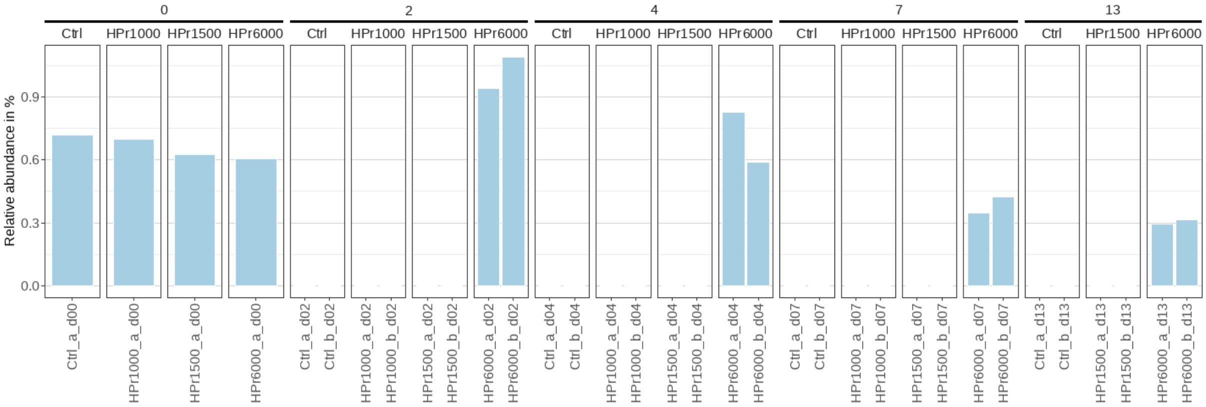


Figure S37. ASV 169. Bacteria, Bacteroidota, Bacteroidia, Chitinophagales, Chitinophagaceae, Aurantisolimonas, unknown species

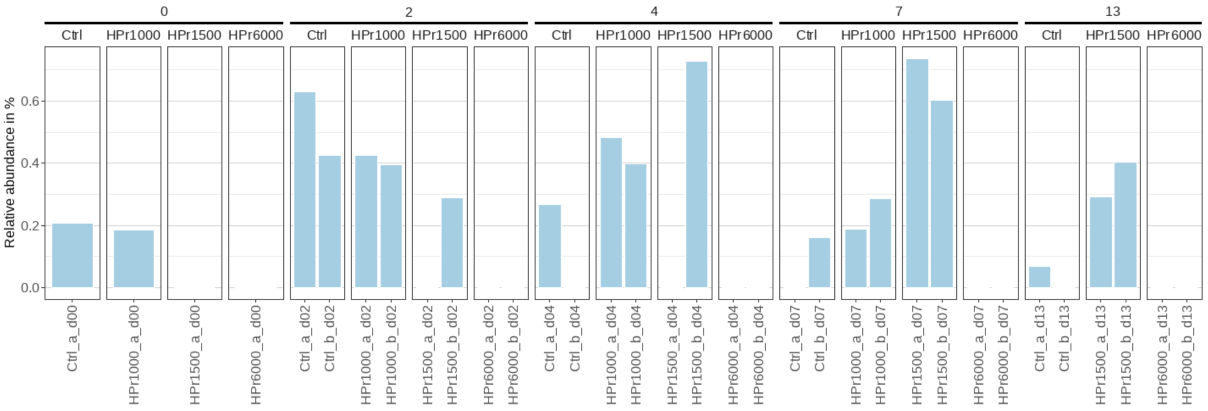


Figure S38. ASV 177. Bacteria, Firmicutes, Negativicutes, Acidaminococcales, Acidaminococcaceae, unknown genus, unknown species

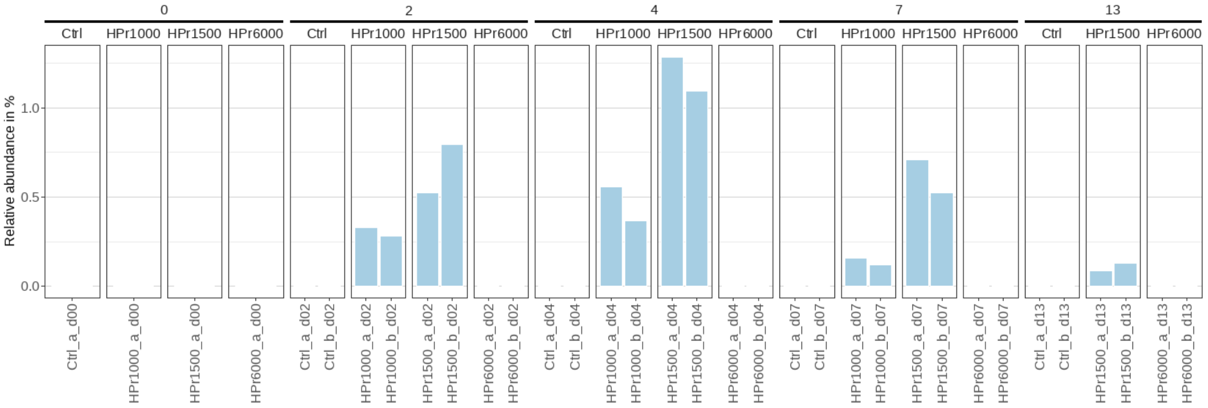


Figure S39. ASV 182. Bacteria, Firmicutes, Clostridia, Oscillospirales, Oscillospiraceae, NK4A214 group, unknown species

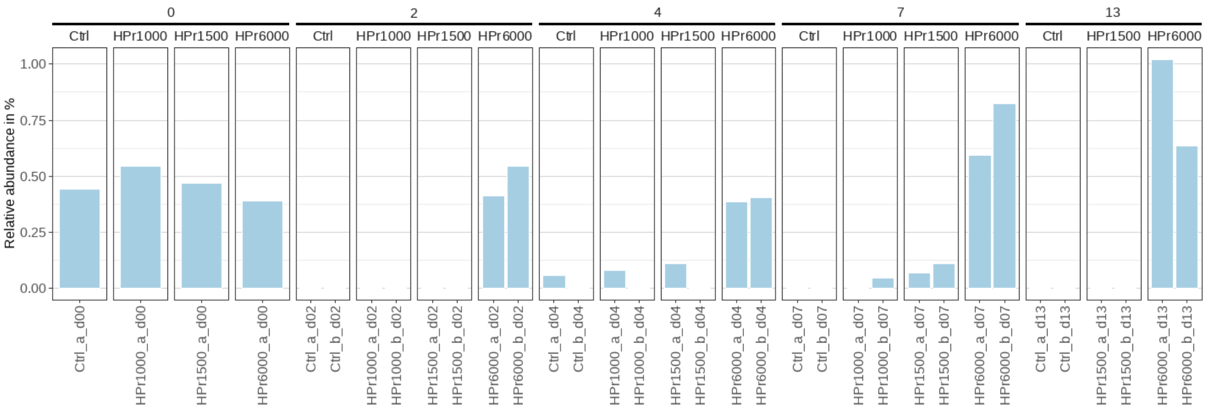


Figure S40. ASV 184. Bacteria, Proteobacteria, Gammaproteobacteria, Xanthomonadales, Rhodanobacteraceae, Dokdonella, unknown species

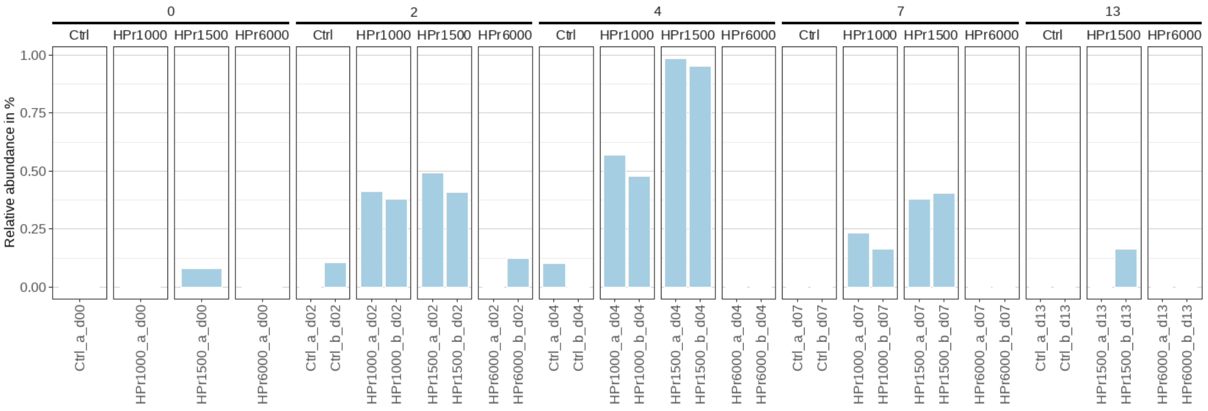


Figure S41. ASV 186. Bacteria, Firmicutes, Clostridia, Oscillospirales, Oscillospiraceae, NK4A214 group, unknown species

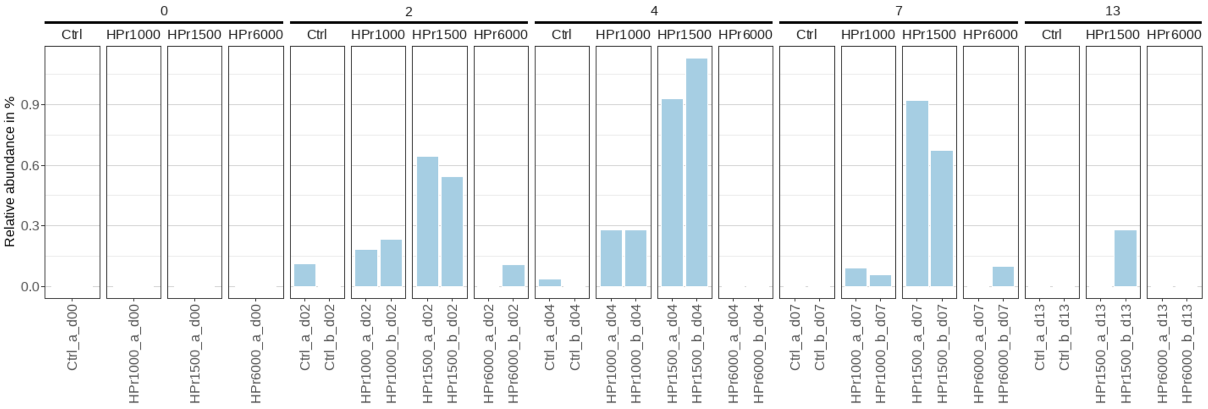


Figure S42. ASV 191. Bacteria, Firmicutes, Clostridia, Oscillospirales, Oscillospiraceae, UCG-005, unknown species

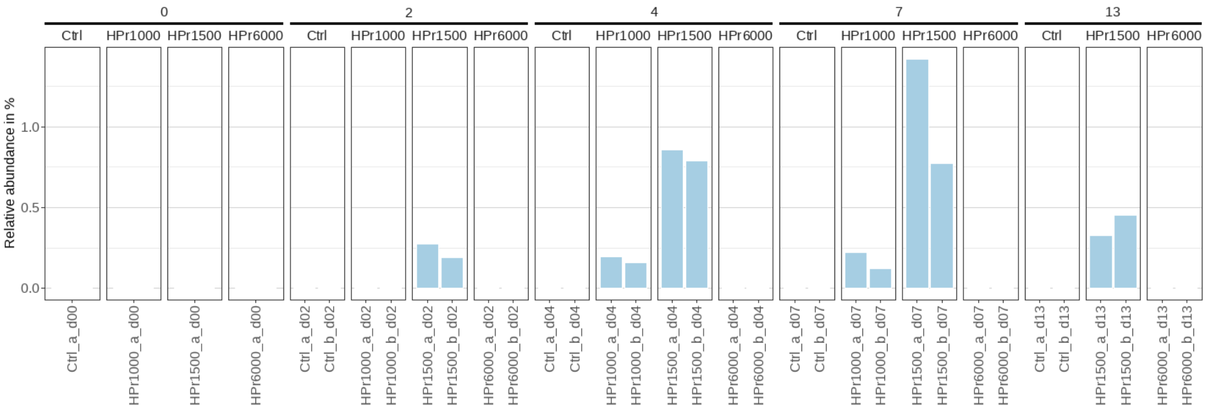


Figure S43. ASV 203. Bacteria, Firmicutes, Clostridia, Oscillospirales, Oscillospiraceae, NK4A214 group, unknown species

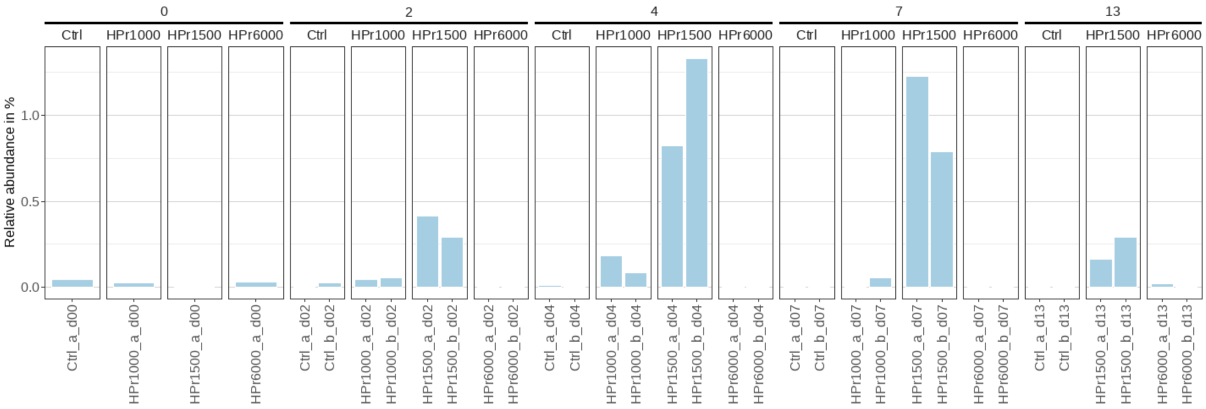


Figure S44. ASV 204. Bacteria, Bacteroidota, Bacteroidia, Bacteroidales, Tannerellaceae, Parabacteroides, unknown species

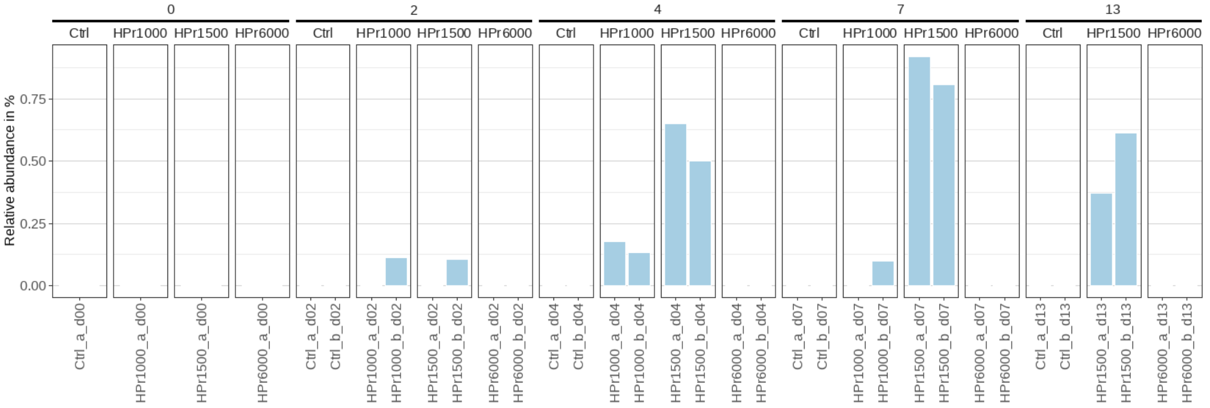


Figure S45. ASV 243. Bacteria, Firmicutes, Clostridia, Oscillospirales, Oscillospiraceae, NK4A214 group, unknown species

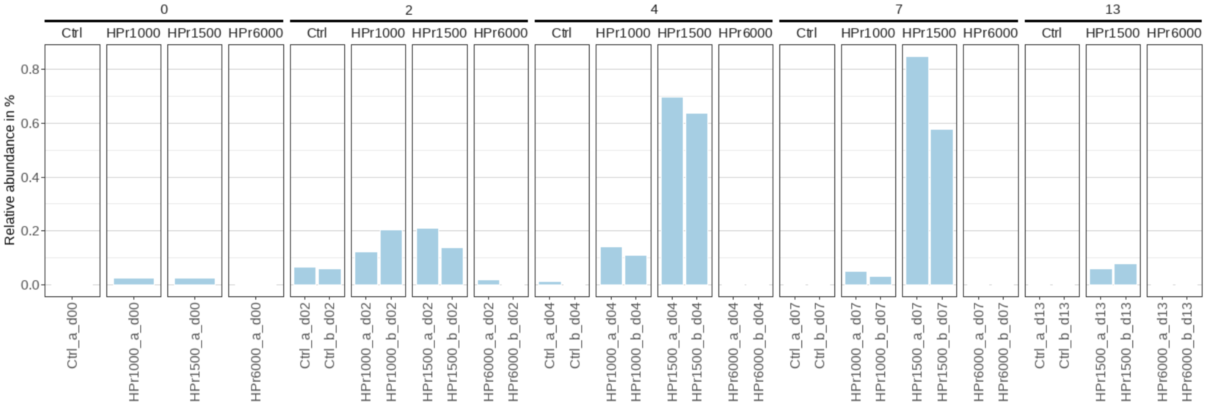


Figure S46. ASV 255. Bacteria, Bacteroidota, Bacteroidia, Bacteroidales, Prevotellaceae, Prevotella, unknown species

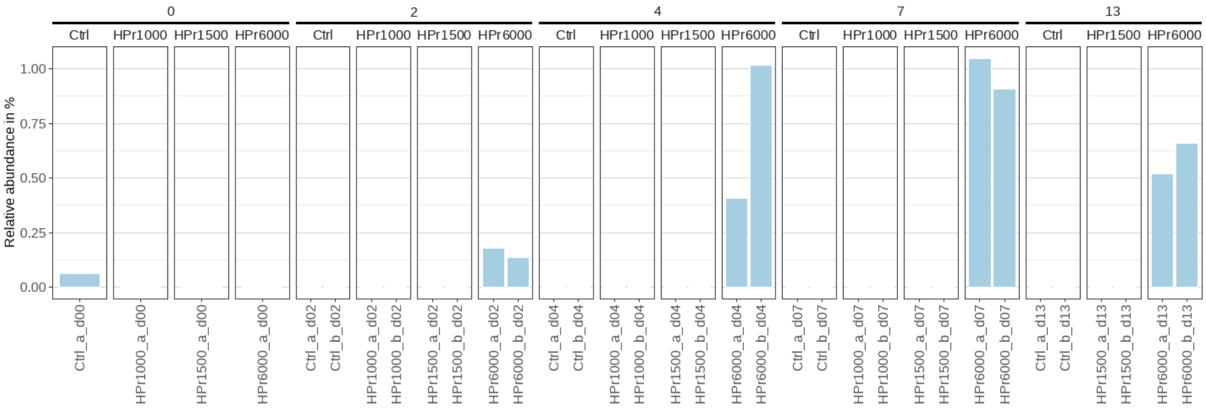


Figure S47. ASV 262. Bacteria, Firmicutes, Negativicutes, Veillonellales-Selenomonadales, Selenomonadaceae, Selenomonas, Multi-affiliation

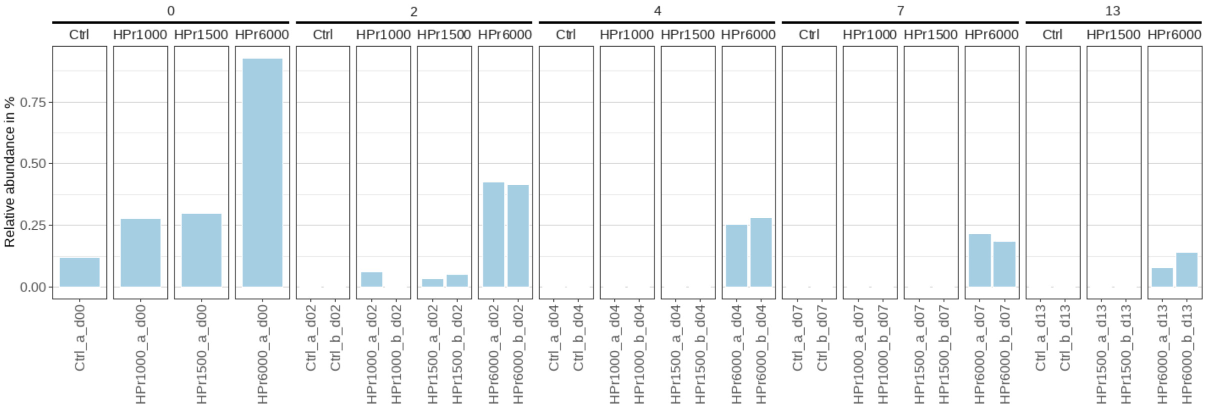


Figure S48. ASV 275. Bacteria, Bacteroidota, Bacteroidia, Bacteroidales, Paludibacteraceae, unknown genus, metagenome

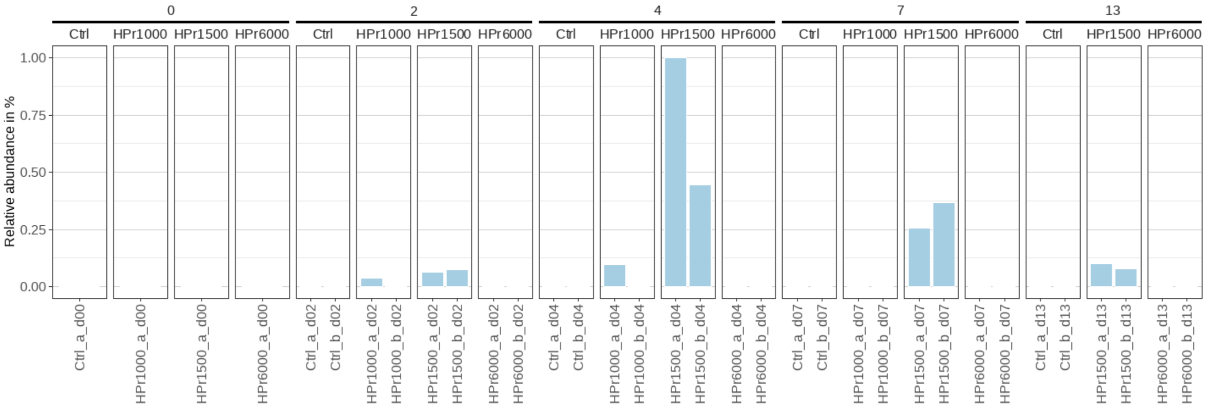


Figure S49. ASV 386. Bacteria, Spirochaetota, Spirochaetia, Spirochaetales, Spirochaetaceae, Treponema, Treponema sp.

## CC3

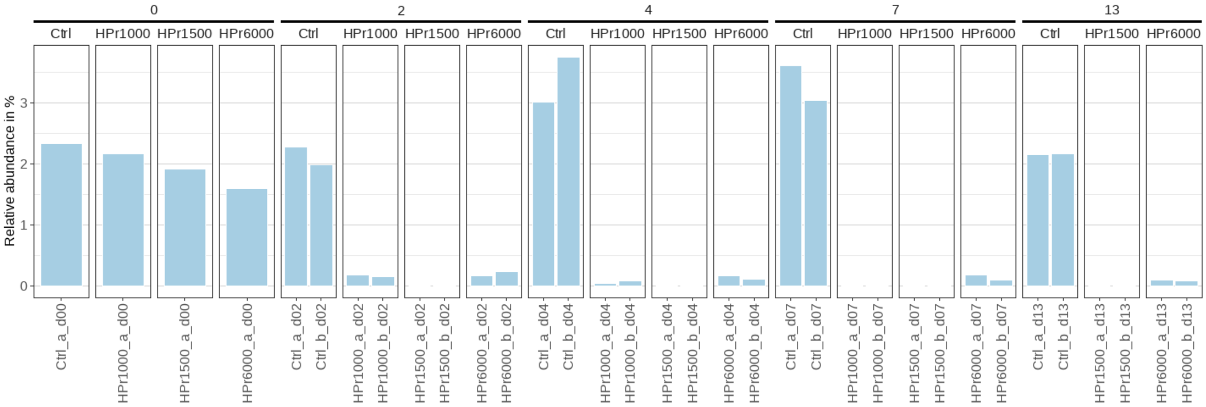


Figure S50. ASV 17. Bacteria, Bacteroidota, Bacteroidia, Sphingobacteriales, ST-12K33, unknown genus, metagenome

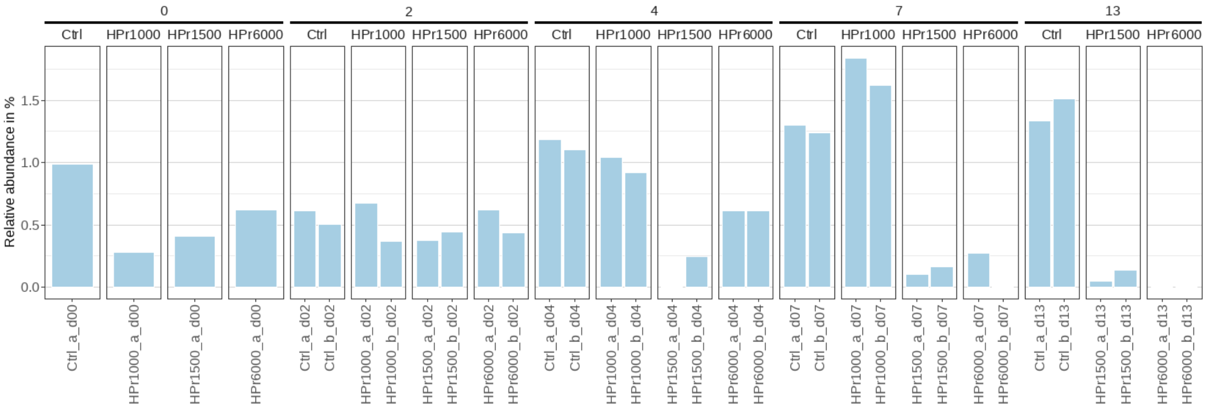


Figure S51. ASV 31. Bacteria, Firmicutes, Clostridia, Peptostreptococcales-Tissierellales, Sedimentibacteraceae, Sedimentibacter, unknown species

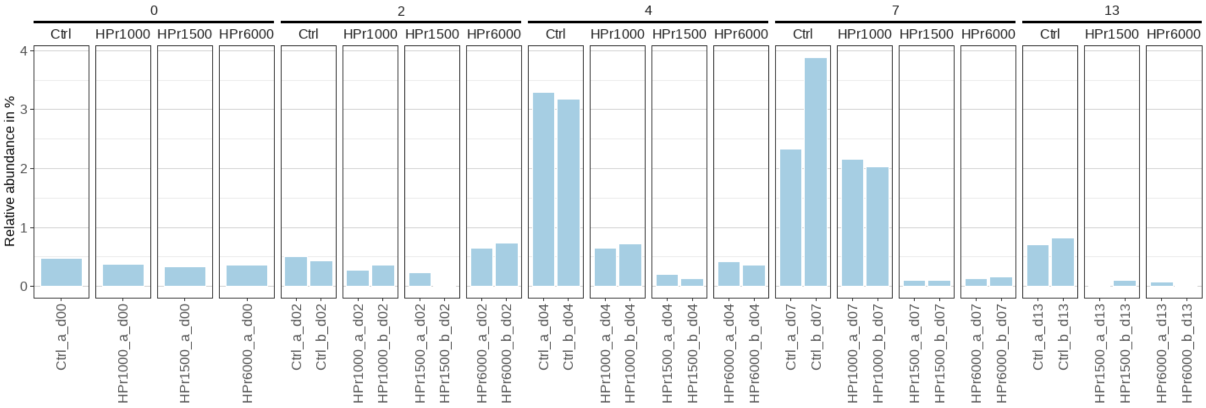


Figure S52. ASV 38. Bacteria, Firmicutes, Syntrophomonadia, Syntrophomonadales, Syntrophomonadaceae, Syntrophomonas, unknown species

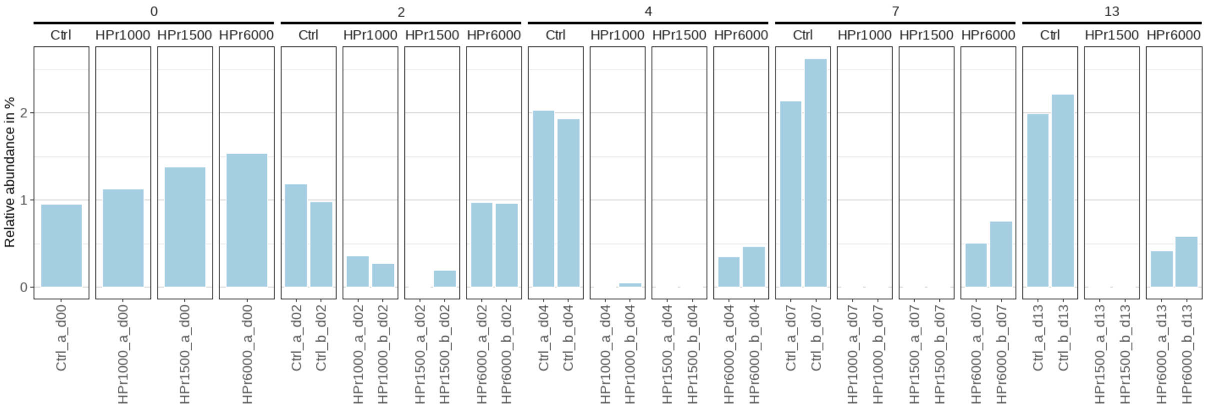


Figure S53. ASV 39. Bacteria, Desulfobacterota, Syntrophia, Syntrophales, Smithellaceae, Smithella, Multi-affiliation

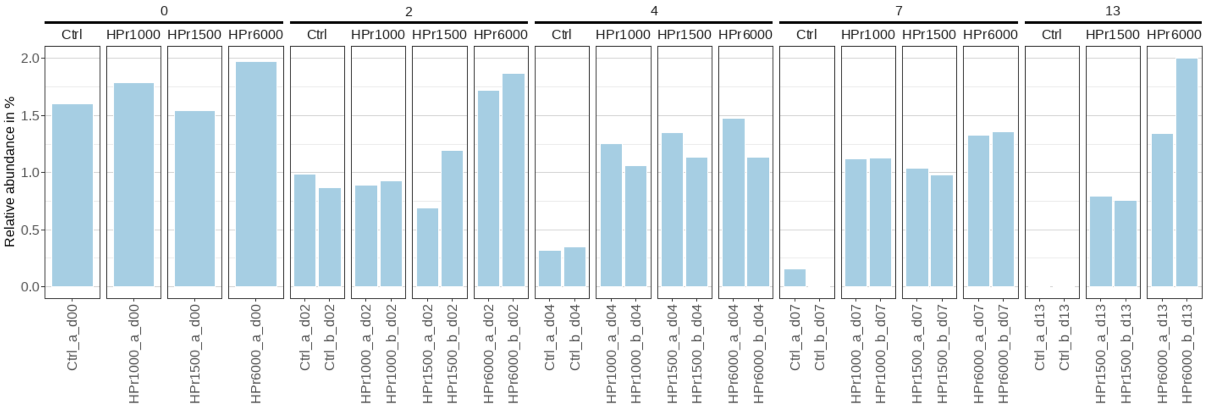


Figure S54. ASV 43. Bacteria, Proteobacteria, Gammaproteobacteria, Burkholderiales, Methylophilaceae, unknown genus, unknown species

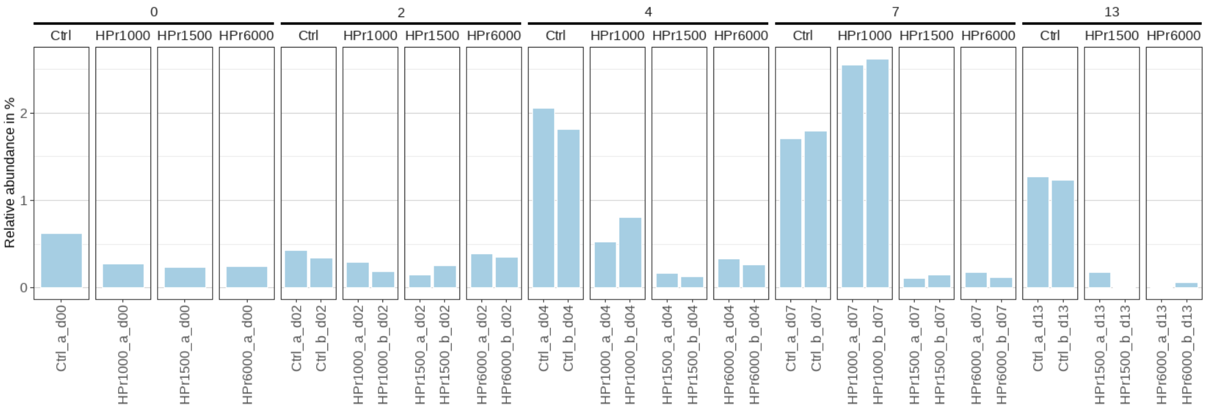


Figure S55. ASV 46. Bacteria, Firmicutes, Syntrophomonadia, Syntrophomonadales, Syntrophomonadaceae, Syntrophomonas, unknown species

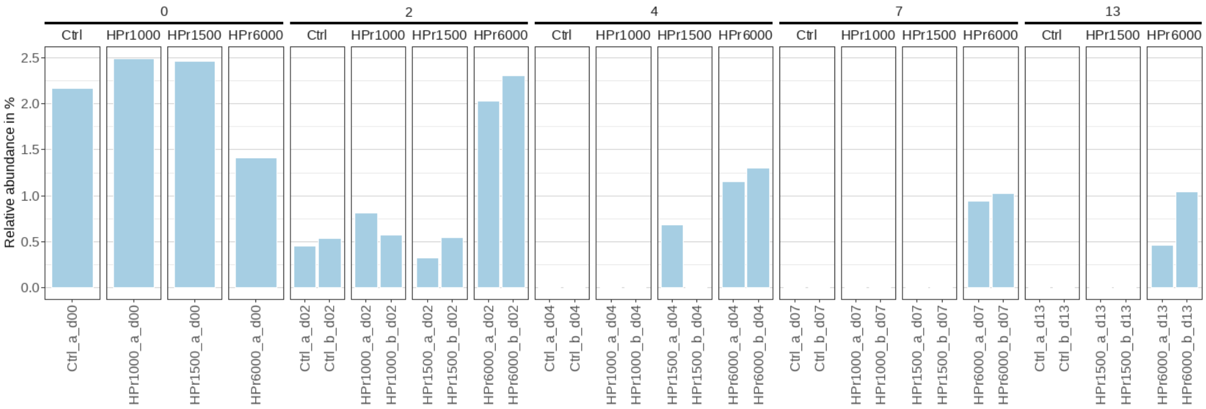


Figure S56. ASV 58. Bacteria, Proteobacteria, Gammaproteobacteria, Burkholderiales, Comamonadaceae, Hydrogenophaga, unknown species

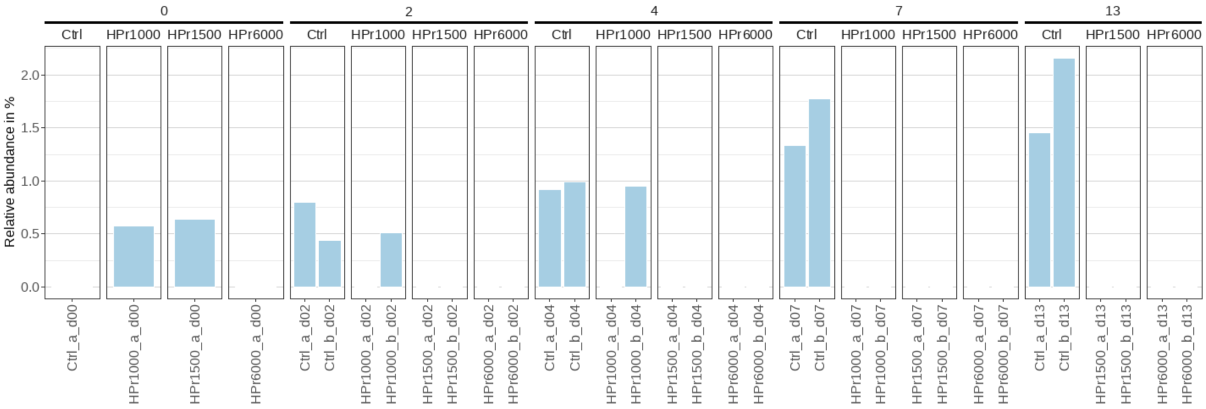


Figure S57. ASV 61. Bacteria, Firmicutes, Clostridia, Peptostreptococcales-Tissierellales, Sedimentibacteraceae, Sedimentibacter, unknown species

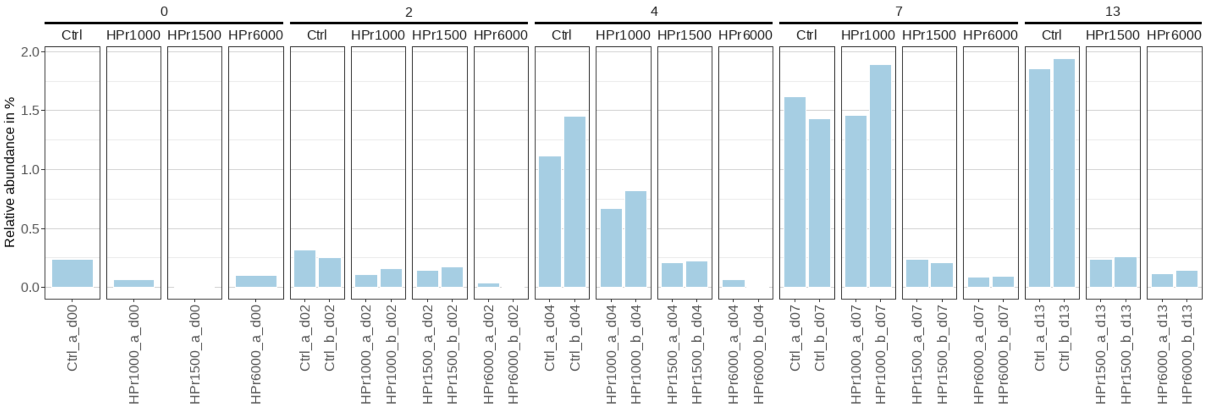


Figure S58. ASV 62. Bacteria, Firmicutes, Syntrophomonadia, Syntrophomonadales, Syntrophomonadaceae, Syntrophomonas, unknown species

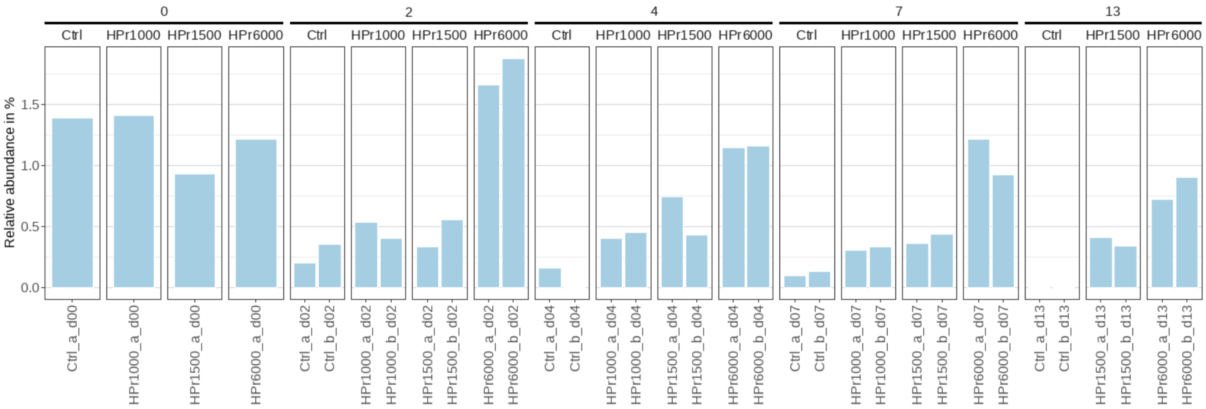


Figure S59. ASV 64. Bacteria, Proteobacteria, Gammaproteobacteria, Burkholderiales, Comamonadaceae, Multi-affiliation, Multi-affiliation

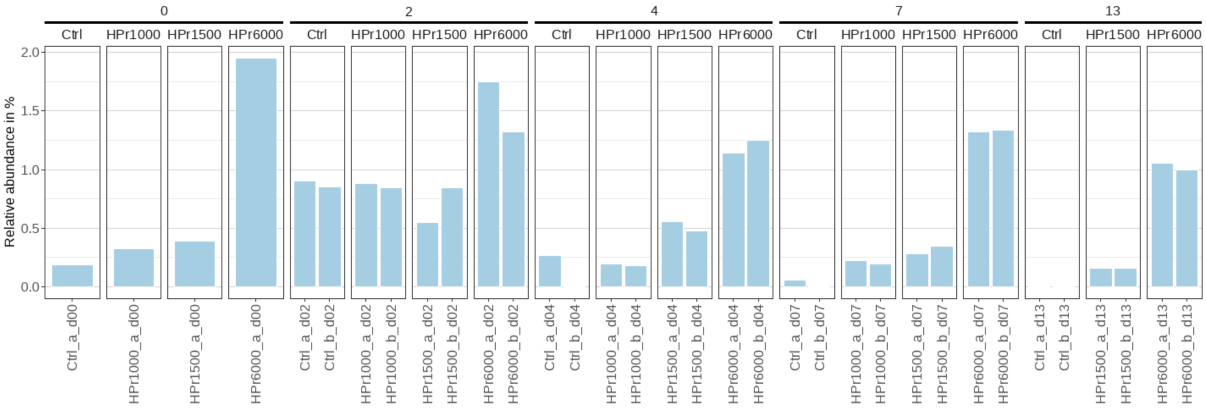


Figure S60. ASV 70. Bacteria, Bacteroidota, Bacteroidia, Bacteroidales, Paludibacteraceae, unknown genus, unknown species

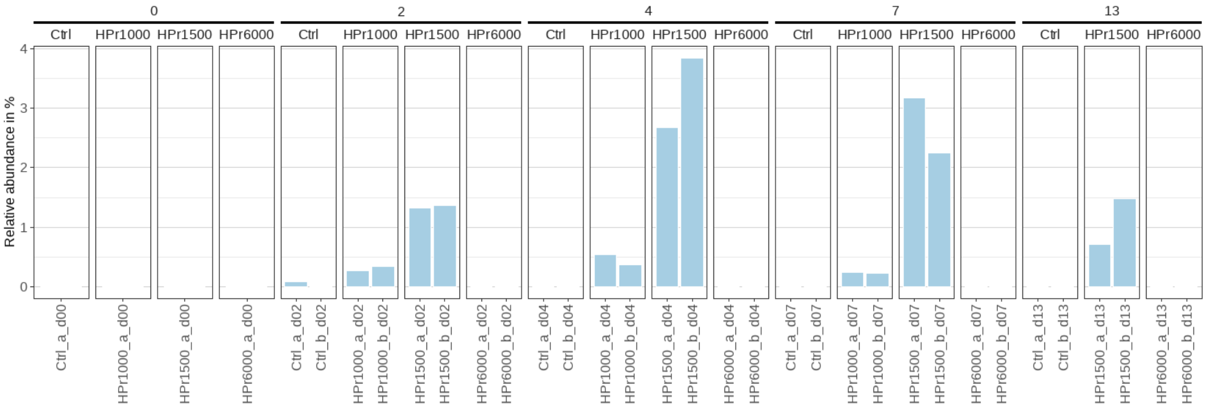


Figure S61. ASV 75. Bacteria, Firmicutes, Clostridia, Oscillospirales, Butyricicoccaceae, UCG-009, unknown species

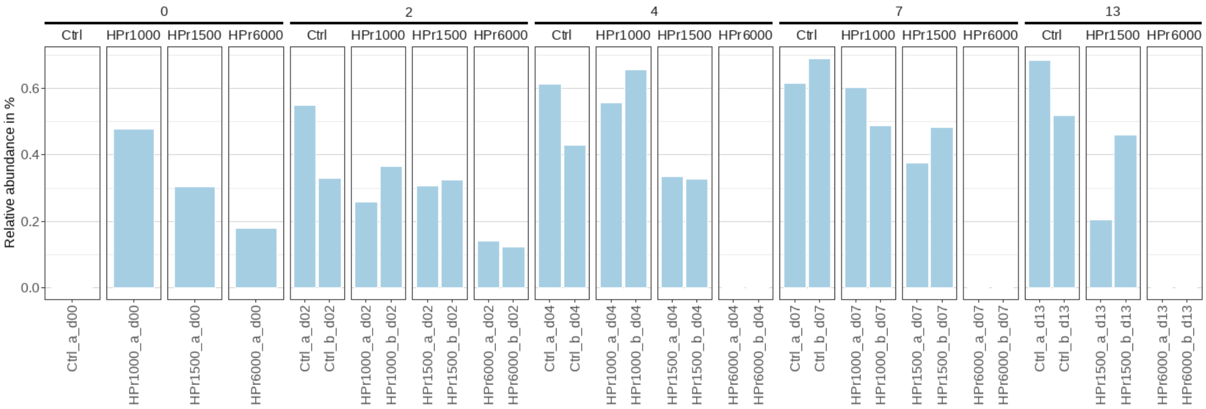


Figure S62. ASV 82. Bacteria, Synergistota, Synergistia, Synergistales, Synergistaceae, unknown genus, unknown species

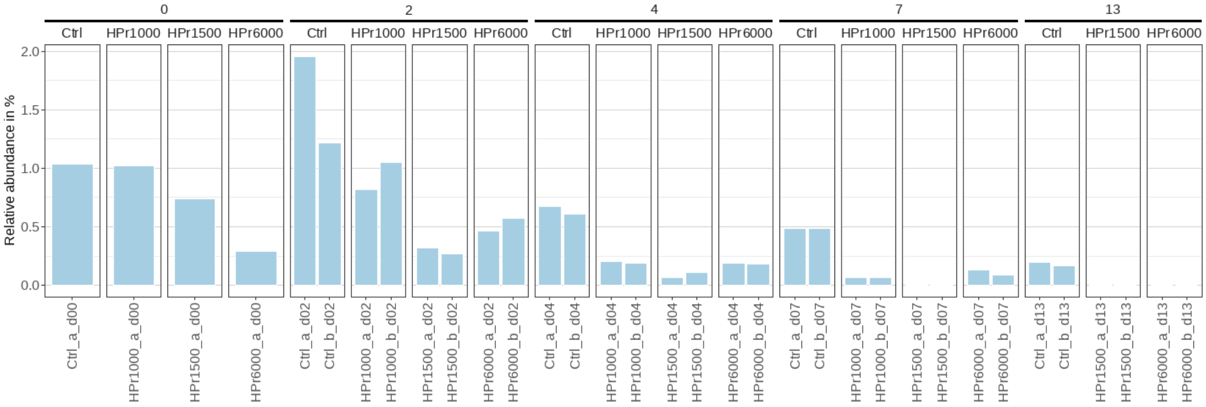


Figure S63. ASV 83. Bacteria, Firmicutes, Clostridia, Peptostreptococcales-Tissierellales, Peptostreptococcaceae, Acetoanaerobium, Multi-affiliation

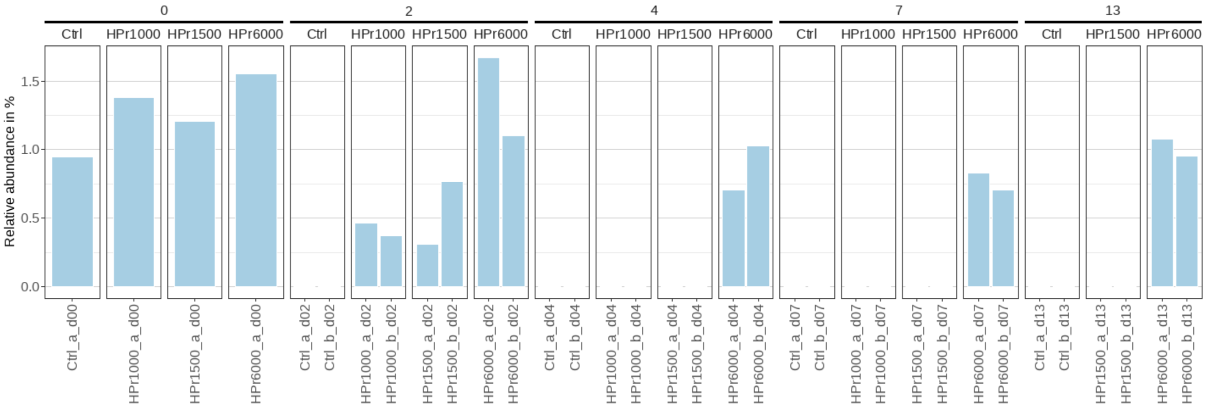


Figure S64. ASV 86. Bacteria, Campilobacterota, Campylobacteria, Campylobacterales, Arcobacteraceae, Arcobacter, Multi-affiliation

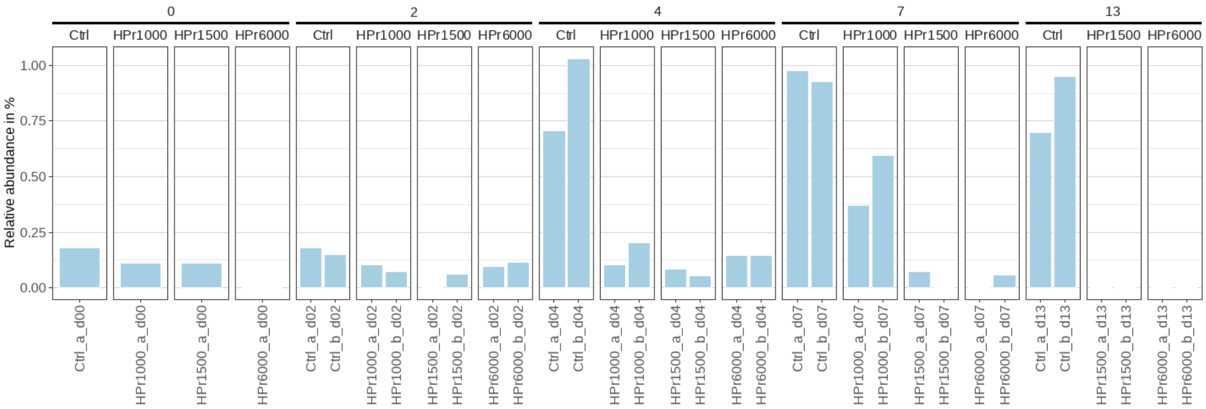


Figure S65. ASV 88. Bacteria, Firmicutes, Syntrophomonadia, Syntrophomonadales, Syntrophomonadaceae, Syntrophomonas, metagenome

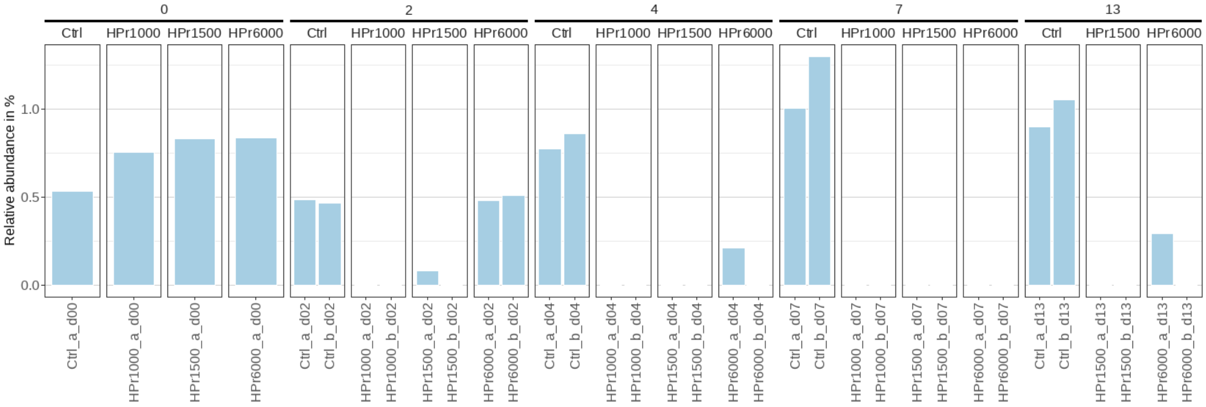


Figure S66. ASV 120. Bacteria, Desulfobacterota, Syntrophia, Syntrophales, Smithellaceae, Smithella, Multi-affiliation

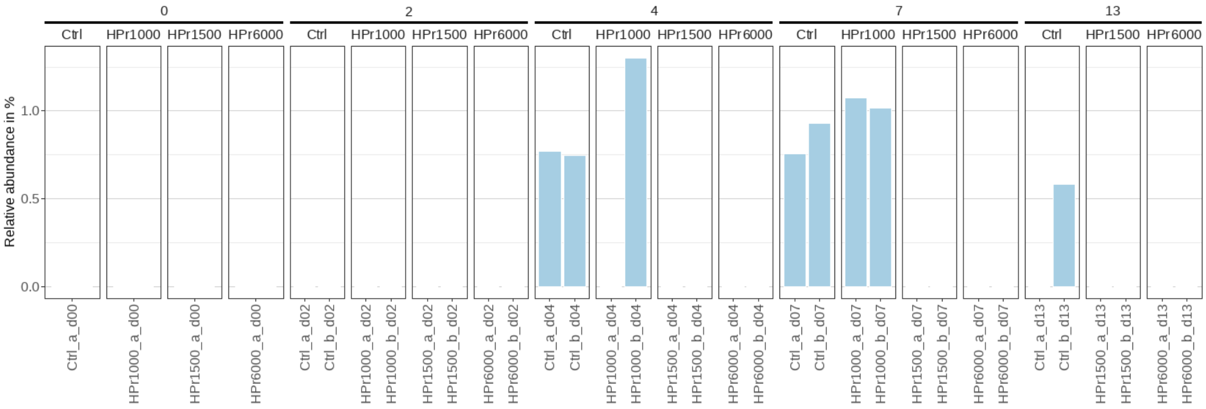


Figure S67. ASV 152. Bacteria, Spirochaetota, Spirochaetia, Spirochaetales, Spirochaetaceae, unknown genus, unknown species

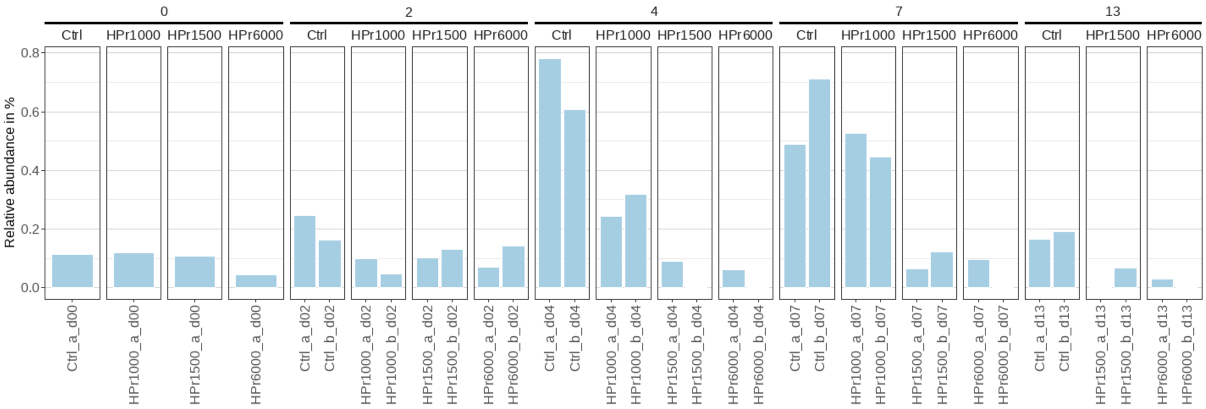


Figure S68. ASV 176. Bacteria, Firmicutes, Syntrophomonadia, Syntrophomonadales, Syntrophomonadaceae, Syntrophomonas, Syntrophomonas wolfei

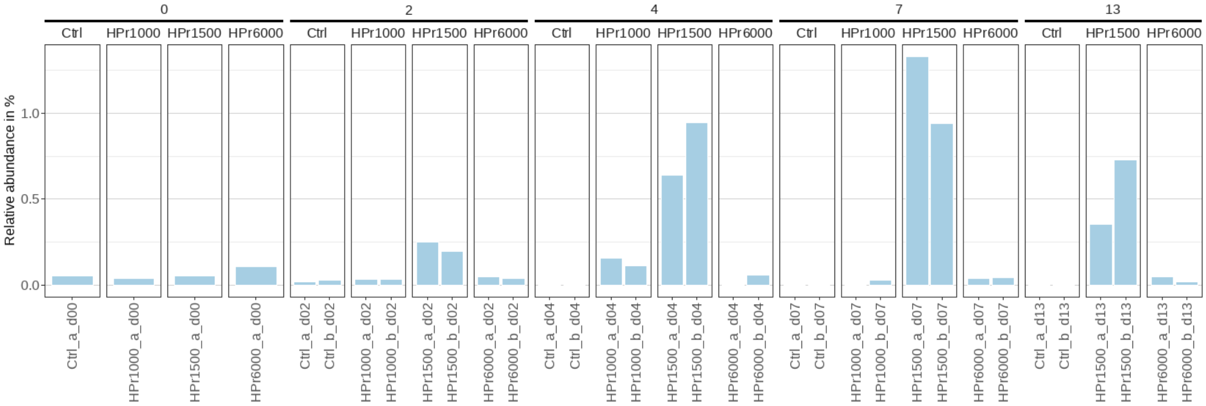


Figure S69. ASV 193. Bacteria, Bacteroidota, Bacteroidia, Bacteroidales, Tannerellaceae, Parabacteroides, unknown species

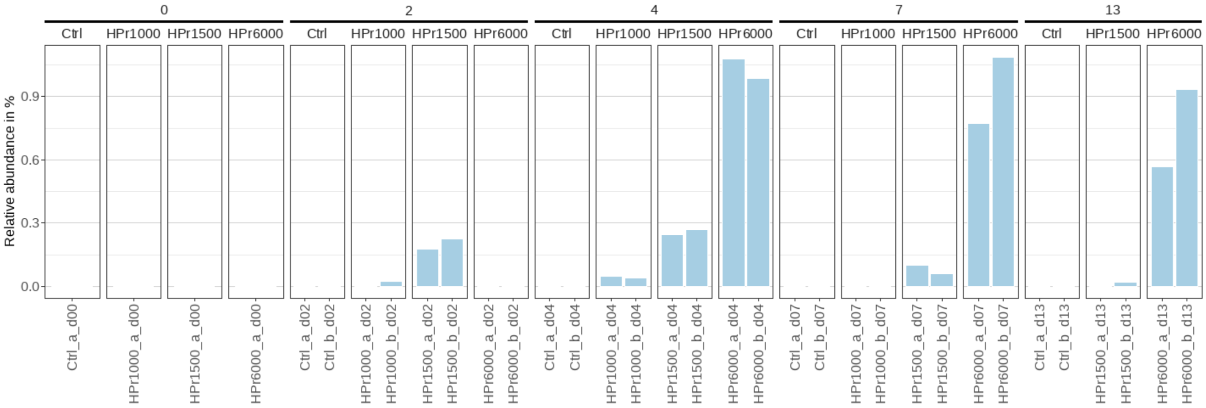


Figure S70. ASV 205. Bacteria, Firmicutes, Negativicutes, Veillonellales-Selenomonadales, Veillonellaceae, Megasphaera, unknown species

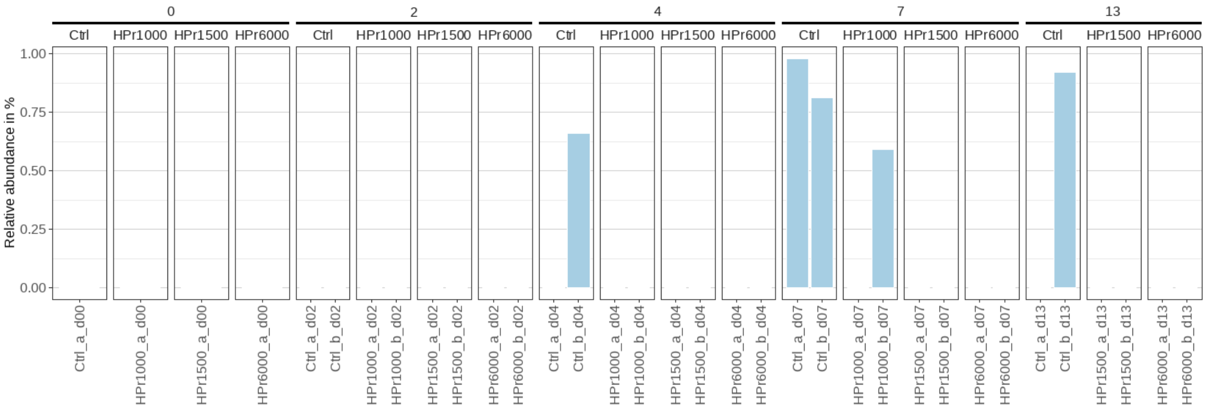


Figure S71. ASV 239. Bacteria, Firmicutes, Clostridia, Peptostreptococcales-Tissierellales, Sedimentibacteraceae, Sedimentibacter, unknown species

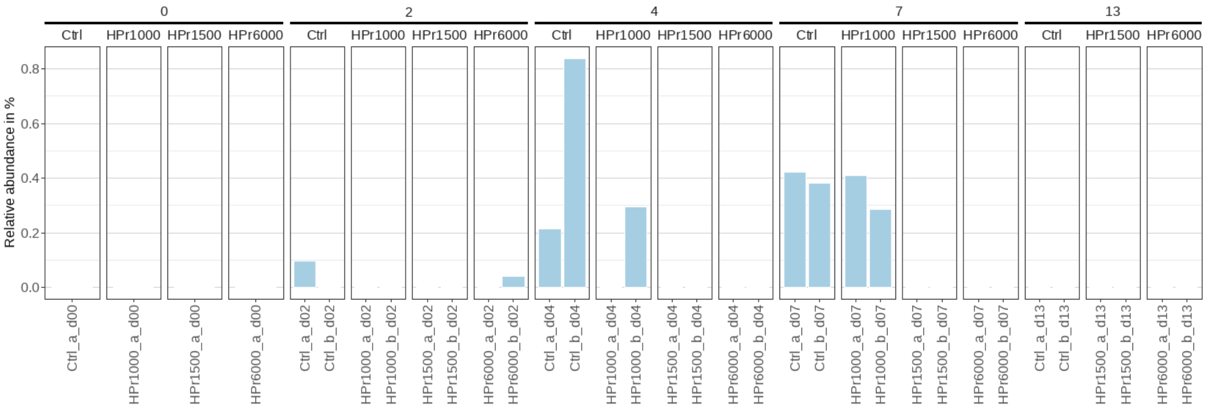


Figure S72. ASV 312. Bacteria, Bacteroidota, Bacteroidia, Sphingobacteriales, Lentimicrobiaceae, Lentimicrobium, iron-reducing bacterium enrichment culture clone HN105

## CC4

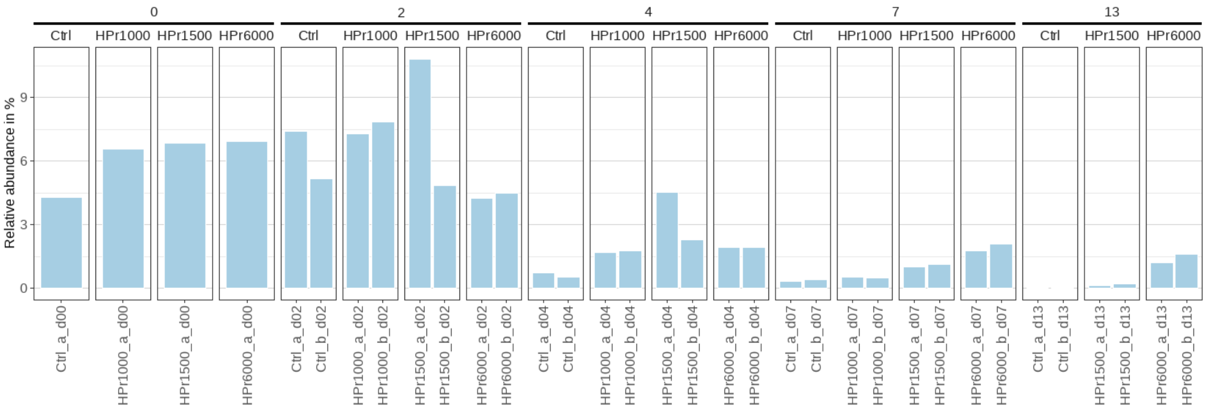


Figure S73. ASV 9. Bacteria, Bacteroidota, Bacteroidia, Bacteroidales, Bacteroidaceae, Bacteroides, Multi-affiliation

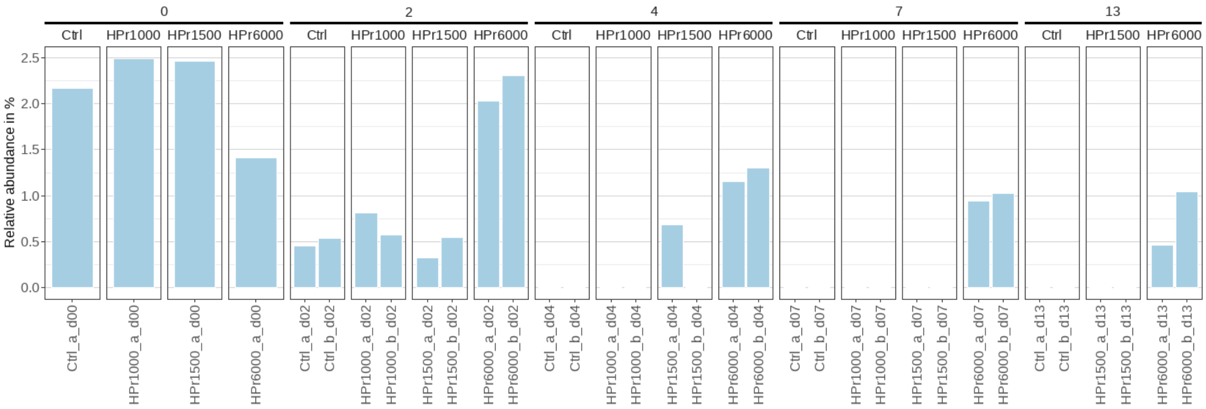


Figure S74. ASV 58. Bacteria, Proteobacteria, Gammaproteobacteria, Burkholderiales, Comamonadaceae, Hydrogenophaga, unknown species

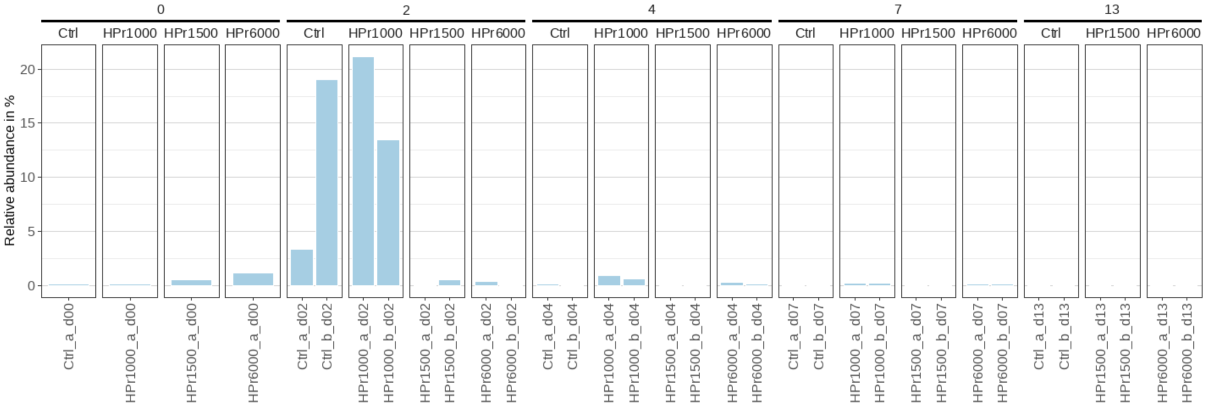


Figure S75. ASV 20. Bacteria, Bacteroidota, Bacteroidia, Bacteroidales, p-251-o5, unknown genus, unknown species

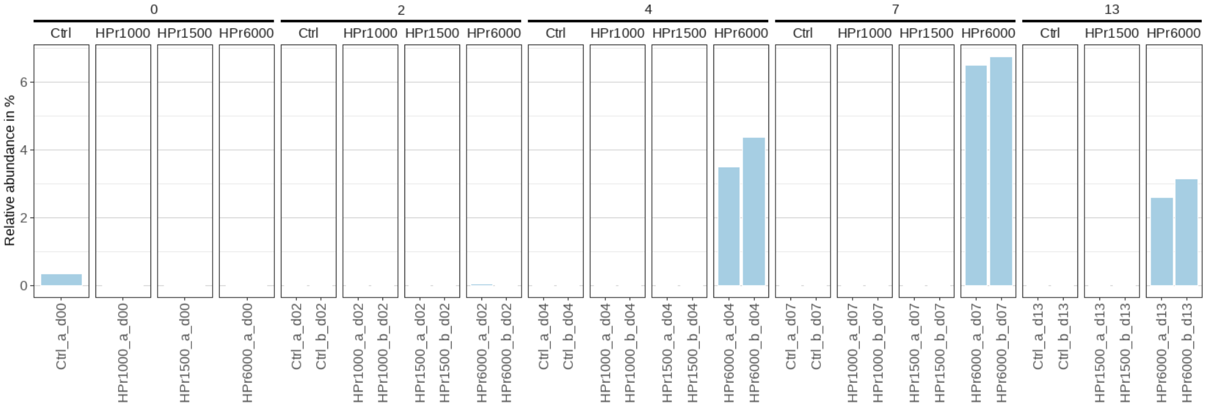


Figure S76. ASV 60. Bacteria, Bacteroidota, Bacteroidia, Bacteroidales, Prevotellaceae, Prevotella, unknown species

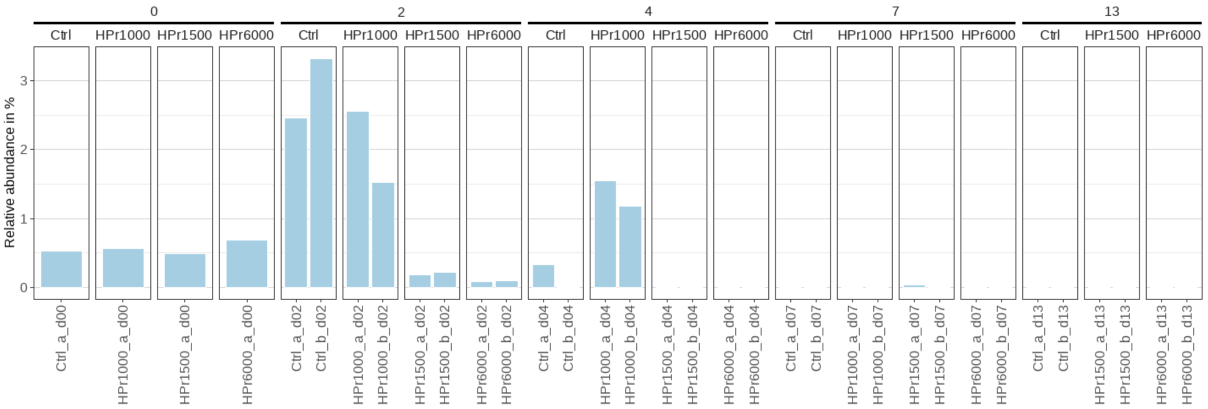


Figure S77. ASV 72. Bacteria, Firmicutes, Clostridia, unknown order, Hungateiclostridiaceae, Saccharofermentans, unknown species

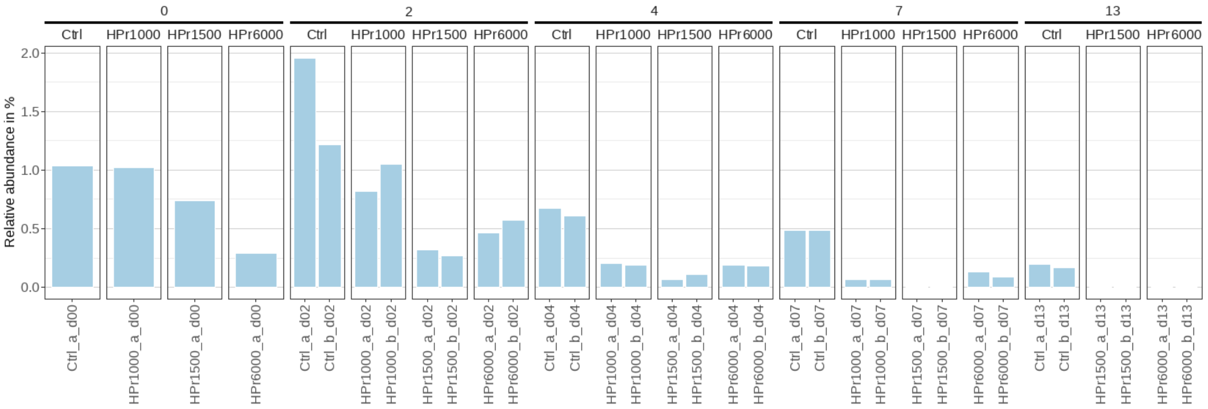


Figure S78. ASV 83. Bacteria, Firmicutes, Clostridia, Peptostreptococcales-Tissierellales, Peptostreptococcaceae, Acetoanaerobium, Multi-affiliation

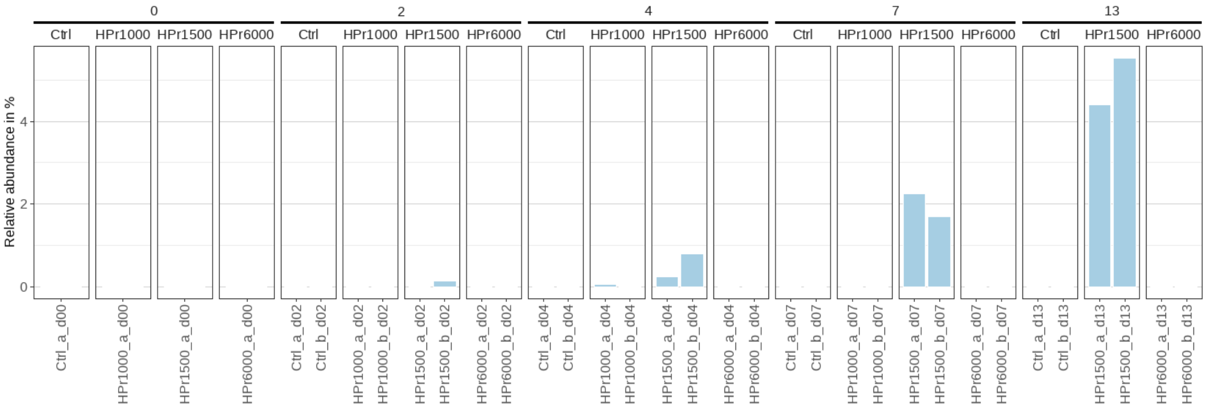


Figure S79. ASV 89. Bacteria, Proteobacteria, Gammaproteobacteria, Burkholderiales, Rhodocyclaceae, unknown genus, unknown species

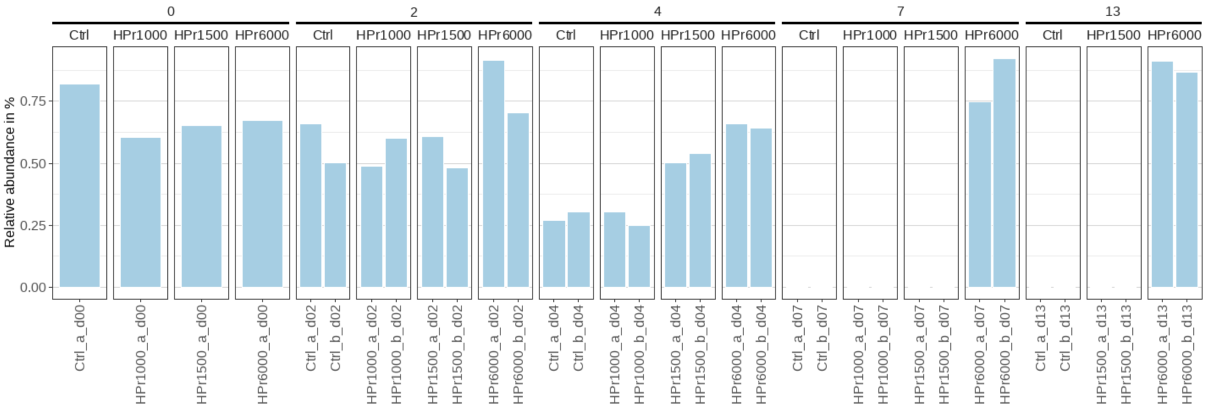


Figure S80. ASV 99. Bacteria, Bacteroidota, Bacteroidia, Bacteroidales, Tannerellaceae, Macellibacteroides, Multi-affiliation

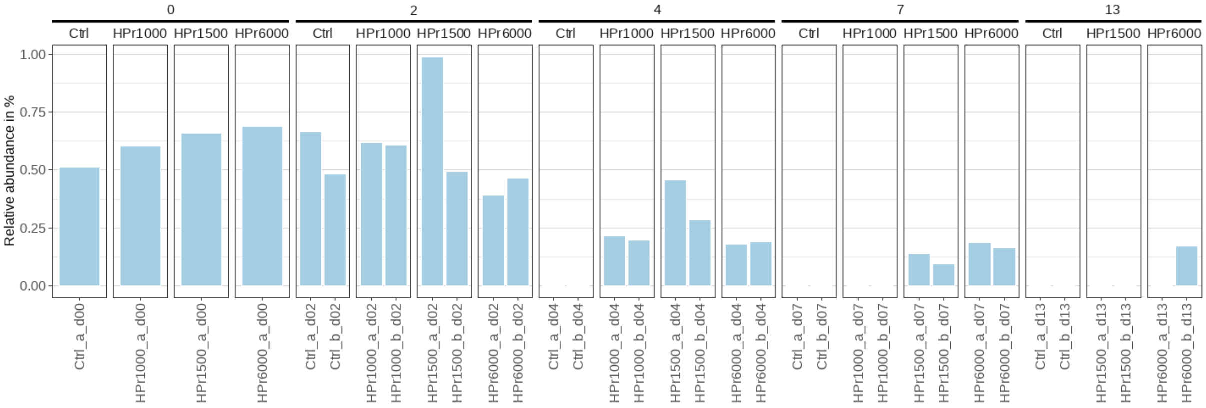


Figure S81. ASV 131. Bacteria, Bacteroidota, Bacteroidia, Bacteroidales, Bacteroidaceae, Bacteroides, Multi-affiliation

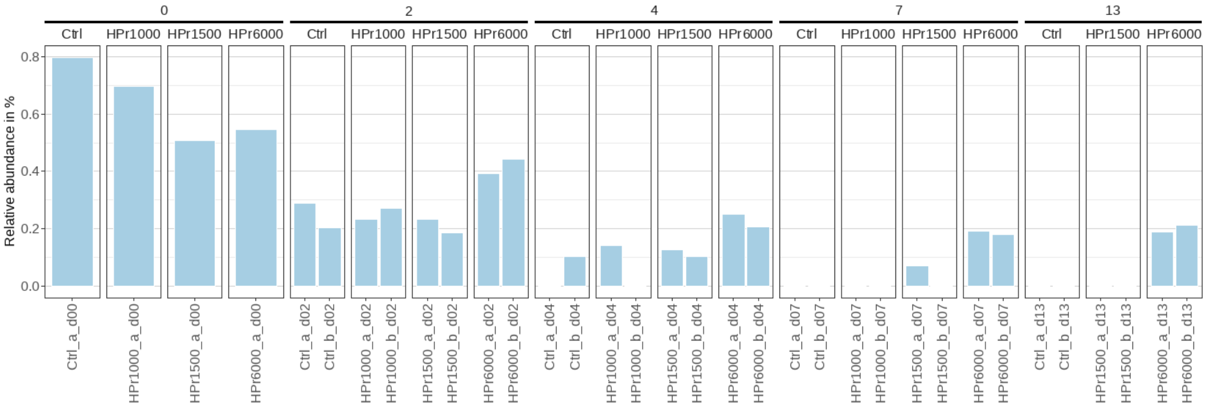


Figure S82. ASV 144. Bacteria, Bacteroidota, Bacteroidia, Flavobacteriales, Weeksellaceae, Cloacibacterium, Multi-affiliation

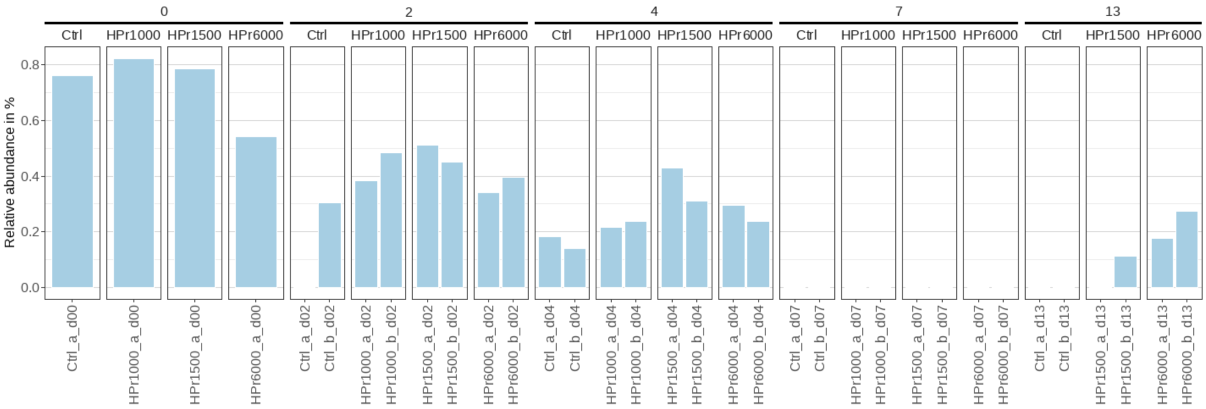


Figure S83. ASV 148. Bacteria, Proteobacteria, Gammaproteobacteria, Burkholderiales, Neisseriaceae, Uruburuella, Multi-affiliation

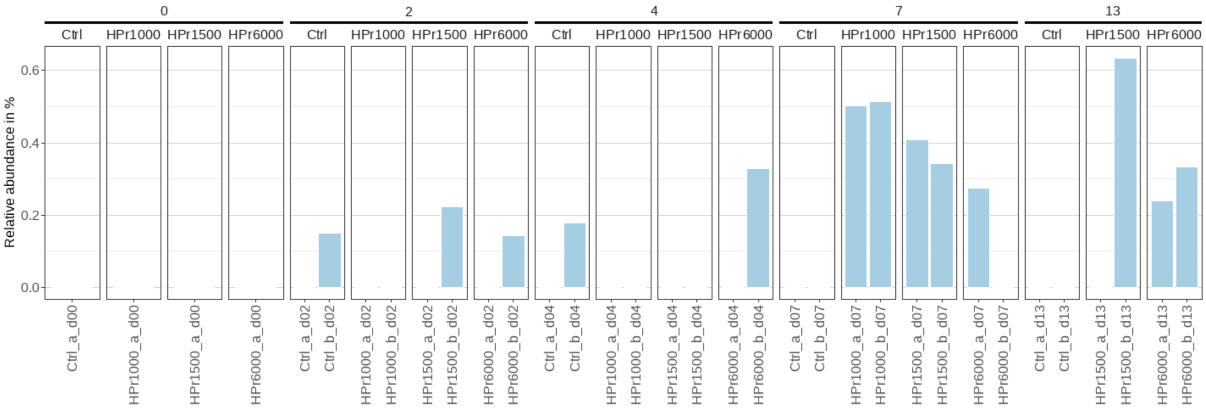


Figure S84. ASV 167. Bacteria, Thermotogota, Thermotogae, Thermotogales, Fervidobacteriaceae, Fervidobacterium, unknown species

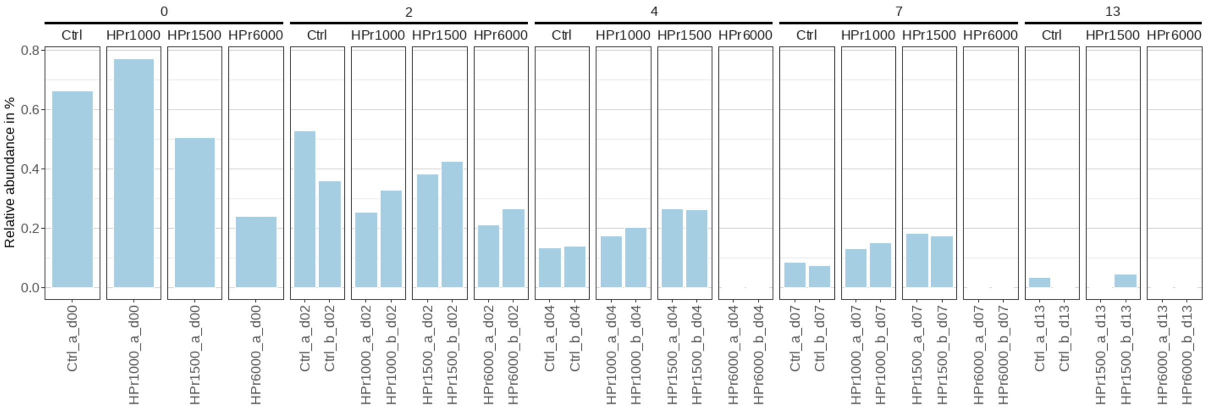


Figure S85. ASV 173. Bacteria, Firmicutes, Clostridia, Peptostreptococcales-Tissierellales, Peptostreptococcaceae, Proteocatella, unknown species

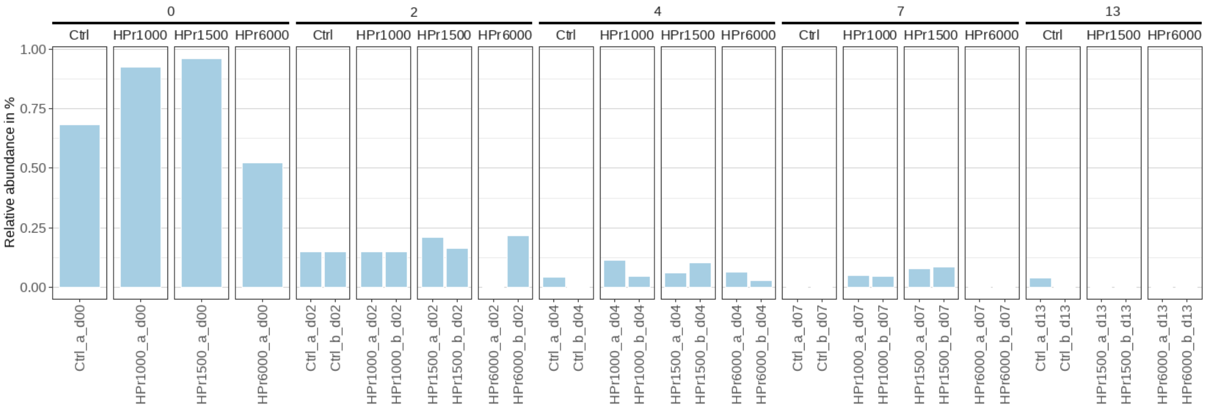


Figure S86. ASV 181. Bacteria, Proteobacteria, Gammaproteobacteria, Aeromonadales, Aeromonadaceae, Aeromonas, Multi-affiliation

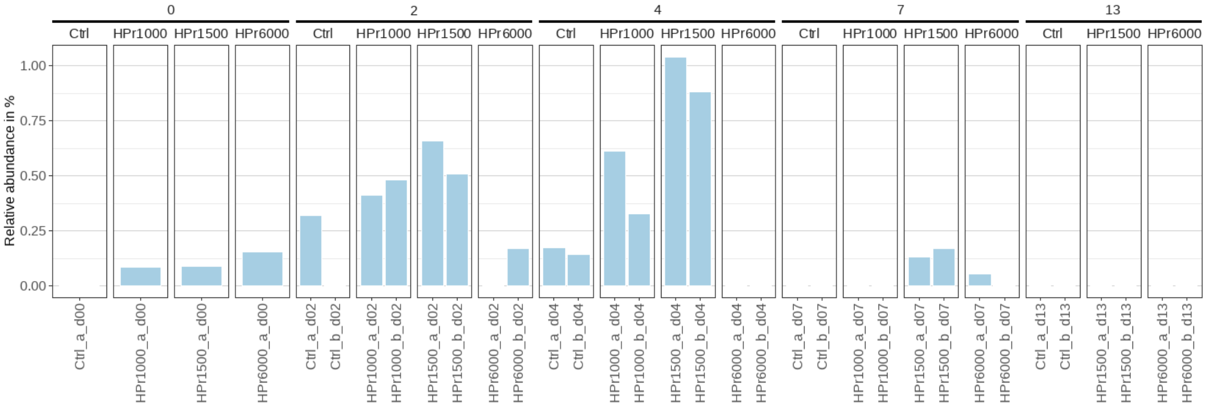


Figure S87. ASV 188. Bacteria, Firmicutes, Clostridia, Oscillospirales, Oscillospiraceae, NK4A214 group, unknown species

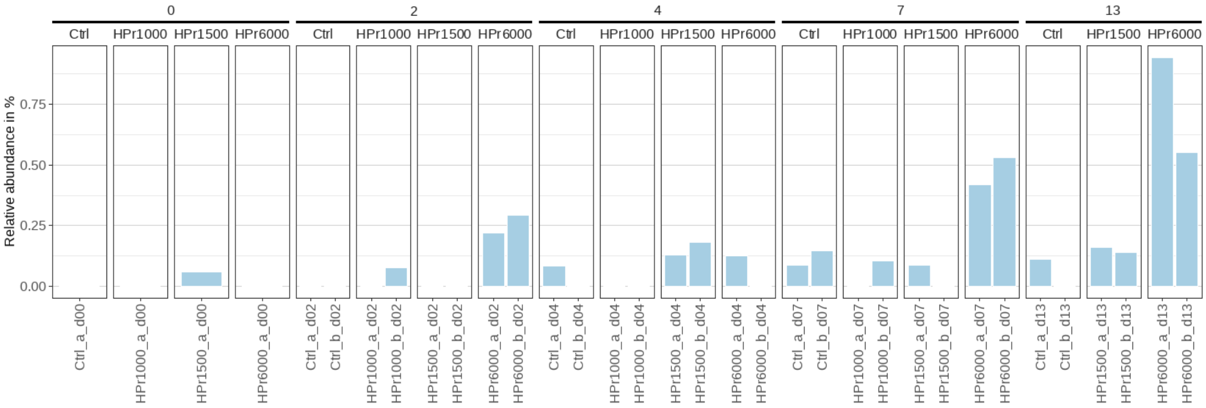


Figure S88. ASV 210. Bacteria, Firmicutes, Clostridia, Clostridiales, Clostridiaceae, Multi-affiliation, Multi-affiliation

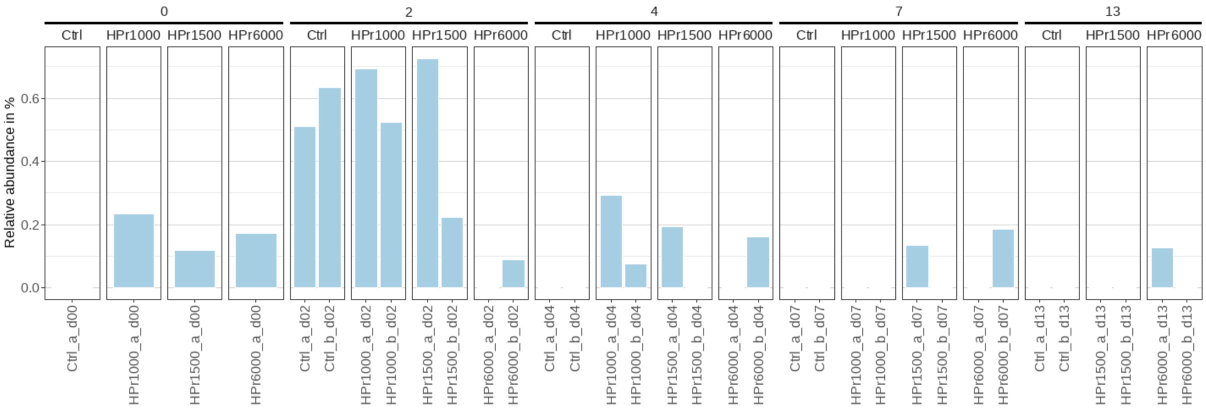


Figure S89. ASV 219. Bacteria, Bacteroidota, Bacteroidia, Bacteroidales, Bacteroidaceae, Bacteroides, Multi-affiliation

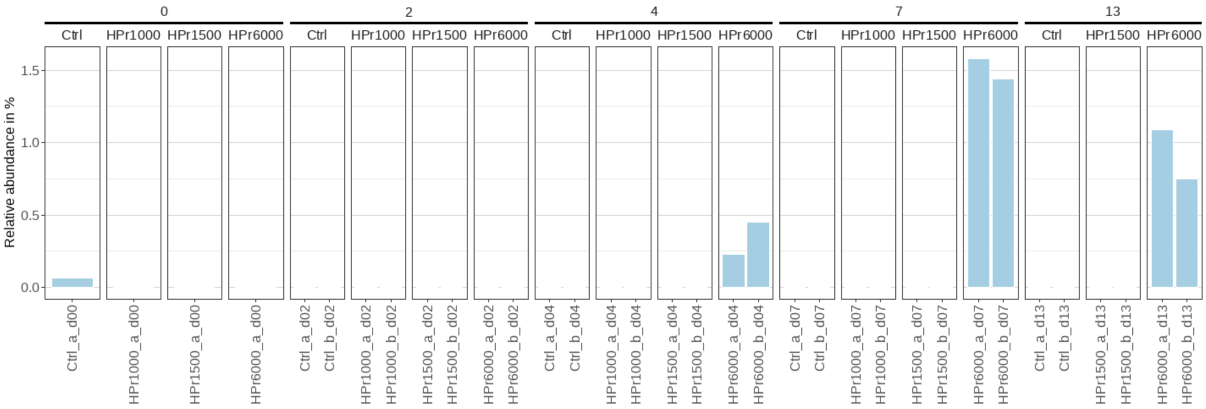


Figure S90. ASV 232. Bacteria, Firmicutes, Negativicutes, Veillonellales-Selenomonadales, Veillonellaceae, unknown genus, unknown species

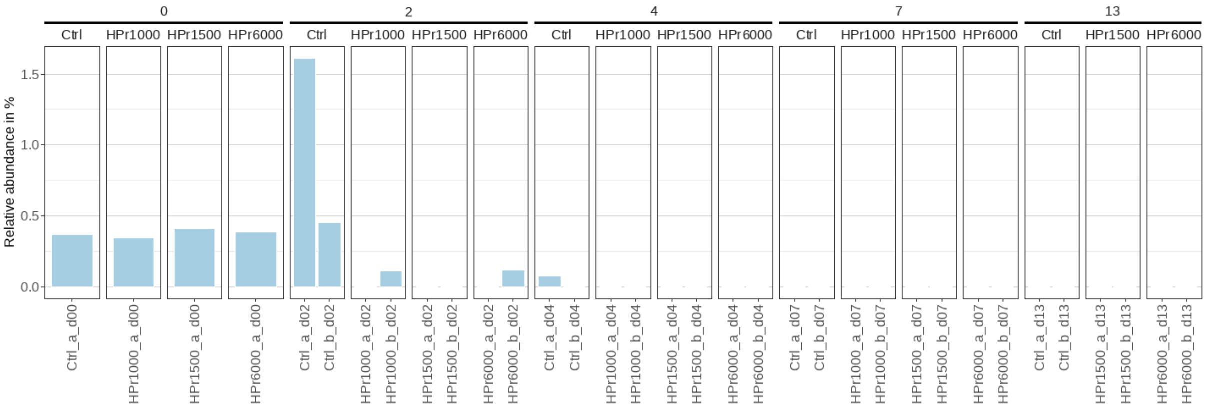


Figure S91. ASV 253. Bacteria, Firmicutes, Clostridia, unknown order, Hungateiclostridiaceae, Pseudobacteroides, unknown species
