## Supplementary File 6 for "Propionic acid-related inhibition during anaerobic digestion: insights into methane production and microbial community adaptation"

### Supplementary Figures S92 to S95

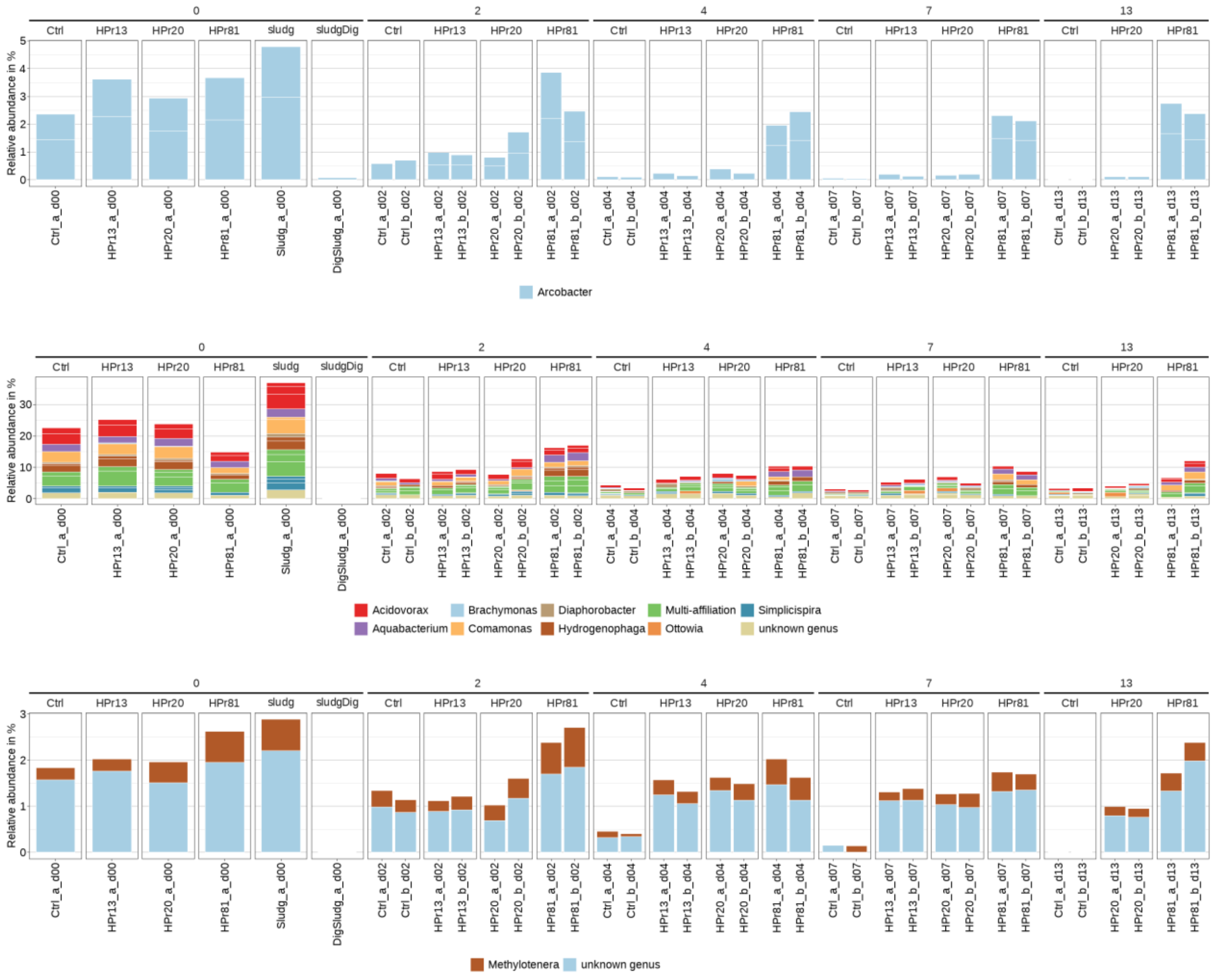


**Figure S92. Relative abundance of the families Arcobacteraceae (top), Comamonadaceae (middle) and Methylophylaceae (bottom) in the first tests, determined by *16S rRNA* gene amplicon sequencing.**

*Sludg* corresponds to the MSS used as substrate, while *sludDig* corresponds to the digested MSS used as inoculum.

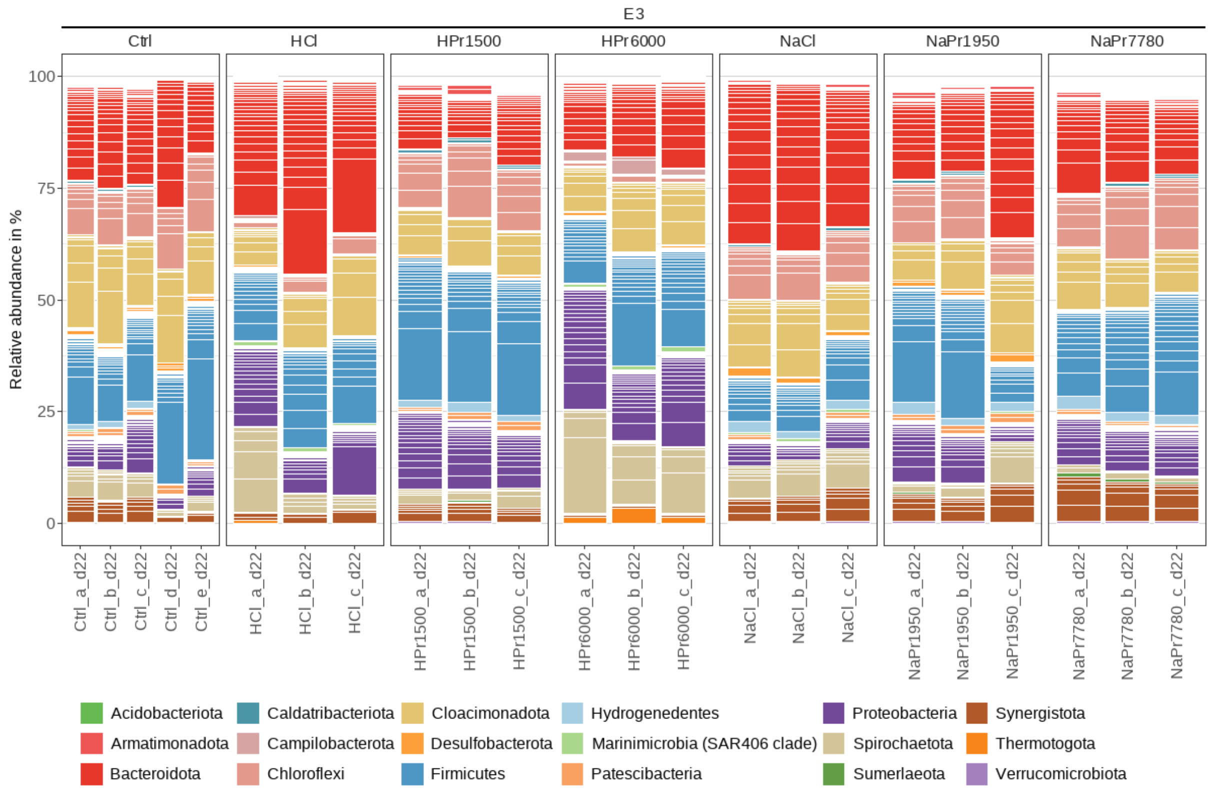


**Figure S93. Composition of the bacterial community in the second tests, at the phylum level, determined by *16S rRNA* gene amplicon sequencing.**

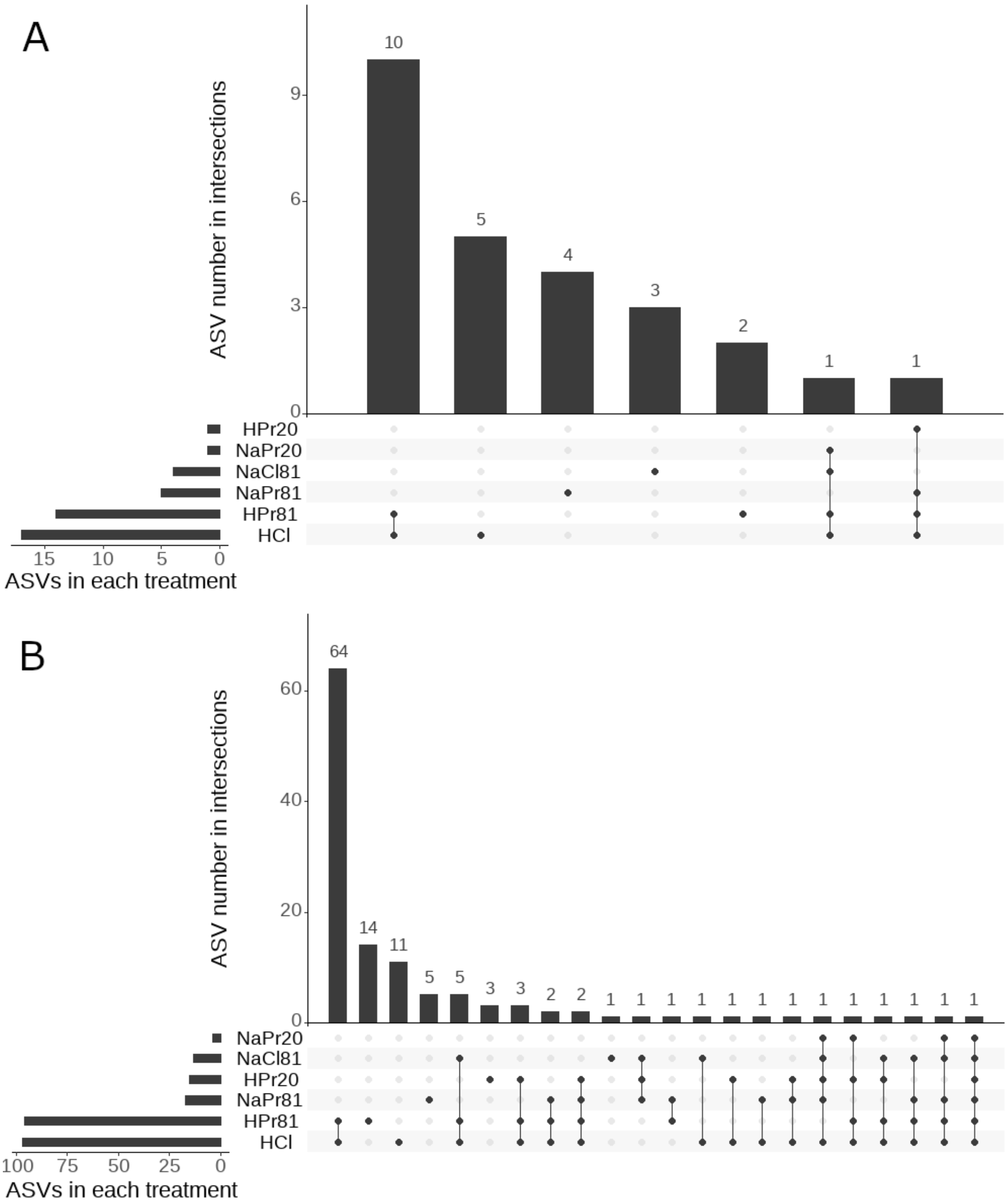


**Figure S94. Upset plot of the ASVs detected as differentially abundant by comparing the different treatments to the control in the second tests.** (A) ASVs significantly more abundant when comparing the treatment to the control Ctrl (total of 26 ASVs). (B) ASVs significantly less abundant when comparing the treatment to the control Ctrl (total of 122 ASVs).

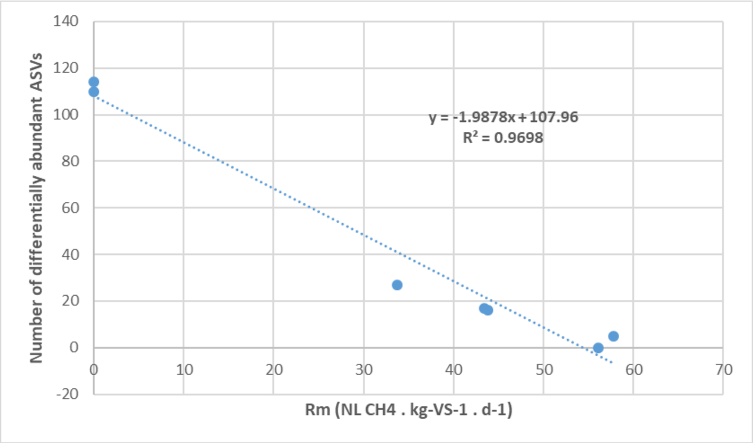


**Figure S95. Linear regression between the maximal methane production rate and the number of differentially abundant ASVs, based on the first tests. The maximal production rates were estimated using the Gompertz model.**
