## Supplementary File 1 for "Propionic acid-related inhibition during anaerobic digestion: insights into methane production and microbial community adaptation"

### Supplementary tables

#### Table S1. Characteristics of inoculum and MSS[a] utilized for the various assays

| | Substrate: fresh MSS | Inoculum: digested MSS |
|---|---|---|
| Total solids – TS (%) | 3.3 – 4.7 | 2.3 – 2.6 |
| Volatile solids – VS (%TS) | 71 – 77 | 61 – 63 |
| pH (–) | 6.0 – 6.3 | 7.4 – 7.6 |

[a]MSS: municipal sewage sludge
