## Supplementary File 2 for "Propionic acid-related inhibition during anaerobic digestion: insights into methane production and microbial community adaptation"

Supplementary Figures S1 to S5

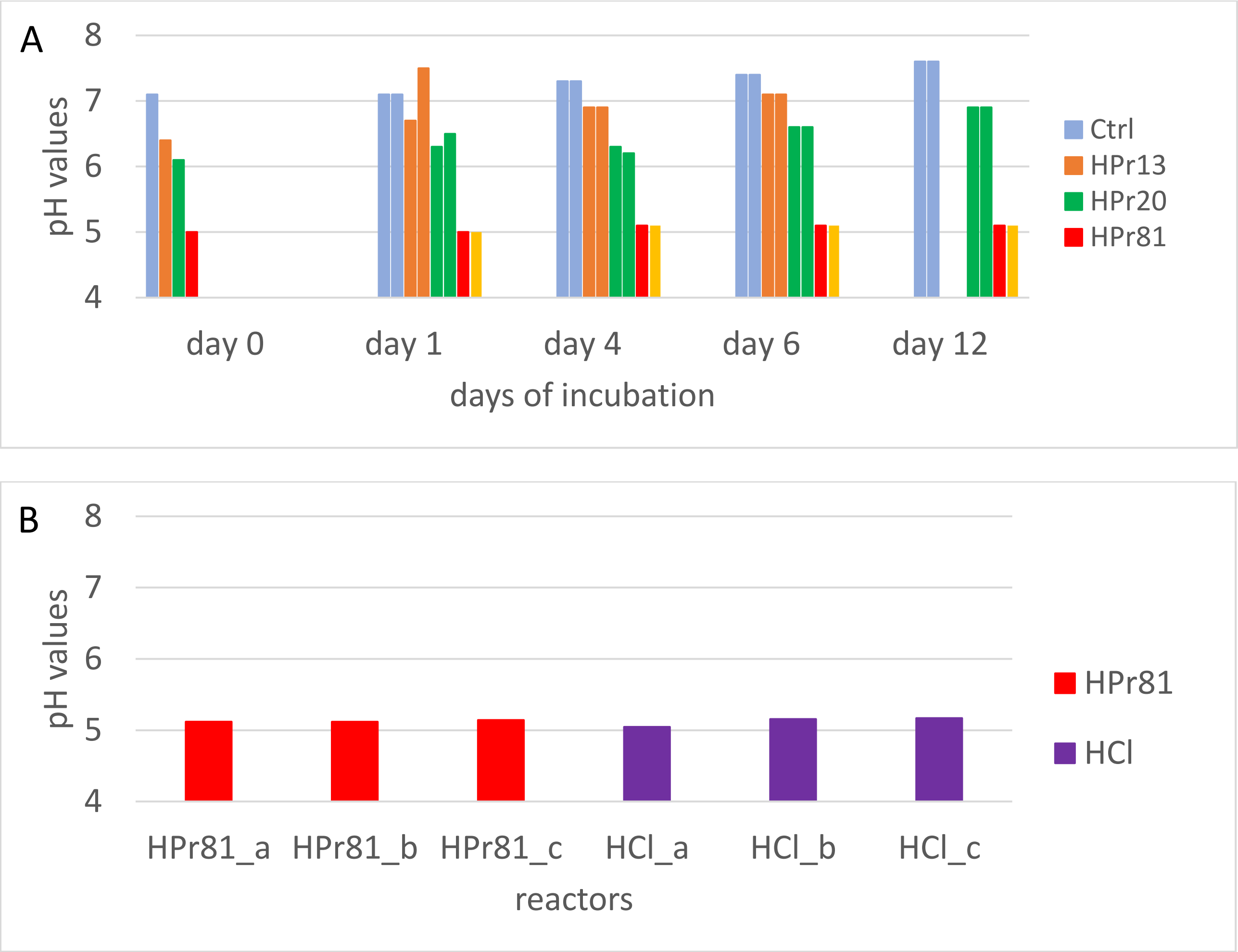


**Figure S1. pH values measured during the first and second tests. A. First tests over time. B. Second tests at day 0.**

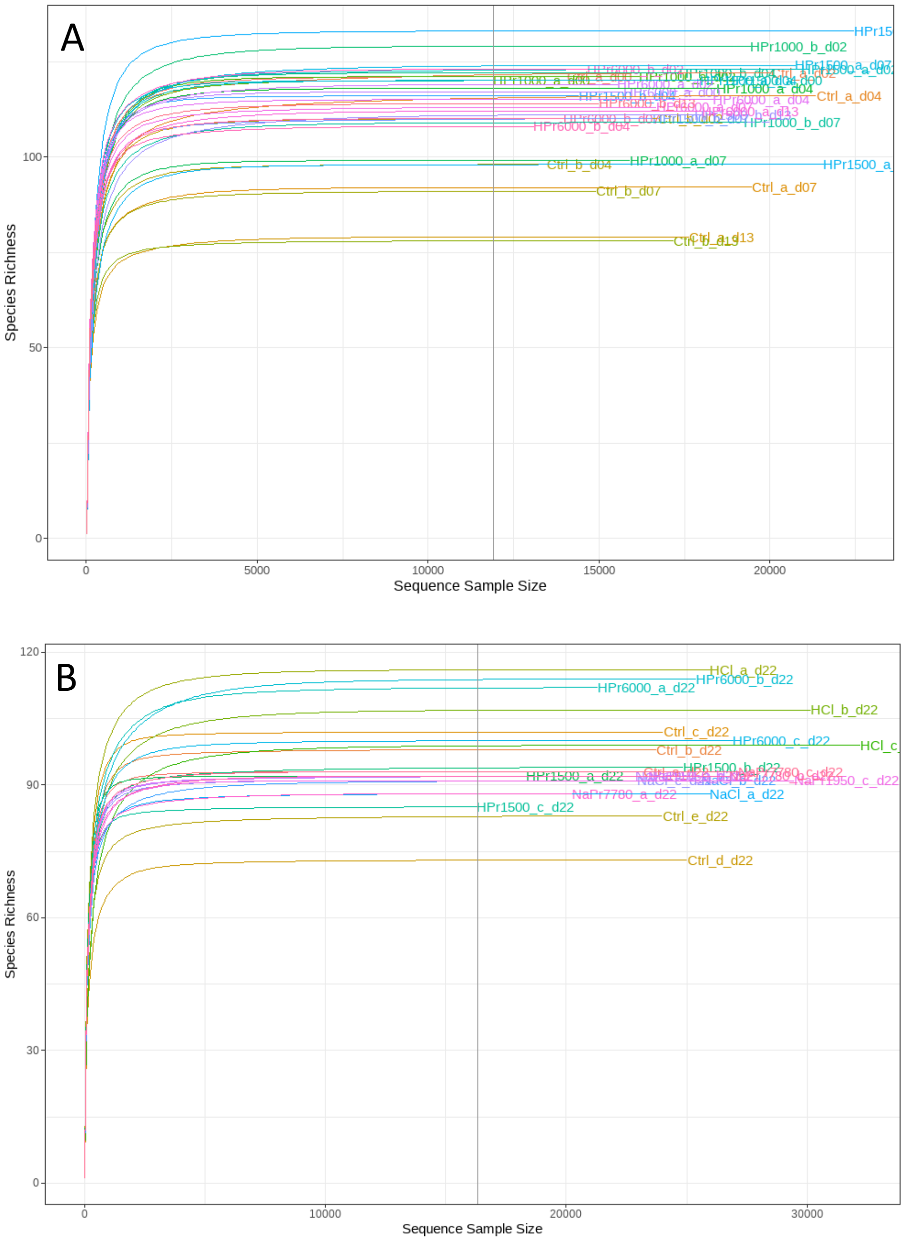


**Figure S2. Rarefaction curves at the ASV level, calculated for each sample analyzed by 16S rRNA gene amplicon sequencing, after read filtering. A. First tests. B. Second tests.**

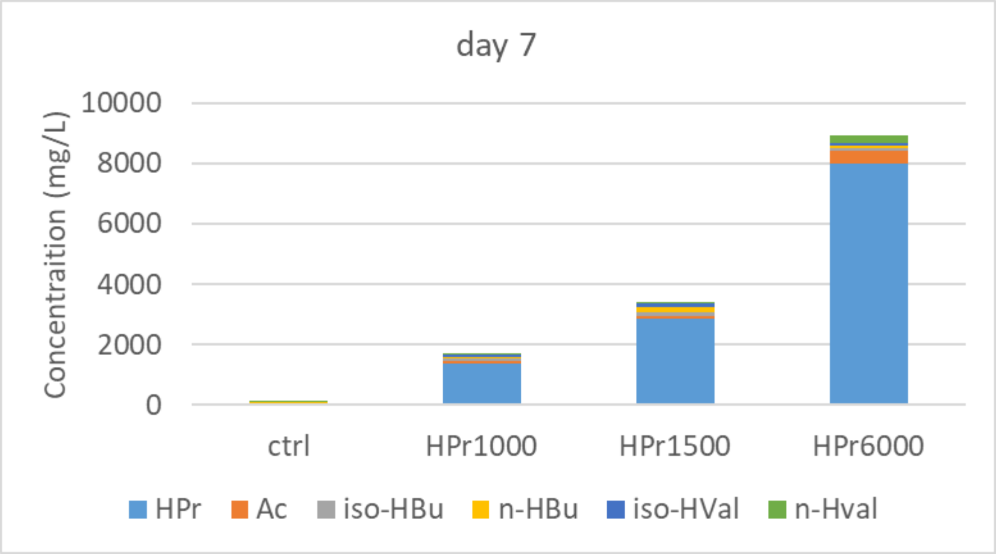


**Figure S3. Volatile fatty acid concentrations measured during the first tests on day 7.**
For each condition and for each volatile fatty acid, the average concentration among the different reactors is shown.

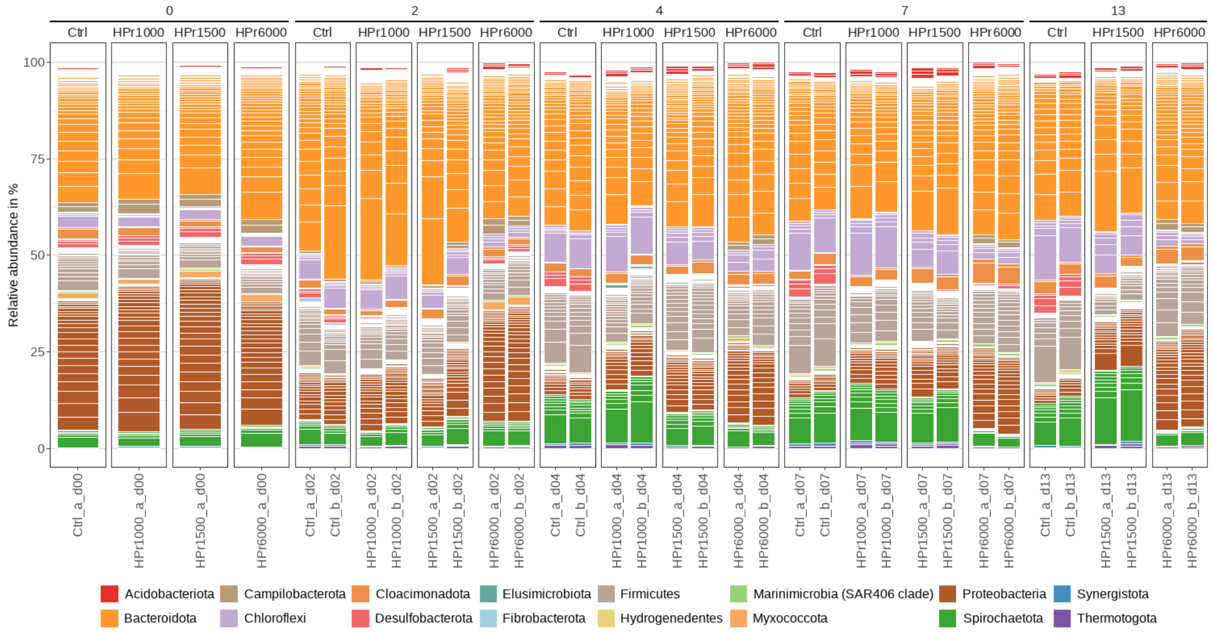


**Figure S4. Composition of the bacterial community in the first tests, at the phylum level, determined by 16S rRNA gene amplicon sequencing.**

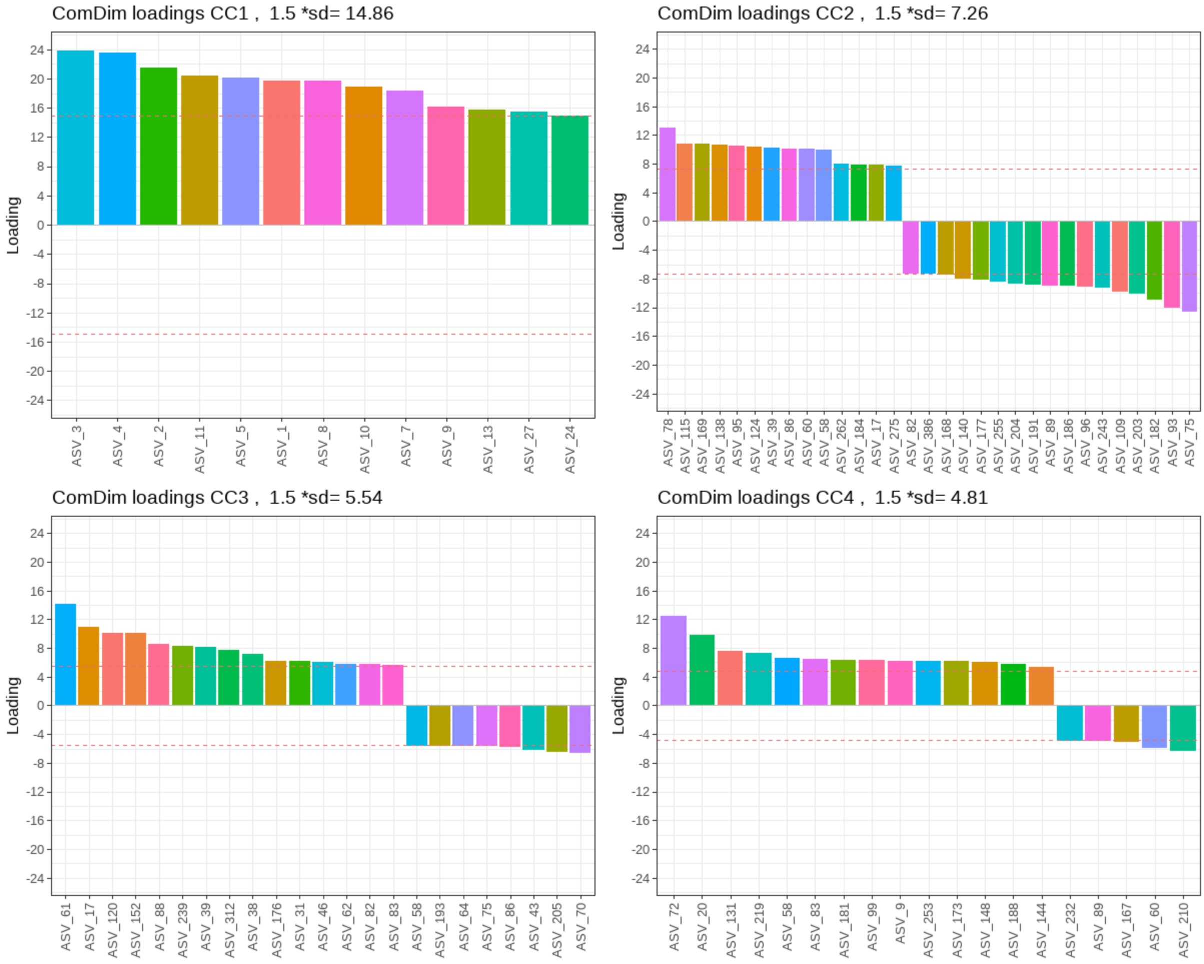


**Figure S5. Loading of the ASVs selected during the Common Component and Specific Weight Analysis based on microbial community composition of the first assay.**

ASVs were selected based on their loadings, which exceeded 1.5 times the standard deviation of all ASV loadings on the respective CC. The red dotted line indicates the selection threshold.
