## Supplementary File 3 for "Propionic acid-related inhibition during anaerobic digestion: insights into methane production and microbial community adaptation"

### Supplementary information

#### Analytical methods

The concentrations of the different VFAs present in the samples before digestion and at reactor opening were measured by gas chromatography coupled with a flame ionization detector (GC-FID). For dry-matter determination, oven drying was used (NF EN 17815-934). Volatile-matter content was then obtained by ignition in a furnace (NF EN 15169). Alkalinity (TAC) was measured by pH-metric titration (NF EN ISO 9963-1). The pH was measured using a pH meter (Mettler Toledo, Switzerland).
